# Engineering an Escherichia coli strain for production of long single-stranded DNA

**DOI:** 10.1101/2024.02.27.582394

**Authors:** Konlin Shen, Jake J. Flood, Zhihuizi Zhang, Alvin Ha, Brian R. Shy, John E. Dueber, Shawn M. Douglas

## Abstract

Long single-stranded DNA (ssDNA) is a versatile molecular reagent with applications including RNA-guided genome engineering and DNA nanotechnology, yet its production is typically resource-intensive. We introduce a novel method utilizing an engineered *E. coli “helper”* strain and phagemid system that simplifies long ssDNA generation to a straightforward transformation and purification procedure. Our method obviates the need for helper plasmids and their associated contamination by integrating M13mp18 genes directly into the *E. coli* chromosome. We achieved ssDNA lengths ranging from 504 to 20,724 nucleotides with titers up to 250 μg/L following alkaline-lysis purification. The efficacy of our system was confirmed through its application in primary T cell genome modifications and DNA origami folding. The reliability, scalability, and ease of our approach promises to unlock new experimental applications requiring large quantities of long ssDNA.

## Introduction

Single stranded DNA (ssDNA) plays a crucial role in biotechnology, particularly in DNA nanotechnology and gene editing^1,2^. Synthesis of long ssDNA exceeding 5000 nucleotides (nt) is challenging, and significant barriers prevent scalable production. Direct chemical synthesis via phosphoramidite chemistry is limited to lengths of 300–400 nt due to incorporation errors and depurination^3^. To obtain longer ssDNA strands, current practices employ double-stranded DNA (dsDNA) as a template. For instance, asymmetric PCR can generate ssDNA up to 15,000 nt in length^4^. Other approaches include PCR amplification using differentially modified primers: phosphorylated and unphosphorylated for lambda exonuclease digestion^5^, or biotinylated and non-biotinylated for streptavidin-bead separation^6^—resulting in isolation of long ssDNA strands. However, these techniques typically yield less than 1 microgram (μg) of ssDNA per 50-microliter (μL) reaction, making the production of milligram quantities costly and inefficient due to the extensive labor and high reagent consumption, underscoring the necessity for more scalable and economical ssDNA production methods.

Bioproduction of ssDNA in *E. coli* offers a scalable alternative. By engineering the M13 bacteriophage to include specific ssDNA sequences, up to 590 mg ssDNA per liter of culture can be recovered once the viral particles are processed^7^. However, the resulting ssDNA products include about 6 kilobases of the M13 genome, and issues with sequence stability and titer can arise for custom insertions longer than 2 kilobases^8,9^. Additionally, utilizing phages carries a contamination risk for other bacterial cultures.

Improvements to M13-based ssDNA production can be made through use of helper phages^10^ or helper plasmids^11,12^. In this approach, the phage genes are encoded on the helper entity and the desired sequence is cloned into a phagemid, a separate plasmid bearing an M13 replication origin and packaging sequence. This separation prevents the inclusion of the M13 genome in the final ssDNA product. Phagemid utilization can yield ssDNA lengths up to 51,000 nt^13^ and, with optimized *E. coli* growth conditions, titers can reach up to 80 mg/L of culture^14^. Nevertheless, the consistency of ssDNA production using helper elements is variable, with outcomes ranging from off-target by-products to a lack of ssDNA^15^.

Here, we describe our novel phagemid-based method that improves the quality of bacteriologically produced ssDNA (Fig. 1). Our method relies on an engineered “helper strain” called “eScaf”. We posited that inconsistent target ssDNA yields may result from high or variable expression of M13 genes which, in turn, may arise from fluctuations in helper plasmid copy number. To address this, we integrated the M13 genes directly into the bacterial chromosome. We assessed the performance of eScaf, considering the chromosomal integration site, and target ssDNA length and composition. We demonstrate that eScaf consistently produces high-quality ssDNA at concentrations of 100–200 μg per liter of bacterial culture using shake flasks. We further validated our method by successfully employing eScaf-generated ssDNA in DNA origami fabrication and in CRISPR-mediated gene editing applications, highlighting its utility in advanced biotechnological applications.

**Fig. 1.**
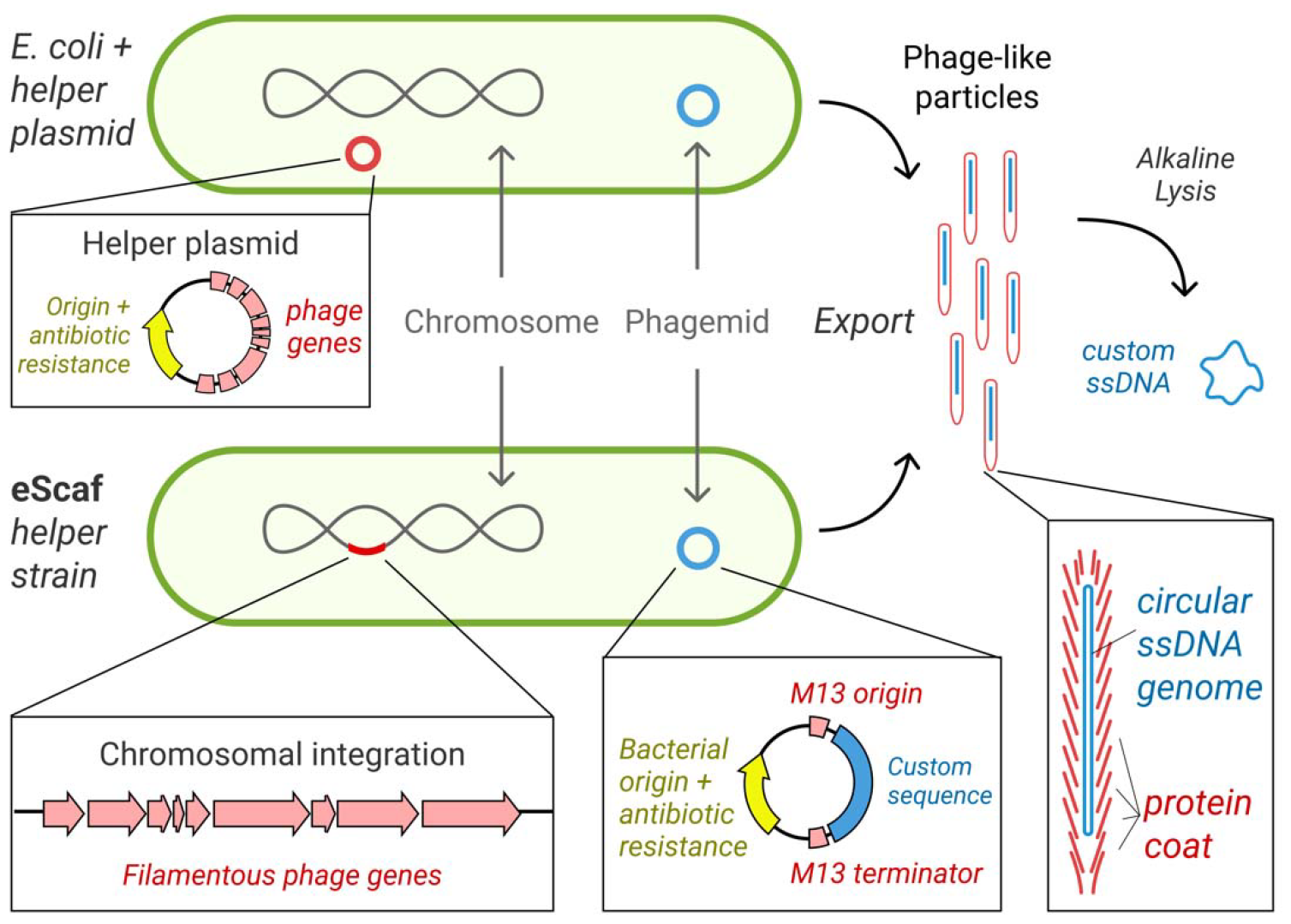
Overview of ssDNA production using the eScaf helper strain. Conventional phagemid-based ssDNA synthesis involves co-transforming an *E. coli* host strain with a helper plasmid and a phagemid, which leads to the production of phage-like particles harboring the desired ssDNA (top). The helper plasmid mediates the expression of the phage genes necessary for this process. In contrast, our eScaf method (bottom) streamlines this by integrating the phage genes directly into the chromosome, establishing a ‘helper strain’ that requires only the phagemid for transformation. For both strategies, phage-like particles containing ssDNA are released into the culture medium and can be isolated through PEG fractionation. The target ssDNA is subsequently extracted from these particles using alkaline lysis followed by ethanol precipitation for purification.

## Methods

### Helper strain construction

We integrated the M13mp18 and M13KO7 genomes into the chromosome of the TG1 *E. coli* strain using a CRISPR RNA-guided transposase system^28^. The M13mp18 and M13KO7 genomes were cloned into “donor” plasmids carrying mini-transposons. gRNA targeting locations within the genome were cloned into a separate “effector” plasmid also carrying the transposition and CRISPR machinery. Transposition was achieved by dual transformation of the donor and effector plasmids into TG1 and growth at 30 °C. Colonies were then screened for successful integration using cPCR. Colonies carrying the insert were then cured of their plasmids by overnight growth in LB without selection and subsequently plated onto regular LB agar plates without selection to use as a master plate. Replica plating was used to transfer those colonies onto plates selecting for either the donor plasmid or the effector plasmid. Colonies which did not grow on either selection plate were then picked from the master plate and integration of M13 genes was once again verified through cPCR. Glycerol stocks and chemically competent cells were made from colonies showing successful integration and curing for banking and ssDNA production, respectively.

### Phagemid construction

Phagemids were created by cloning inserts into the pScaf^11^ backbone using Gibson assembly or golden gate cloning. Phagemids designed for testing sequence length between 504-nt and 10,080-nt were constructed from 1 to 10 gBlocks (IDT), 1008 bp in length using Gibson assembly. To construct the 20,874-nt phagemid, a 9 kb fragment from the dynein gene and an RFP expression operon were cloned into the 10,080-nt phagemid using Gibson assembly. The phagemid for the DNA origami tile was constructed using golden gate cloning to insert a 500 bp gBlock into the phagemid backbone. The phagemid designed for the BCMA-CAR CRISPR knock-in was constructed via PCR by amplifying the template from plasmids bearing the construct and cloning them into the pScaf backbone using Gibson assembly.

### Amplification and purification of ssDNA from helper strains

We first transformed the strains with a phagemid of interest, and then picked a colony and subsequent growth in liquid culture with antibiotic selection. Cultures were grown in standard shaker flasks at 30 °C for 12∼16 hours. To harvest ssDNA, cultures were first chilled on ice for 35’, then bacteria and phage-like particles were separated by pelleting the bacteria using centrifugation for 15’ at 7 kRCF, 4 °C. The supernatant was collected and the phage-like particles were purified out by PEG precipitation. PEG-8k (40 g/L) and NaCl (30 g/L) were added to the mixture and incubated on ice for 30’. Then the phage-like particles were pelleted by spinning the mixture at 9 kRCF for 15’ at 4 °C, discarding the supernatant, and resuspending the pellet in TE. To prevent possible contamination of the ssDNA products by bacterial DNA, any residual bacteria in the phage-like particle solutions were pelleted by spinning the phage-like particle solutions at 15 kRCF and 4 °C for 15’, transferring the supernatant to clean tubes, and spinning the solution again at 15 kRCF, 4 °C for 15’ to ensure no bacteria remained in the solution. The phage coat proteins were subsequently stripped via alkaline lysis. 2x volumes of P2 buffer (Qiagen) was added to the phage solution for lysis and 1.5x volumes of P3 buffer (Qiagen) was immediately added to neutralize the reaction. The mixture was then incubated on ice for 15’ and the phage debris was pelleted by centrifugation at 15 kRCF, 4 °C for 15’. The supernatant of the mixture was then moved to a clean tube and 3x volumes of cold 100% EtOH was added to the mixture and the ssDNA precipitated at -20 °C overnight. Finally, the ssDNA was pelleted by centrifugation at 16 kRCF, 4 °C at 15’. The supernatant was aspirated off and the pellet washed in cold 70% EtOH by a second centrifugation step at 15 kRCF, 4 °C for 15’. The supernatant was aspirated off again, and the pellet was dried at room temperature by leaving the tube inverted on the bench. Finally, the pellet was resuspended in TE buffer. Concentrations of ssDNA were measured using a NP80 nanophotometer (IMPLEN) on the ssDNA setting and purity was evaluated using agarose gel electrophoresis (2% TBE gel with 11 mM Mg added).

### Fabrication of DNA origami tile

We designed a small DNA origami tile using Cadnano^29^ and optimized the staple routing path using a custom software developed by the Douglas lab (manuscript in prep.). A custom 3024-nt origami scaffold was produced using the methods described above for phagemid cloning and scaffold purification. Staples were ordered from IDT. The DNA origami tile was folded by mixing the custom scaffold with a 10x molar excess of staples and subjecting the mixture to the following thermal ramp:

1. Incubate at 65 °C for 15’.
2. Incubate at 60 °C for 1 hour, decreasing 1°C per cycle.
3. Go to Step 2 and repeat an additional 40 times.

DNA origami nanostructures were purified via PEG precipitation by adding an equivalent volume of a 15% PEG-8000 solution (15% w/v PEG-8000, 5 mM Tris, 1 mM EDTA, 5 mM NaCl, 20 mM MgCl2) and centrifuging for 25’ at 16 kRCF. The supernatant was aspirated, and the pellet was resuspended in 1xFOB8 (5 mM Tris, 1 mM EDTA, 8 mM MgCl2) and allowed to resuspend for 30 minutes at room temperature. The PEG precipitation step was repeated one more time.

DNA origami tiles were evaluated using gel agarose electrophoresis (2% TBE with 11 mM Mg) and negative-stain transmission electron microscopy. Briefly, formvar-coated copper grids (Ted Pella) were plasma treated with a Pelco glow discharge unit (30s hold/30s glow discharge). 1 uL of DNA origami at a concentration of 6.55 nM was incubated on the grid for 3 minutes, then the excess sample was wicked away using filter paper. Grids were washed twice in 1xFOB8 and washed once in freshly prepared 2% uranyl formate. A droplet of 2% uranyl formate was then placed on the grid and incubated for 1 minute at room temperature for staining, then wicked away and the grid was dried by a vacuum aspirator. Micrographs of the negatively stained grids were collected on a Tecnai T12 (FEI). 2D class averages were made using EMAN2.

### BCMA-CAR knock-in T cells using ssDNA products generated by helper strains

Cas9 ribonucleoproteins (RNPs) targeting the TRAC locus were produced by complexing gRNA to Cas9 with the addition of an ssODN electroporation enhancer. To create the gRNA, synthetic CRISPR RNA (crRNA) and *trans*-activating crRNA (tracrRNA) were chemically synthesized individually (Dharmacon) and each strand was resuspended in IDT duplex buffer at a concentration of 160 μM. The two strands were mixed 1:1 v/v and annealed by incubation at 37 °C for 30’ to form an 80 μM gRNA solution. ssODN enhancer was mixed into the gRNA solution at a 0.8:1 volume ratio followed by 40 μM Cas9-NLS (Berkeley QB3 MacroLab) at a 1:1 v/v. The entire mixture was incubated for 15-30’.

Primary adult blood cells were obtained from healthy human donors as a leukapheresis pack purchased from StemCell Technologies, Inc. Primary human CD3+ T cells were isolated by negative selection using magnetic cell isolation kits (StemCell, catalog no. 17951). The isolated cells were then activated with magnetic anti-human CD3/CD28 dynabeads (CTS, ThermoFisher). Beads were mixed at a 1:1 ratio with cells and cells were cultured at a density of 1 x 10^6^ cells/mL in complete X-Vivo 15 culture media (Lonza, 5% fetal bovine serum, 10 nM N-acetyl-L-cysteine, 50 μM 2-mercaptoethanol) supplemented with 5 ng/mL of IL-7 and IL-15 for 48 hours.

Activated primary T cells were debeaded 48 hours post-activation using an EasySep magnet (StemCell) and electroporated using the Lonza 4D 96-well electroporation system. Immediately before electroporation, cells were centrifuged at 90 x g for 10’ then resuspended at 0.5 x 10^6^ cells per 20 μL of Lonza P3 electroporation buffer. cssDNA encoding the BCMA-CAR and RNPs were mixed and incubated for at least 5’, then combined with cells and transferred to the Lonza 96-well electroporation shuttle. 50 pmol of RNP was used for each electroporation. Primary T cells were electroporated using pulse code EH-115, immediately rescued with prewarmed culture media, and incubated at 37°C for at least 15’. Cells were then transferred to 96-well plates and maintained at 1.0 x 10^6^ cells/mL in complete X-Vivo 15 culture media. Fresh cytokines and media were added every 2-3 days and split using identical dilution factors for all samples to prevent cell overgrowth and maintain comparable cell counts.

Knock-in percentage and cell counts were measured on day 10 of culturing (day 8 post-electroporation) by flow cytometry using the Attune NxT flow cytometer with a 96-well autosampler (ThermoFisher Scientific). Briefly, cells were resuspended in EasySep Buffer (PBS, 2% fetal bovine serum, 1mM EDTA), then incubated with biotintlyated recombinant BCMA protein (Acro Biosystems) for 20’ at 4 °C. Cells were washed and stained with GhostDye 780 (Tonbo), TCRab-BV421 (BioLegend), and Streptavidin-APC (BioLegend) for 20’ at 4 °C. Events were recorded from a fixed volume for all samples to obtain comparable live cell counts between conditions.

## Results

### eScaf successfully generated custom ssDNA from 504-nt to 20,724-nt in length

We constructed two eScaf helper strains by inserting the genomes of M13mp18 and M13KO7 into the chromosome of the JM101 E. coli derivative, TG1. These strains were transformed with phagemids carrying ssDNA sequences varying from 504 to 20,724 nucleotides. The transformed strains were cultured in LB media, after which we isolated phage-like particles from the culture supernatant and extracted the ssDNA by alkaline lysis. While both the M13mp18 and M13KO7 strains successfully produced ssDNA across a range of lengths, the M13mp18 strain achieved notably higher ssDNA yields (Fig. 2a). For a 3396-nt ssDNA strand, M13mp18 helper strain produced average titers of 206 μg/L whereas M13KO7 strains produced average titers of 11 μg/L (Fig. 2b). While both integrated strains showed target band purities above 90%, the M13mp18 strain produced fewer off-target species than the M13KO7 strain (Fig. 2c). Both strains showed limited efficiency in producing the 504-nt ssDNA (Fig. S2), consistent with previous reports that short ssDNA is difficult to produce using phagemid approaches^12^. Below we summarize specific factors that affected titer and purity of ssDNA using our new method.

**Fig. 2.**
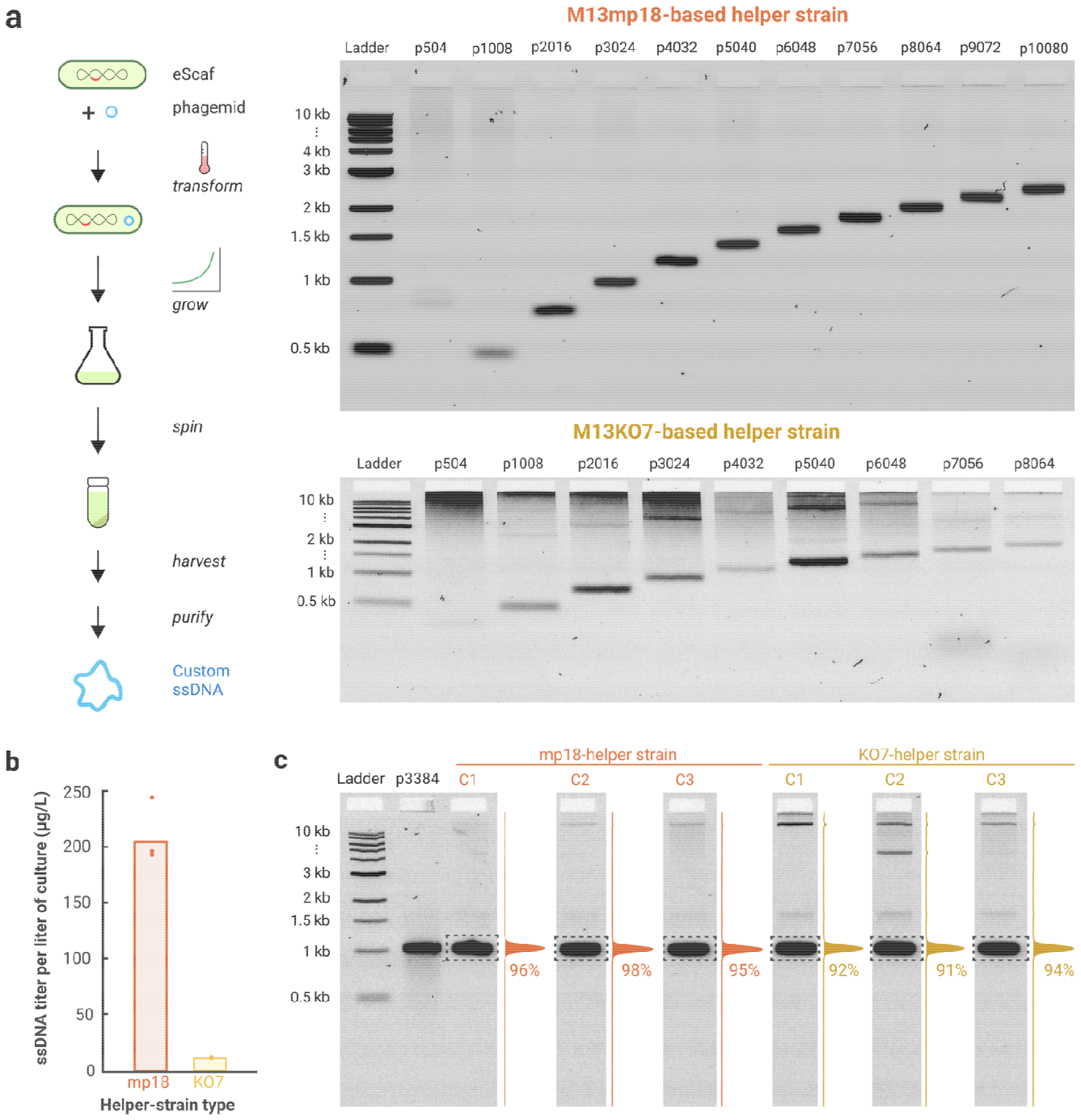
Comparison of ssDNA production by M13mp18 and M13KO7 helper strains. **a)** Both M13mp18 and M13KO7-based helper strains can produce ssDNA within a size range of 504 to 20,724 nucleotides. The M13KO7-based strains exhibit a higher presence of off-target species and background than the M13mp18-based strains. **b)** The total ssDNA yield is greater when using M13mp18-based helper strains compared to M13KO7-based strains. **c)** While ssDNA purity exceeds 90% for both helper strains, M13mp18 strains produce ssDNA that is marginally purer than that from M13KO7 strains. Purity percentages were determined by quantifying the intensity of the target band within the dotted box and normalizing this value against the total intensity of the full lane. All ladders are 1 kb ladders from New England Biolabs.

### Phagemid-helper strain compatibility affects ssDNA yield

Contrary to earlier reports suggesting superior ssDNA purity from M13KO7 plasmid systems^12^, our findings indicate a decrease in purity when employing M13KO7 helper strains. We hypothesized that this discrepancy could be linked to the compatibility between the M13 packaging sequences in the phagemids and the helper strains. Our phagemids, based on the pScaf backbone^11^, incorporate a modified packaging sequence, which we suspected might influence the final ssDNA product’s quality. To test this hypothesis, we conducted experiments with our M13KO7 and M13mp18 helper strains using two distinct phagemids: the pScaf-derived phagemid and another that was previously paired with the M13KO7 helper plasmid^12^ (referred to as the “KO7 phagemid”).

Our data revealed a clear dependency of ssDNA yield on the phagemid used (Fig. 3a). Optimal yields were achieved when pairing the M13mp18-helper strain with the pScaf phagemid and the M13KO7- helper strain with the KO7 phagemid. In contrast, using the KO7 phagemid with the M13mp18-helper strain or the pScaf phagemid with the M13KO7-helper strain resulted in significantly lower ssDNA production. These results indicate that both M13KO7 and M13mp18 helper strains can be effective when paired with the appropriate phagemid. Given the slight edge in ssDNA purity and our primary use of the pScaf backbone in previous phagemid cloning, we performed all subsequent work with the M13mp18-helper strain.

**Fig. 3.**
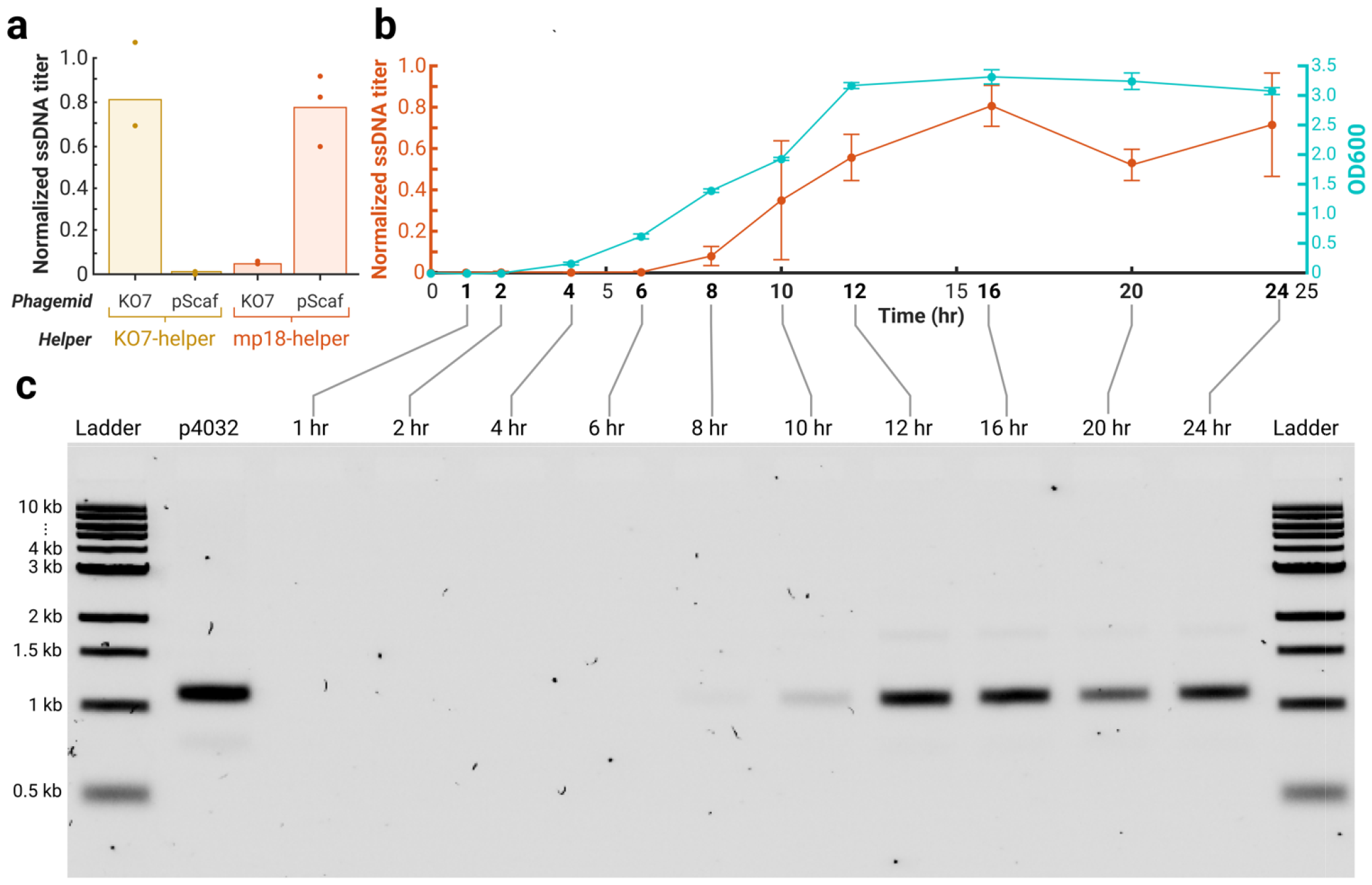
Single-stranded DNA production requires phagemid-host strain compatibility and is time dependent. **a)** Optimal titer results are achieved when both the helper strain and phagemid are suitably matched. M13KO7-integrated helper strains fail to package ssDNA derived from the pScaf phagemid but work well with the KO7 phagemid. Conversely, M13mp18-integrated helper strains show high compatibility pScaf-based phagemids, are less effective with the KO7 phagemid. **b)** Time series of ssDNA production for a 4032-nt ssDNA strand in the M13mp18 helper strain. Measurable ssDNA production begins 8 hours after inoculation and ceases following 16 hours of growth, with the culture reaching saturation at approximately 12 hours. **c)** Agarose-gel analysis of temporal ssDNA production by the M13mp18 helper strain shows a predominant desired on-target ssDNA bands with a minor presence of an off-target band. Ladders are 1 kb ladders from New England Biolabs.

### Production of ssDNA peaked after 16 hours of growth at 30 °C

We observed that ssDNA production reached its maximum after 16 hours of growth at 30 °C, which occurs roughly 4 hours after culture saturation (Fig. 3b). Notably, ssDNA titer exhibited a reproducible decline following its peak, followed by a marginal increase later in the growth cycle. The final increase did not appear to be due to cellular lysis, which would be indicated by additional gel bands typically associated with extraneous nucleic acid contamination (Fig. 3c).

### Integration location affects ssDNA titer, but does not affect purity

We found that integrating the M13mp18 genome in different genomic loci of TG1 affected ssDNA titer, but not its purity (Fig. 4a, b). We tested five different insertion sites, both intergenic and intragenic (Table 1), but did not find that insertion into intergenic regions or intragenic correlated with ssDNA titer. In fact, we found the location with the highest titer was within the *Ula*D gene (155 μg/L). On the other hand, insertion into an intergenic region close to the origin of replication resulted in no ssDNA production. In all integration sites, ssDNA purity was above 90% (Fig. S3).

**Table 1.**
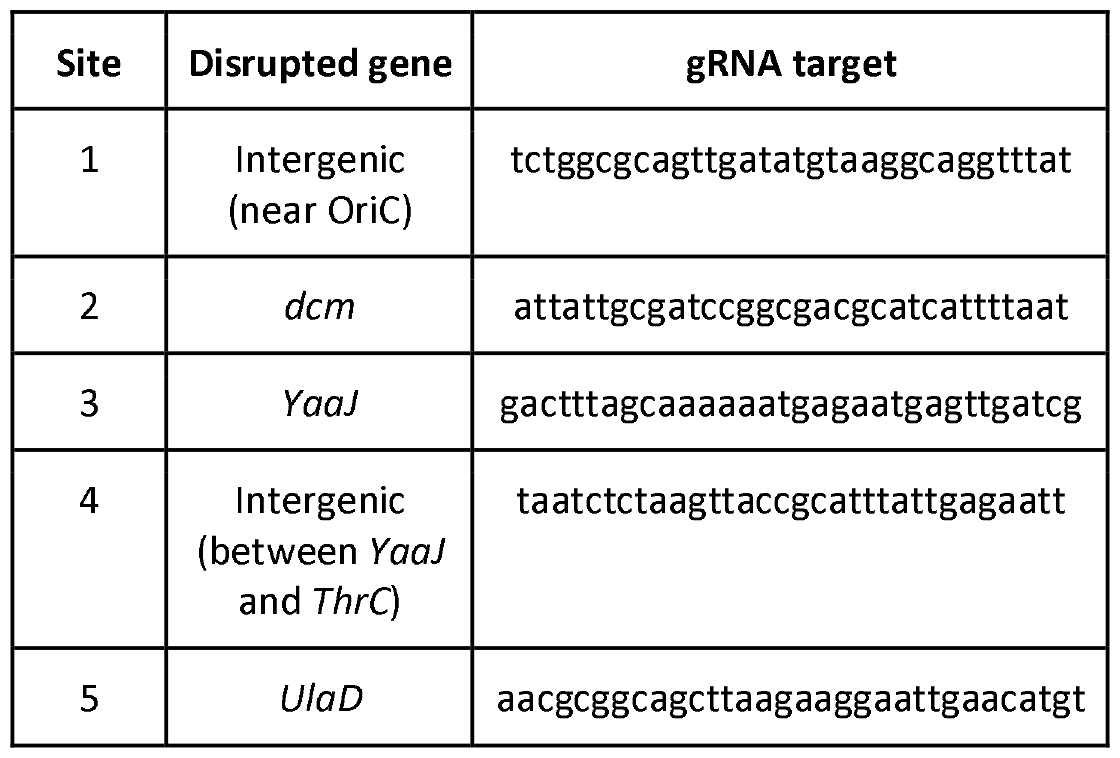
Integration locations.

**Fig. 4.**
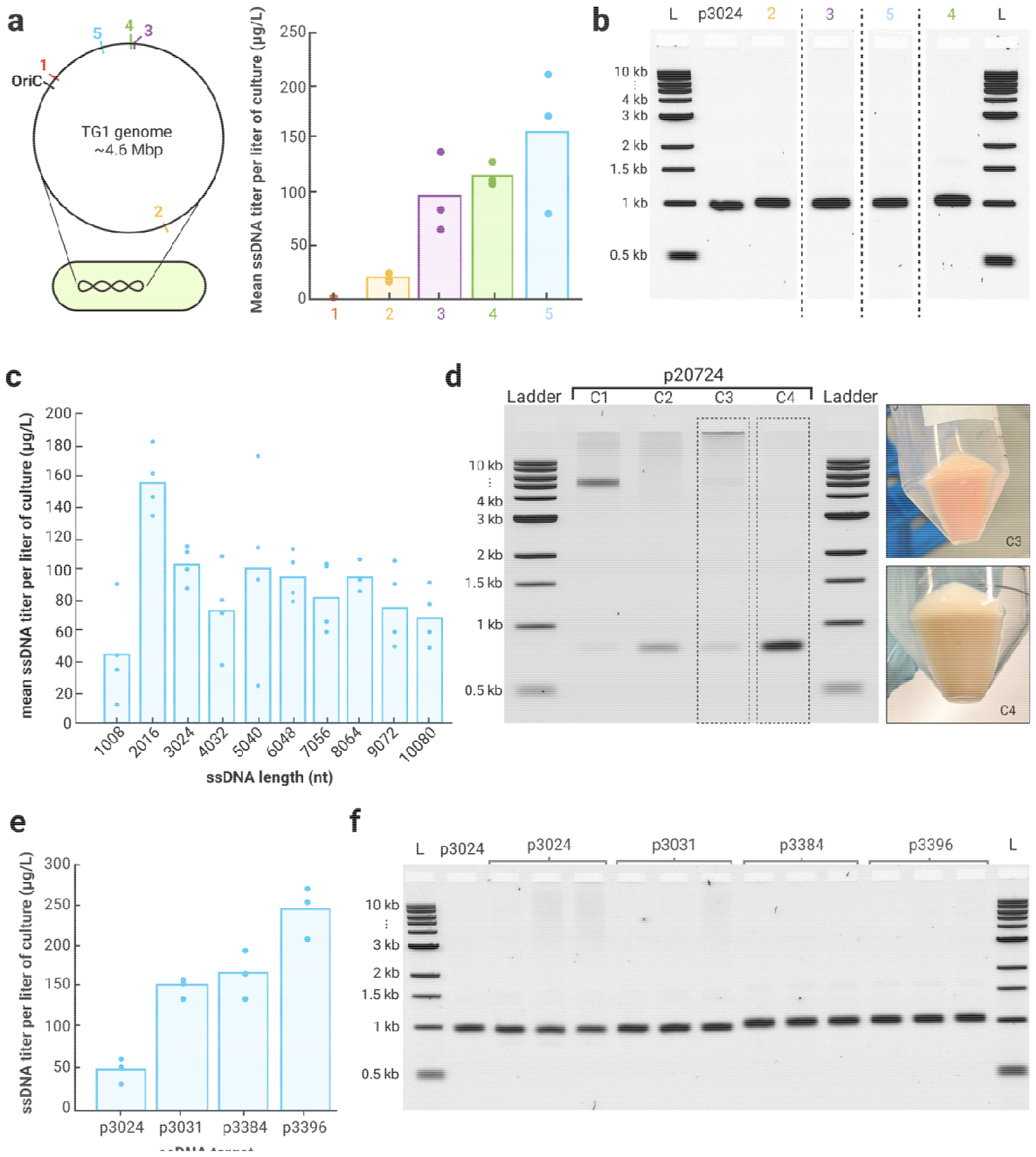
M13mp18-helper strain characterization. **a)** Integration location of M13mp18 genome in the TG1 genome affects ssDNA yield. **b)** Integration location does not affect ssDNA purity. M13mp18- helper strains with integrations into site 1 yielded no ssDNA products so gel results were not included. **c)** ssDNA yield from the M13mp18-helper strain does not show strong dependence on ssDNA length. The 1008-nt ssDNA product does show lower yields than other ssDNA products with lengths ranging from 2008-nt to 10,080-nt. **d)** 20,724-nt ssDNA production is not as consistent as shorter length ssDNA products. Of four tested preps, all preps showed off-target contamination and we found only two produced the correct product. Preps which produced the correct product did show a red pellet, corresponding to the RFP gene encoded in the ssDNA product. **e)** Sequence composition influences ssDNA titer. **f)** Sequence composition does not have a strong effect on ssDNA purity. All ladders are 1 kb ladders from New England Biolabs.

### The length of ssDNA has minimal impact on production titer

Our investigation revealed no substantial correlation between the titer and the length of ssDNA within the range of 2,016 to 10,080 nucleotides. Notably, the ssDNA strand of 1008 nucleotides consistently showed lower titers (refer to Fig. 4c). We designed a series of phagemids encoding ssDNA sequences, increasing in length by increments of 1008 nucleotides. Each subsequent phagemid contained the entirety of the preceding sequence plus an additional 1008 nucleotides. Although the yield for the 1008-nt product was lower, it was producible, unlike the 504-nt product, indicating a minimum threshold for efficient ssDNA synthesis exists between 1008 and 2016 nucleotides.

To explore the upper limits of ssDNA production, we cloned a 20,784-nucleotide target into the pScaf vector. This extended ssDNA sequence included all sequences from the 10,080-nt construct with a red fluorescent protein (RFP) operon incorporated within the additional 10,704 nucleotides. The resulting ssDNA products using this phagemid showed increased variability and the presence of off-target byproducts (Fig. 4d), with only half of the tested colonies displaying the expected band size and all colonies exhibiting additional bands corresponding to the vector backbone. However, the expression of RFP in bacterial cells was correlated with the successful production of the target ssDNA. This outcome hints at the potential instability of long ssDNA sequences within phagemids and suggests that incorporating a secondary selection marker might enhance the quality of ssDNA, particularly for constructs exceeding 10,080 nucleotides.

### Influence of ssDNA sequence on yield and purity

Our investigation into the effects of ssDNA sequence on production yield and purity involved four constructs with lengths ranging from 3024 to 3396 nt. These constructs included a randomly generated sequence (p3024), a segment from the dynein gene’s coding region (p3031), and two sequences constituting complete operons for a B-cell maturation antigen chimeric antigen receptor (p3384, p3396).

Notably, the sequence composition significantly influenced yield; for instance, p3396 resulted in the highest yield at 245 μg/L, while p3024 had the lowest at 49 μg/L. Contrary to initial expectations, complete operon sequences led to higher yields compared to the coding sequence or a random sequence. Despite a high degree of similarity (93%) between the two operon sequences with the highest yields, their yield differed by nearly 50%, underscoring the critical role of sequence in ssDNA production efficiency.

Purity, however, did not exhibit significant variations across the constructs, as demonstrated in Figure 4e and f. Base composition was relatively consistent among the sequences, with each having a base content ranging from 22% to 27% (detailed in Fig. S4).

### Utilization of eScaf-derived ssDNA in DNA nanotechnology applications

To validate the practical applications of our eScaf-derived ssDNA, we employed it in the construction of DNA origami nanostructures. DNA origami is a technique where a long ssDNA scaffold is intricately folded into predetermined shapes with the help of numerous short ssDNA staple strands. We used eScaf to produce a custom 3024-nt scaffold and folded it into a 24 nm × 43 nm × 6 nm rectangular tile using 68 commercially purchased staple strands (Fig. 5a, Fig. S5). Agarose gel electrophoresis and negative stain transmission electron microscopy (TEM) confirmed the integrity and accuracy of the folded structures (Fig. 5b). These data demonstrate that eScaf-derived ssDNA can be used to make DNA nanostructures.

**Fig. 5.**
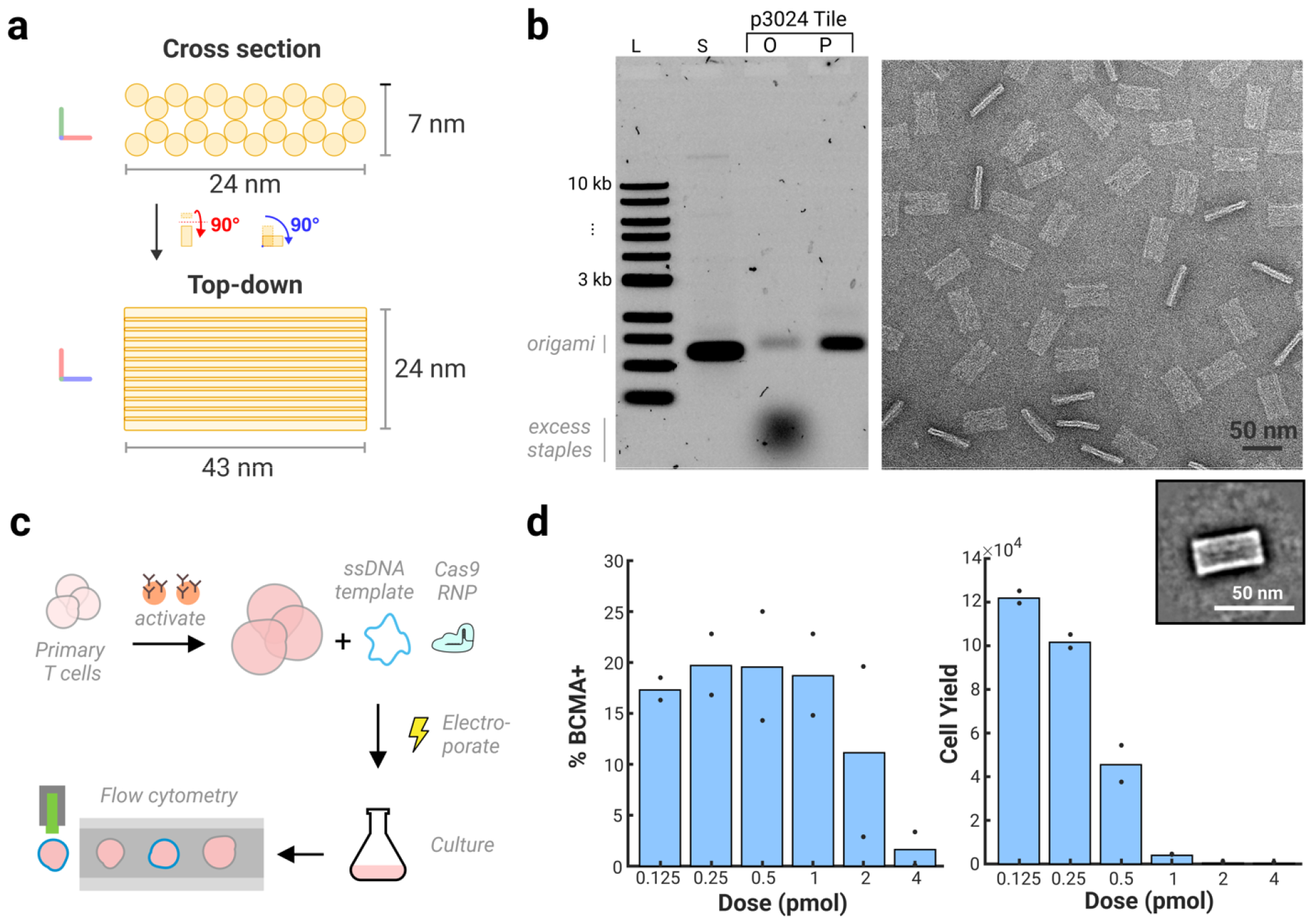
Applications of ssDNA products in biotechnology. a,b) ssDNA products can be used for DNA origami. **a**) Schematic of DNA origami tile made from custom 3024-nt ssDNA scaffold. **b)** *Left:* Gel data of a rectangular DNA origami tile folded from a custom 3024-nt scaffold made using the eScaf system. Lane 3 shows the tile immediately after folding with a well-defined origami band. Lane 4 shows the tile post staple purification. L: NEB 1 kb ladder S: 3024 scaffold O: original origami without purification P: purified origami. *Right:* Negative stain TEM of the purified 3024 tile shows well-formed tiles. “Narrow” tiles show when the tile lands on its side rather than on its face. Box: 2D class average of the tile. **c**,**d) eScaf derived ssDNA products can be used for CRISPR-mediated knock-ins. c)** Schematic of knock-in procedure using eScaf produced BCMA-CAR HDR template for human primary T cell knock-ins. Knock-in efficiency and cell yield were evaluated by flow cytometry. **d)** *Left:* Cells using eScaf derived templates can successfully be knocked-in at all template doses. However, the efficiency of knock-in decreases substantially with template dose. At non-toxic doses, knock-in efficiency using eScaf-derived ssDNA is comparable to published literature^16^. *Right:* Cell yield decreases as increasing amounts of eScaf-derived templates are electroporated. The toxicity of eScaf-derived products may be due to endotoxins from bacterial culture.

### Application of eScaf-derived ssDNA in CRISPR knock-ins for primary T cells

We synthesized a single-stranded CRISPR homology-directed repair (HDR) template for a B-cell maturation antigen chimeric antigen receptor (BCMA-CAR) knock-in, which includes a 1600-nt coding sequence, homology arms approximately 600 nt in length, and additional 500-nt non-coding regions comprising the M13 packaging sequence, and confirmed successful integration via flow cytometry (Fig. 5c).

Flow cytometry analysis revealed that, at low doses (0.125 pmol and 0.25 pmol) of repair template, eScaf-derived ssDNA produced knock-in efficiencies approaching 20% (Fig. 5d), consistent with prior knock-in rates using DNA templates for primary T cell engineering^16^. At higher doses, cell yield decreased dramatically for cells electroporated with eScaf-derived templates (Fig. 5d). Although we purify the ssDNA from the phage proteins using ethanol purification, endotoxins can be retained with DNA in alcohol-based DNA precipitation methods^17^. We suspect that additional residual endotoxins from bacterial production of the templates may cause cell death, contributing to the decreased knock-in efficiency. Despite this drawback, our results demonstrate substantial promise for using eScaf as a cost-effective method to create cssDNA templates for use in gene editing.

## Discussion

Our eScaf helper strain consistently produces long ssDNA with higher purity compared to conventional techniques utilizing helper plasmids or helper phages. Our helper plasmid ssDNA products typically yielded below 60% of the desired ssDNA product, while ssDNA products made by our helper strain typically achieved greater than 90% purity (Fig. S1). For ssDNA lengths ranging from 1,008 to 10,080 nucleotides synthesized via our process, even though occasional faint off-target bands were observed, the purity was sufficient to forgo additional purification steps.

The eScaf system enables the production of ssDNA within a five-day period, including transformation, culture growth, phage-like particle harvesting, and purification. Using standard shake flasks and LB media, we consistently generated BCMA-CAR templates at a concentration of 100 μg per liter of bacterial culture. This translates to an estimated cost of $0.04 per μg of ssDNA, representing a notable cost-efficiency. Further cost reductions may be achieved by optimizing bacterial media and growth conditions, or by employing bioreactors for high-density cultures.

Commercial kits for ssDNA synthesis are currently constrained by their limited production length (typically less than 5000 nucleotides) and high costs compared to our eScaf method. While commercial vendors can synthesize longer strands, the associated expenses and lengthy production times are substantially more burdensome than using eScaf, which offers a faster and more cost-effective solution for long ssDNA synthesis. Additionally, the higher purity of eScaf products compared to those generated by phage or helper entities reduces the amount of labor needed to generate target ssDNA. Our system’s reliance on phagemids and a helper strain—rather than on self-replicating phages— eliminates the risk of contaminating other bacterial cultures, enhancing biosafety and reducing the potential for cross-contamination. The requirement for just a single phagemid avoids the dual antibiotic selection required by helper plasmids systems and thereby reduces reagent cost.

It is important to note, however, that phagemid-based ssDNA production, including eScaf, does not export completely customized ssDNA, as the target sequence must contain the M13 packaging sequences. Recent advances have shortened the size of these packaging sequences to 234 nt, significantly less than the original 2–3 thousand nucleotides^12,18^. Moreover, self-cleaving DNAzymes can then be used to remove the M13 sequences post-harvest, leaving less than 10 nt of residual sequence^19,20^. While eScaf also does not efficiently produce 1000-nt or shorter ssDNA strands, the use of self-cleaving DNAzymes can facilitate the generation of shorter ssDNA from longer ones.

By enabling the inexpensive production of pure ssDNA, our eScaf method holds significant promise for the field of DNA nanotechnology. The DNA origami method depends on ssDNA for constructing nanoscale structures, where precise control over sequence length and composition can be crucial. Considering the potential need for hundreds of unique strands per nanostructure, the affordability of our approach is particularly advantageous for those seeking to explore various design variants, assemble complex structures, or scale up DNA nanodevice production for clinical or industrial applications. Additionally, our method has the potential to revolutionize the assembly of DNA origami by producing long ssDNA strands up to 20,000 nucleotides, enabling the creation of larger nanostructures on a single phagemid. Combining ssDNA staples and scaffolds on a single construct enhances folding reaction efficiency and reduces cost^19^. This could dramatically expand the size and complexity of structures that can be synthesized.

Our method to economically produce custom long ssDNA will also benefit gene-editing applications. First, the efficiency of CRISPR-mediated HDR in mice is significantly higher when using ssDNA templates than when using dsDNA templates^21^. Long ssDNA templates also are less toxic and have increased knock-in efficiency compared to dsDNA templates in human T cell genome editing^2,16^. A secondary benefit from phagemid-based ssDNA production is that the ssDNA produced is naturally circularized due to the intrinsic rolling-circle amplification and ligation steps employed in filamentous phage propagation. Circularized ssDNA has been shown to improve knock-in efficiency over linear ssDNA^22,23^. The increased efficiency of gene editing using cssDNA HDR templates will facilitate CAR-T cell manufacturing and gene therapies and allow insertion of larger gene circuits for new synthetic biology constructs. Applying well-established methods to remove endotoxins from purified DNA may further improve cell viability and knock-in efficiency.

Conventional flask-based growth optimizations have yielded several milligrams of ssDNA per liter of phage culture, suggesting further improvements to our eScaf system are possible. To increase titer, future studies could investigate additional chromosomal integration sites^24^, explore ssDNA sequence variations, and even consider alternative E. coli strains that may offer superior performance^25^. Genetic modifications to the host strain, such as knocking out RecA, may also improve titer and stabilize very large (10,000-nt+) ssDNA production.

Modifications to the M13 genome could also boost ssDNA titer, as evidenced by our observation of improved ssDNA yield by using M13mp18 over M13KO7 and literature indicating that downregulation of pV increases ssDNA yield in standard phage preps^26^. Changes must be made judiciously, due to the delicate balance of the M13 lifecycle in which modifications to the expression of any protein may have significant downstream effects^27^. For example, we designed and built an inducible version of the helper strain by replacing two native M13 promoters with T7lac promoters to test the hypothesis that constitutive phage gene expression resulted in toxicity. Although we observed suppression of ssDNA production prior to the addition of the IPTG inducer, the purity and yield of the exported ssDNA was poor (Fig. S6). Our experiments with inducible helper strains indicate that directed evolution of the M13 genes may be a more effective approach to optimize ssDNA production.

In conclusion, our eScaf method presents a scalable and economical solution akin to bacterial protein manufacturing. By integrating M13 genes into the *E. coli* chromosome, we have markedly enhanced the purity of ssDNA, overcoming a major hurdle towards the production and utilization of ssDNA-dependent biotechnological tools. We anticipate that the high-purity ssDNA amplification methods we have developed will be instrumental in advancing research in gene editing and DNA nanotechnology, among other fields.

## Data availability statement

The data underlying this article are available at: doi.org/10.6084/m9.figshare.25301443

## Acknowledgments

This work was supported by NSF grants ECCS-2036865 and DBI-1548297, and NIH grant R35GM125027. K.S. was supported by a Ruth L. Kirchstein Postdoctoral Fellowship grant no. 1F32GM147967. Plasmids for the Cas-guided transposon system (pSL1777 and pSL1119) were gifts from Samuel H. Sternberg (Addgene plasmid #160731; http://n2t.net/addgene:160731; RRID:Addgene_160731, Addgene plasmid #160732; http://n2t.net/addgene:160732 RRID:Addgene_160732). The phagemid used for the phagemid-helper strain compatibility study (pre_IV_CS6_4323_v1) was a gift from Hendrik Dietz (Addgene plasmid #126859; http://n2t.net/addgene:126859; RRID:Addgene_126859). We thank Leslie Biennen of C3 Science Communication for edits and comments on the manuscript.

## Notes

https://www.doi.org/10.6084/m9.figshare.25301443

### Competing Interest Statement

The authors have declared no competing interest.

### Summary of Updates

We corrected the spelling of an author name, and moved the supplemental figures from the main text into a separate document.

