## Supplementary Information for "Engineering an Escherichia coli strain for production of long single-stranded DNA"

### Supplementary Figures

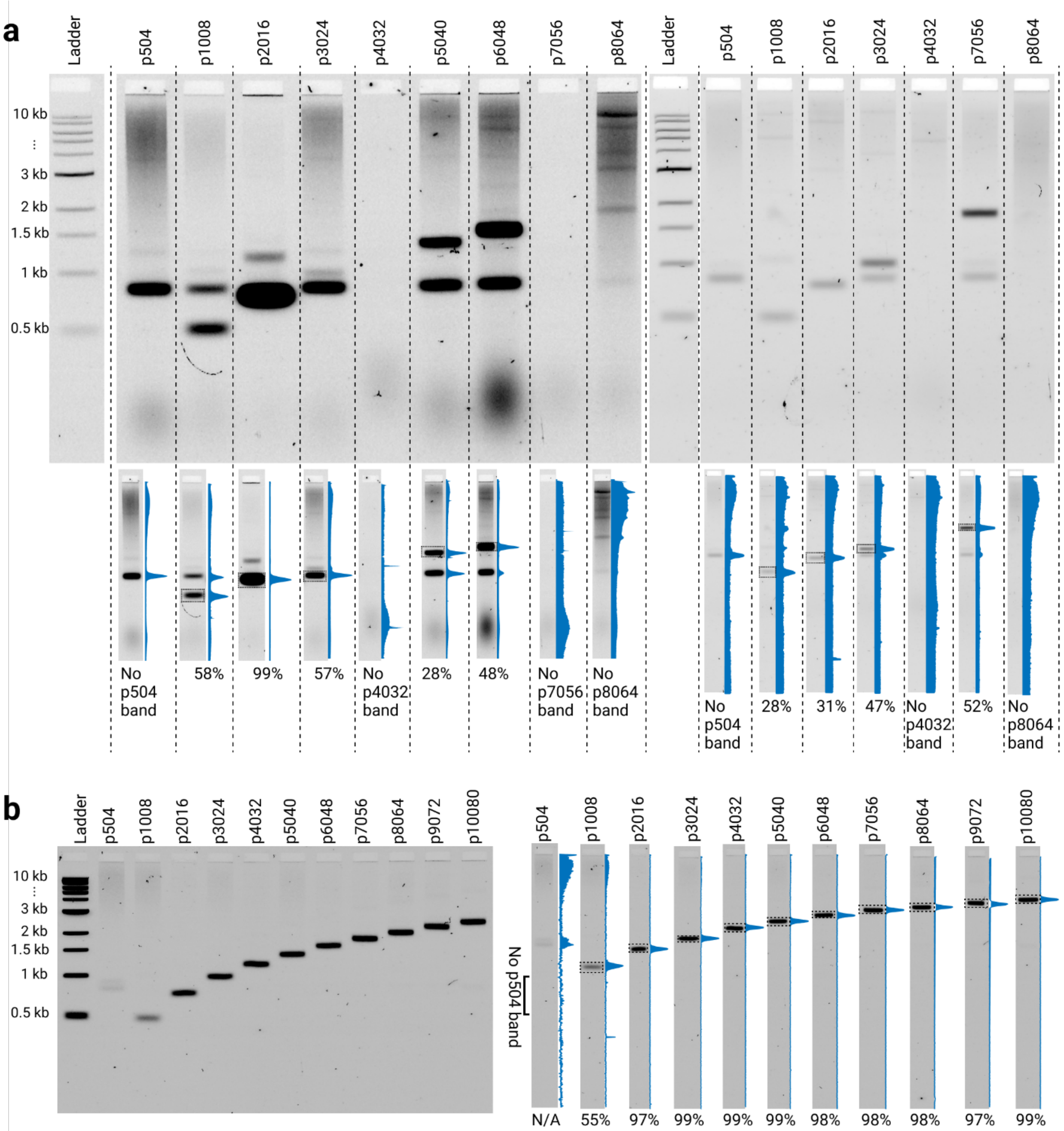


**Fig. S1 ssDNA products generated by helper plasmids are of inconsistent quality**. **a)** Agarose gels showing ssDNA products produced by TG1 bacteria dual-transformed with both M13mp18 helper plasmids and pScaf-based phagemids (left) show inconsistent products. Except for the 2016-nt ssDNA sample, the majority band of all lanes composed of less than 60% of the lane intensity. **b)** Agarose gel from ssDNA produced by eScaf. For ssDNA products 2016-nt and longer, the purity was consistently greater than 95%. All ladders are 1 kb ladders from New England Biolabs. Reported yields were calculated by integrating the intensity of the target band and dividing by the integrated intensities of all features.

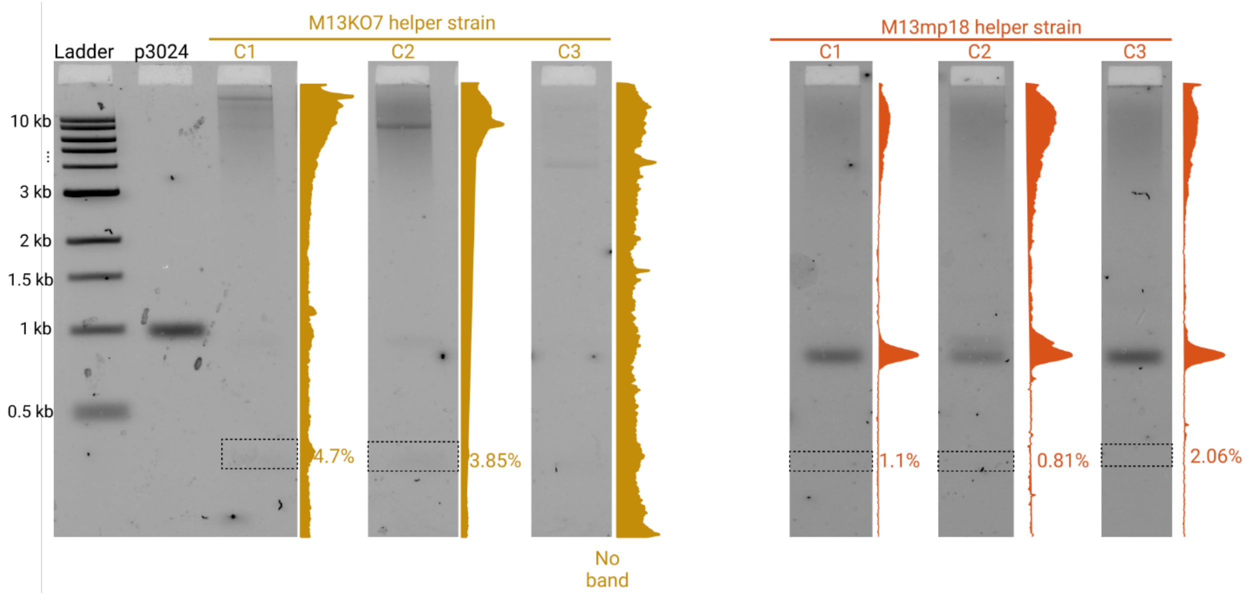


**Fig. S2. Short ssDNA is not well produced by either the M13KO7 or M13mp18 helper strains.** Quantification of 504-nt ssDNA band in M13KO7 helper strains show slightly higher products than in M13mp18 helper strains, but in both strains, 504-nt ssDNA production is poor. Ladder is a 1 kb ladder from New England Biolabs.

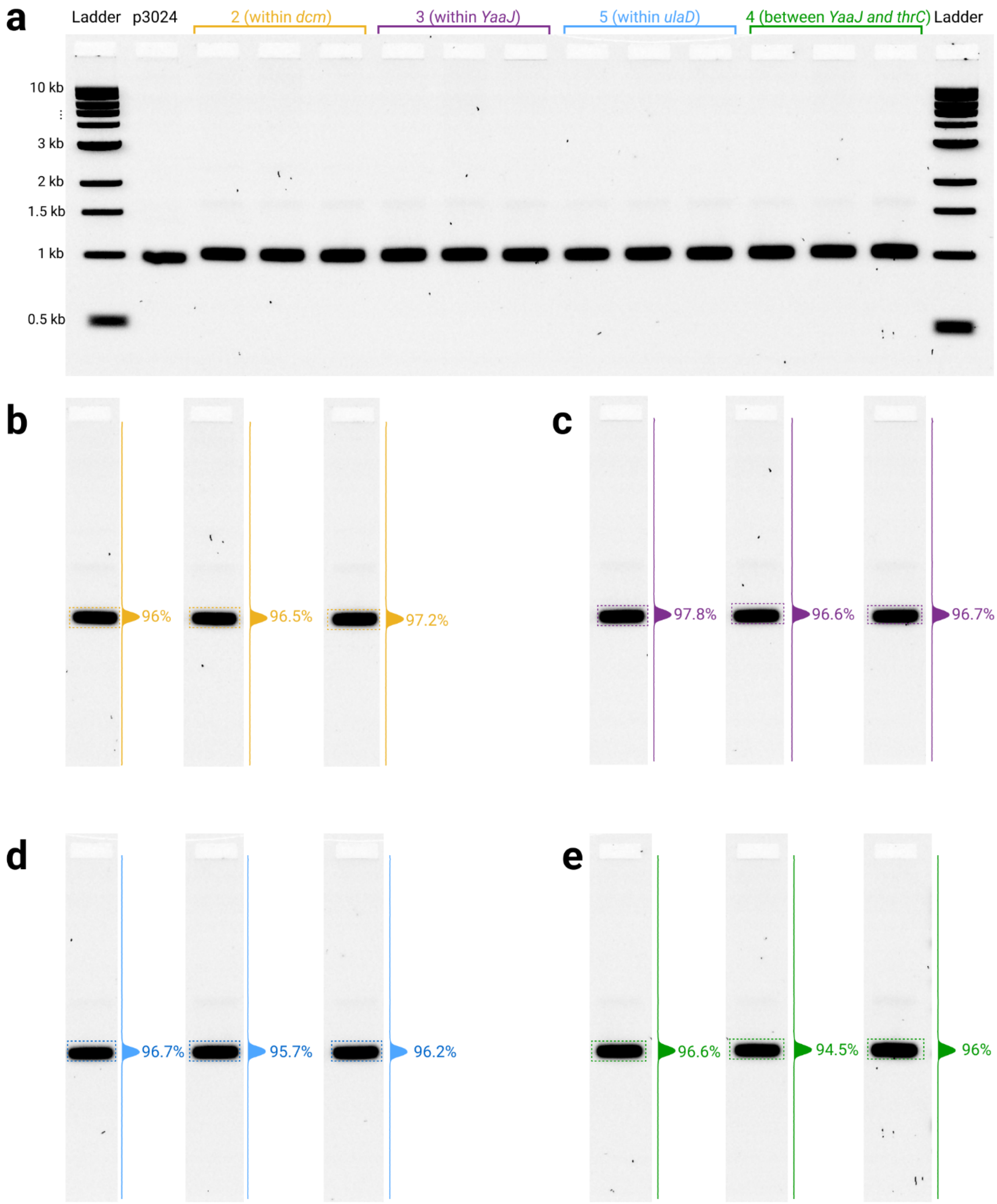


**Fig. S3 Scaffold purity is not affected by integration location. a)** Full agarose gel showing species within ssDNA products integrated into four different locations. Target is a 3024-nt ssDNA product. Ladders are 1 kb ladders from New England Biolabs. **b,c,d,e)** 1D line histograms showing majority of signal (between 95~97%) is located in the target band for all samples over all integration locations.

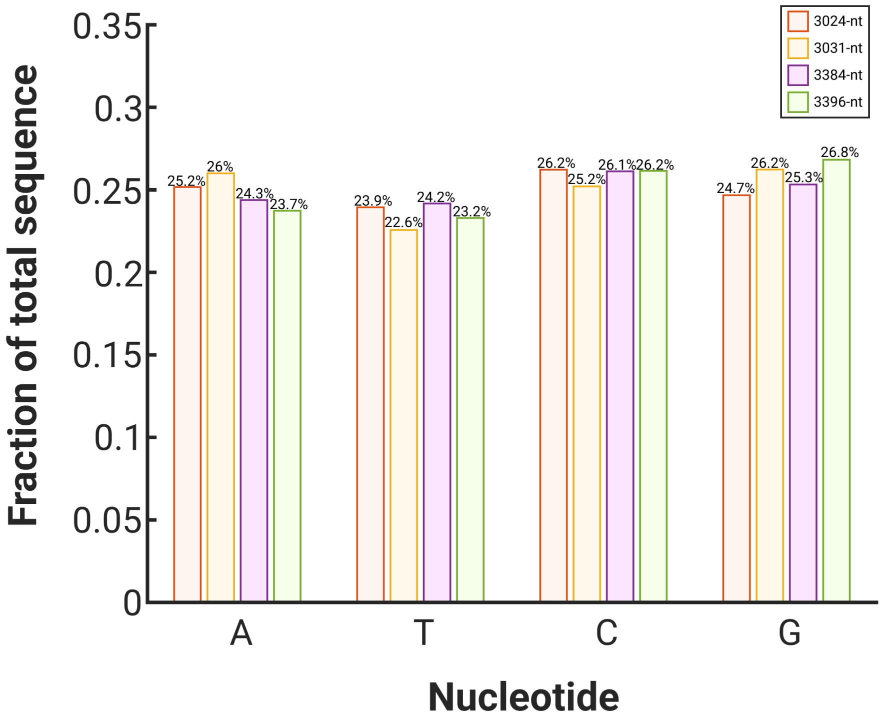


**Fig. S4 Nucleotide composition in ~3000-nt ssDNA products are all similar.** Base composition ranges between 22% and 27%. No nucleotide composition varies more than 3% between any ssDNA product, suggesting that differences in ssDNA titer are not due to composition alone.

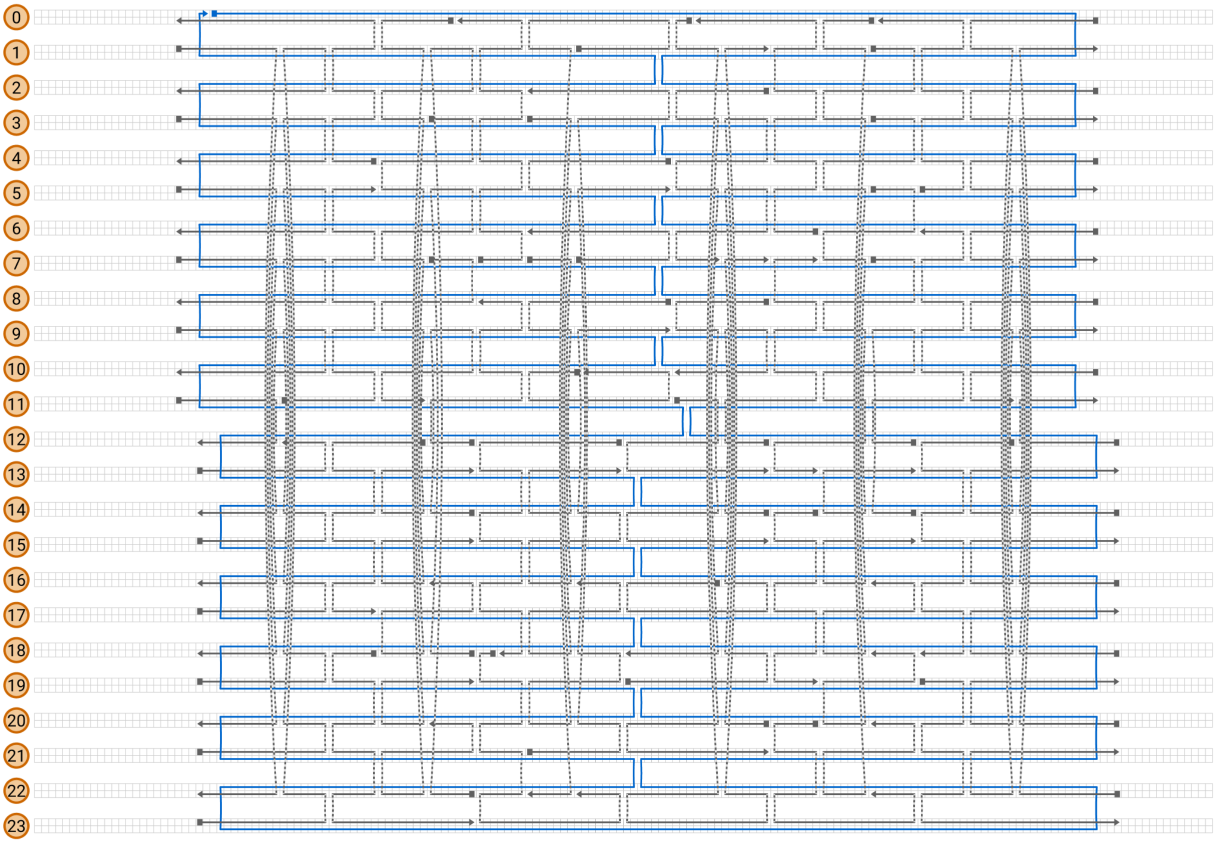


**Fig. S5 Cadnano strand diagram of DNA origami tile**. The custom 3024-nt scaffold strand is represented by the blue line. The staples are represented by grey lines. Dotted grey lines represent staple crossovers.

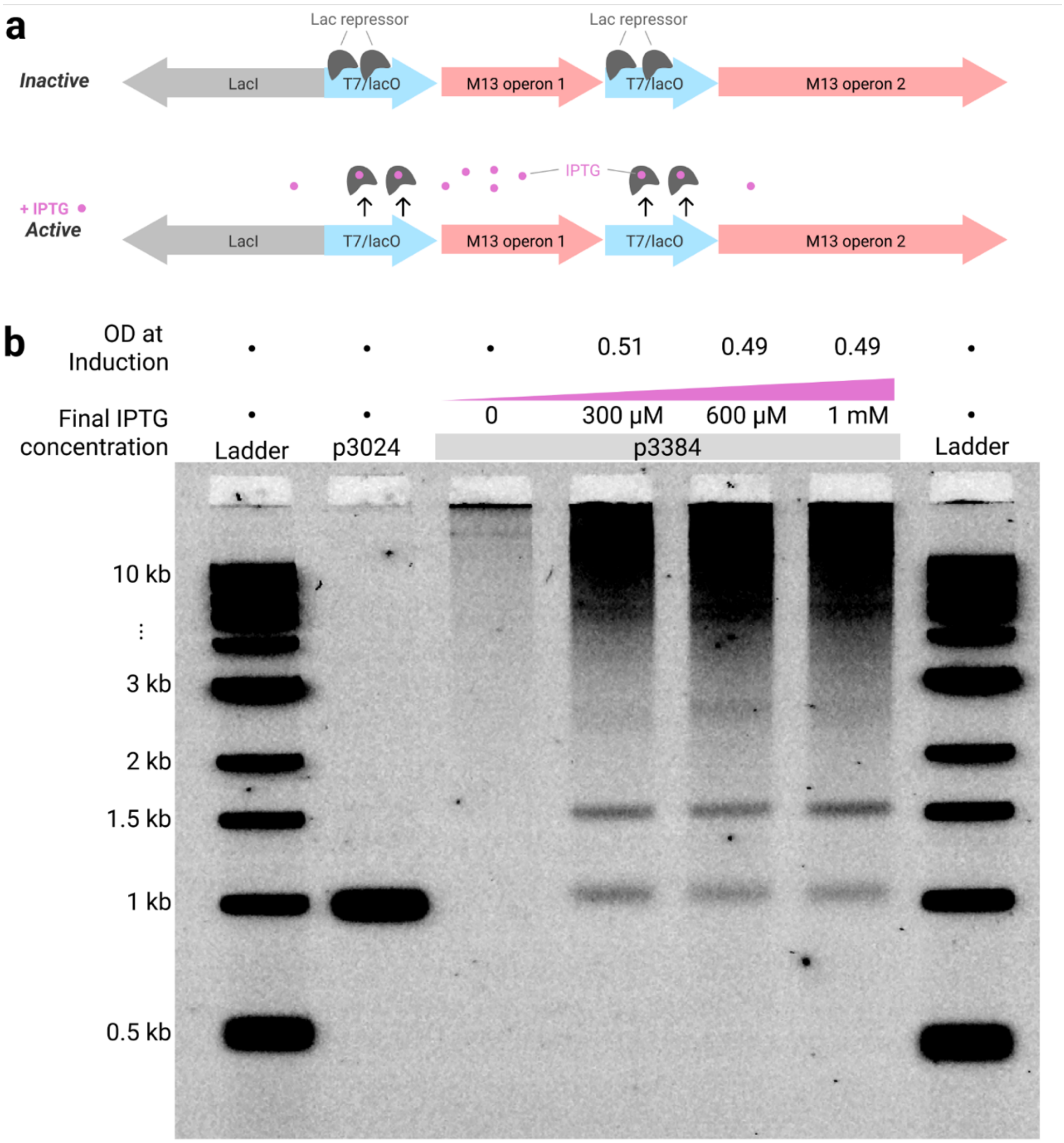


**Fig. S6 T7-regulated M13mp18-helper strains show inducible production of ssDNA, but ssDNA products are impure and have high background**. **a)** Schematic of T7-regulated M13mp18 cassette. We replaced two native promoters in the M13mp18 genome with T7lac and placed the *LacI* gene upstream of the cassette. We integrated this entire cassette into the NEB T7express *E. coli* genome. **b)** We transformed our T7-regulated helper with the p3384 pScaf phagemid. Liquid cultures were grown at 30 °C and when the O.D.600 reached 0.5, induced the cultures at varying levels of IPTG. ssDNA products were collected 3-4 hours post induction and run on a gel alongside a 1 kb ladder from New England Biolabs. Target band (near 1 kb band) appears when induced, but significant off-target species (large smear near well, higher order bands near 1.5 kb band in ladder) also appear.

### DNA sequences used in this work

| Description | Sequence |
|---|---|
| 504-nt ssDNA sequence for length test | aatagtggactcttgttccaaactggaacaacactcaaccctatctcgggctattcttttga<br>tttataagggattttgccgatttcggaacgggtacctacgaagagttccagcagggattcca<br>agaaatggccaatgaagattgctgagagcagcactgtgcaagactggaaaggtggtgaagca<br>atacaccatgaatcccggatcgctgacgcgccctgtagcggcgcattaagcgcggcgggtgt<br>ggtggttacgcgcagcgtgaccgctacacttgccagcgccctagcgcccgctcctttcgctt<br>tcttcccttcctttctcgccacgttcgccggctttccccgtcaagctctaaatcgggggctc<br>cctttagggttccgatttagtgctttacggcacctcgaccccaaaaaacttgatttgggtga<br>tggttcacgtagtgggccatcgccctgatagacggtttttcgccctttgacgttggagtcca<br>cgttcttt |
| 1008-nt ssDNA sequence for length test | aatagtggactcttgttccaaactggaacaacactcaaccctatctcgggctattcttttga<br>tttataagggattttgccgatttcggaacgggtacctacgaagagttccagcagggattcca<br>agaaatggccaatgaagattgtttcccgcacctgggctctttcgcgccaatcgacgttaaac<br>acttgaatttaagatgagcatcgcagaagtggatgatatgcgcgcctacggtccaatcggct<br>gggtgagcgtccctataatctcacggataccgacgaaagtagggcaataatcgcgctgcatg<br>agaacggatagcccggatgcaaaggtggaacgtaattgttaagagaagcaaatacgttaagc<br>tatcatttcccgtaaatcttttacatacgtaaggtggaagtctaaactagcctagctgcaca<br>agcatcggactgcttgccctctatctttgtggaagttcaaggtaatggaagcggtagaacgt<br>atgatcgtcacgcagacctaaagaatcacgccgcgcttacgtccaaccttgggacctacatc<br>tgtacagacccaggtccaggccgaggttatggatgtagatttgacttggctgatctacagtc<br>tcgagttacaatggcgtcactgctacctgctgagagcagcactgtgcaagactggaaaggtg<br>gtgaagcaatacaccatgaatcccggatcgctgacgcgccctgtagcggcgcattaagcgcg<br>gcgggtgtggtggttacgcgcagcgtgaccgctacacttgccagcgccctagcgcccgctcc<br>tttcgctttcttcccttcctttctcgccacgttcgccggctttccccgtcaagctctaaatc<br>gggggctccctttagggttccgatttagtgctttacggcacctcgaccccaaaaaacttgat<br>ttgggtgatggttcacgtagtgggccatcgccctgatagacggtttttcgccctttgacgtt<br>ggagtccacgttcttt |
| 2016-nt ssDNA sequence for length test | aatagtggactcttgttccaaactggaacaacactcaaccctatctcgggctattcttttga<br>tttataagggattttgccgatttcggaacgggtacctacgaagagttccagcagggattcca<br>agaaatggccaatgaagattgtttcccgcacctgggctctttcgcgccaatcgacgttaaac<br>acttgaatttaagatgagcatcgcagaagtggatgatatgcgcgcctacggtccaatcggct<br>gggtgagcgtccctataatctcacggataccgacgaaagtagggcaataatcgcgctgcatg<br>agaacggatagcccggatgcaaaggtggaacgtaattgttaagagaagcaaatacgttaagc<br>tatcatttcccgtaaatcttttacatacgtaaggtggaagtctaaactagcctagctgcaca<br>agcatcggactgcttgccctctatctttgtggaagttcaaggtaatggaagcggtagaacgt<br>atgatcgtcacgcagacctaaagaatcacgccgcgcttacgtccaaccttgggacctacatc<br>tgtacagacccaggtccaggccgaggttatggatgtagatttgacttggctgatctacagtc<br>tcgagttacaatggcgtcactgctacctccatatgaaggataagaggcagccacaactcagc<br>caccatcctaccggaaacgctttaagggcgaacgtgaagggaagtgcctacacgaccctcta<br>tgtgggaagtagtctacgatacacgatgattggaagtgtggcggcataggatgcatctacgg<br>accgacactagttatagaagggtgcagtccttgacgcatacgaatagcgttagctcgcgatt<br>tcccgctgggttactcatggagagcggctcagacgcctagcccatactgaccgtgtaccaac<br>gatccaccaattattttcagaacagcccttttatggcgaagggaaaacagagccttagtatg<br>ttacagtccgctctattgggagatctcgggcaggcggaagatattaattagaaccgcagacc<br>gcatgatgtgccgtcctcaaatggcctggatatcccgtcacacgaattatctcagttgaagt<br>gatcctttagggcccgagagcaccacaagctcccaaccgctgcgaagagtgctaacgagttg<br>tcctatgttaagttattcgtatttaacccacgcccgcaccacctccgggaataatgtccaag<br>atcatatcaaggaaacccagggcacgataagcggcacacagacaaccactttggctagttaa<br>gataacggatactctgacgtacagccggccattcgttctacttttgtgtagtcagatggtcc<br>gtaacaatggcccggctctgacaacaagtataagactgcatgacgggtgggtccacgttaga<br>tgttcgtgaggtagcccatacatcattctttacgaacaagcctctcccgagccagcagagga<br>ggaatcagtgatcgttcacatggcgtacaacaaggcgtgcggcgccgtcggctagcgctgtg<br>aacgtcgcagcgttgattccaaccctctgcccacgacttcgccgctaactggaagcggtatt |

| | |
|---|---|
| | aaacccgaaggttccacgggtggccacttggtgaaggcggcatcgctgagagcagcactgtg<br>caagactggaaaggtggtgaagcaatacaccatgaatcccggatcgctgacgcgccctgtag<br>cggcgcattaagcgcggcgggtgtggtggttacgcgcagcgtgaccgctacacttgccagcg<br>ccctagcgcccgctcctttcgctttcttcccttcctttctcgccacgttcgccggctttccc<br>cgtcaagctctaaatcgggggctccctttagggttccgatttagtgctttacggcacctcga<br>ccccaaaaaacttgatttgggtgatggttcacgtagtgggccatcgccctgatagacggttt<br>ttcgccctttgacgttggagtccacgttcttt |
| 3024-nt ssDNA sequence for length test and composition test | aatagtggactcttgttccaaactggaacaacactcaaccctatctcgggctattcttttga<br>tttataagggattttgccgatttcggaacgggtacctacgaagagttccagcagggattcca<br>agaaatggccaatgaagattgtttcccgcacctgggctctttcgcgccaatcgacgttaaac<br>acttgaatttaagatgagcatcgcagaagtggatgatatgcgcgcctacggtccaatcggct<br>gggtgagcgtccctataatctcacggataccgacgaaagtagggcaataatcgcgctgcatg<br>agaacggatagcccggatgcaaaggtggaacgtaattgttaagagaagcaaatacgttaagc<br>tatcatttcccgtaaatcttttacatacgtaaggtggaagtctaaactagcctagctgcaca<br>agcatcggactgcttgccctctatctttgtggaagttcaaggtaatggaagcggtagaacgt<br>atgatcgtcacgcagacctaaagaatcacgccgcgcttacgtccaaccttgggacctacatc<br>tgtacagacccaggtccaggccgaggttatggatgtagatttgacttggctgatctacagtc<br>tcgagttacaatggcgtcactgctacctccatatgaaggataagaggcagccacaactcagc<br>caccatcctaccggaaacgctttaagggcgaacgtgaagggaagtgcctacacgaccctcta<br>tgtgggaagtagtctacgatacacgatgattggaagtgtggcggcataggatgcatctacgg<br>accgacactagttatagaagggtgcagtccttgacgcatacgaatagcgttagctcgcgatt<br>tcccgctgggttactcatggagagcggctcagacgcctagcccatactgaccgtgtaccaac<br>gatccaccaattattttcagaacagcccttttatggcgaagggaaaacagagccttagtatg<br>ttacagtccgctctattgggagatctcgggcaggcggaagatattaattagaaccgcagacc<br>gcatgatgtgccgtcctcaaatggcctggatatcccgtcacacgaattatctcagttgaagt<br>gatcctttagggcccgagagcaccacaagctcccaaccgctgcgaagagtgctaacgagttg<br>tcctatgttaagttattcgtatttaacccacgcccgcaccacctccgggaataatgtccaag<br>atcatatcaaggaaacccagggcacgataagcggcacacagacaaccactttggctagttaa<br>gataacggatactctgacgtacagccggccattcgttctacttttgtgtagtcagatggtcc<br>gtaacaatggcccggctctgacaacaagtataagactgcatgacgggtgggtccacgttaga<br>tgttcgtgaggtagcccatacatcattctttacgaacaagcctctcccgagccagcagagga<br>ggaatcagtgatcgttcacatggcgtacaacaaggcgtgcggcgccgtcggctagcgctgtg<br>aacgtcgcagcgttgattccaaccctctgcccacgacttcgccgctaactggaagcggtatt<br>aaacccgaaggttccacgggtggccacttggtgaaggcggcatcttaaagttctgacctcaa<br>ttactatggtgccgtcatctaaccaaccattaagaagtagaattgtaaccggttatcggaat<br>cagtgtcgacatactcaatgcccttaccctccgtaactctatgttttccgggttgttcatat<br>cggacatgctatcgcttagacgactcgtctatatagggcgttagtatatcatgggcgtacta<br>tcccggtcaatcgtctacgtcgcaaatcacctccctttgggcaggagagcacttctacactg<br>cctggcgctggcaccgcacaatcaataatgagattgtccggtgagacttacaatcccatcag<br>cgagtctgcaatcccgctgtcacgataacatagtactcgccttctggcggttcctcagctat<br>cctaacgtctatctaaactcaatgtggcgttatttggttcaagcaggcgggtcgtcaacgcg<br>aaacgtaagtcccacacggccgtaacgcaatgtggcaattccagagttaccgatcgaggcca<br>atggcccggccaagccgcaaccgcatcctccgccggtaaaatgactaaatgagtggaaccgt<br>cgcgcttacgctctcggtgcccggaaaactctacagcatattgtctcattggtcctttggcg<br>tatacagacacttagactaactctgactgtcgagtaaattccaatgaacgccagcactcagt<br>acgcacggtgcgtccagggatggttacaaagctagagatgtatgtgctctcatcgaataacc<br>cgtagtatttggacctgaataccctggaatccgagggtgagaccattttacgttatcactgt<br>gacatgctttttccggtcaatacacgggaaactacggacgctatggtcgggataaacgccgc<br>atcatgtcaagctgaggccgctaaacaaagcatcaaggtagatttatttacaagcaaggcct<br>tgcgggcccagtccatgcgctgctgagattacagggattccgtgccagcccttcaacgtcgc<br>tgagagcagcactgtgcaagactggaaaggtggtgaagcaatacaccatgaatcccggatcg<br>ctgacgcgccctgtagcggcgcattaagcgcggcgggtgtggtggttacgcgcagcgtgacc<br>gctacacttgccagcgccctagcgcccgctcctttcgctttcttcccttcctttctcgccac<br>gttcgccggctttccccgtcaagctctaaatcgggggctccctttagggttccgatttagtg<br>ctttacggcacctcgaccccaaaaaacttgatttgggtgatggttcacgtagtgggccatcg<br>ccctgatagacggtttttcgccctttgacgttggagtccacgttcttt |

| | |
|---|---|
| 4032-nt ssDNA sequence for length test | aatagtggactcttgttccaaactggaacaacactcaaccctatctcgggctattcttttga<br>tttataagggattttgccgatttcggaacgggtacctacgaagagttccagcagggattcca<br>agaaatggccaatgaagattgtttcccgcacctgggctctttcgcgccaatcgacgttaaac<br>acttgaatttaagatgagcatcgcagaagtggatgatatgcgcgcctacggtccaatcggct<br>gggtgagcgtccctataatctcacggataccgacgaaagtagggcaataatcgcgctgcatg<br>agaacggatagcccggatgcaaaggtggaacgtaattgttaagagaagcaaatacgttaagc<br>tatcatttcccgtaaatcttttacatacgtaaggtggaagtctaaactagcctagctgcaca<br>agcatcggactgcttgccctctatctttgtggaagttcaaggtaatggaagcggtagaacgt<br>atgatcgtcacgcagacctaaagaatcacgccgcgcttacgtccaaccttgggacctacatc<br>tgtacagacccaggtccaggccgaggttatggatgtagatttgacttggctgatctacagtc<br>tcgagttacaatggcgtcactgctacctccatatgaaggataagaggcagccacaactcagc<br>caccatcctaccggaaacgctttaagggcgaacgtgaagggaagtgcctacacgaccctcta<br>tgtgggaagtagtctacgatacacgatgattggaagtgtggcggcataggatgcatctacgg<br>accgacactagttatagaagggtgcagtccttgacgcatacgaatagcgttagctcgcgatt<br>tcccgctgggttactcatggagagcggctcagacgcctagcccatactgaccgtgtaccaac<br>gatccaccaattattttcagaacagcccttttatggcgaagggaaaacagagccttagtatg<br>ttacagtccgctctattgggagatctcgggcaggcggaagatattaattagaaccgcagacc<br>gcatgatgtgccgtcctcaaatggcctggatatcccgtcacacgaattatctcagttgaagt<br>gatcctttagggcccgagagcaccacaagctcccaaccgctgcgaagagtgctaacgagttg<br>tcctatgttaagttattcgtatttaacccacgcccgcaccacctccgggaataatgtccaag<br>atcatatcaaggaaacccagggcacgataagcggcacacagacaaccactttggctagttaa<br>gataacggatactctgacgtacagccggccattcgttctacttttgtgtagtcagatggtcc<br>gtaacaatggcccggctctgacaacaagtataagactgcatgacgggtgggtccacgttaga<br>tgttcgtgaggtagcccatacatcattctttacgaacaagcctctcccgagccagcagagga<br>ggaatcagtgatcgttcacatggcgtacaacaaggcgtgcggcgccgtcggctagcgctgtg<br>aacgtcgcagcgttgattccaaccctctgcccacgacttcgccgctaactggaagcggtatt<br>aaacccgaaggttccacgggtggccacttggtgaaggcggcatcttaaagttctgacctcaa<br>ttactatggtgccgtcatctaaccaaccattaagaagtagaattgtaaccggttatcggaat<br>cagtgtcgacatactcaatgcccttaccctccgtaactctatgttttccgggttgttcatat<br>cggacatgctatcgcttagacgactcgtctatatagggcgttagtatatcatgggcgtacta<br>tcccggtcaatcgtctacgtcgcaaatcacctccctttgggcaggagagcacttctacactg<br>cctggcgctggcaccgcacaatcaataatgagattgtccggtgagacttacaatcccatcag<br>cgagtctgcaatcccgctgtcacgataacatagtactcgccttctggcggttcctcagctat<br>cctaacgtctatctaaactcaatgtggcgttatttggttcaagcaggcgggtcgtcaacgcg<br>aaacgtaagtcccacacggccgtaacgcaatgtggcaattccagagttaccgatcgaggcca<br>atggcccggccaagccgcaaccgcatcctccgccggtaaaatgactaaatgagtggaaccgt<br>cgcgcttacgctctcggtgcccggaaaactctacagcatattgtctcattggtcctttggcg<br>tatacagacacttagactaactctgactgtcgagtaaattccaatgaacgccagcactcagt<br>acgcacggtgcgtccagggatggttacaaagctagagatgtatgtgctctcatcgaataacc<br>cgtagtatttggacctgaataccctggaatccgagggtgagaccattttacgttatcactgt<br>gacatgctttttccggtcaatacacgggaaactacggacgctatggtcgggataaacgccgc<br>atcatgtcaagctgaggccgctaaacaaagcatcaaggtagatttatttacaagcaaggcct<br>tgcgggcccagtccatgcgctgctgagattacagggattccgtgccagcccttcaacgtctc<br>aagacaactaacaggccttgaattcggccacactcaccggtcccacaatgtgccgggttcgc<br>atcaccgctgcttgggatagtatgcacacaaaagtagcttccacgagcggttgcccaattag<br>atggcgacccgcgtacaggctctaggaggctggaaagtccctctcaccgtcaaatatctgag<br>acgttatacccgacccatcatacgcgaataaagtactagtctcgcctgtcgccctcaggttt<br>accaccaataggacacggaaggcgctctgacaccagccaactagacagacatcggcgaggtg<br>tacttgacgcactttagaagctgctcccttgtgggaaccattgggcaaccgaacatagccgc<br>aatccagtcgcatcatcggtgggtcactgaccgaggattttggcggctcacgccttctgggc<br>tagcttcttctggcttgttagttattgcgtttacgttactctatatcccactaactatctat<br>atacctgtgctttcactacaatggctgcacagttatcttattttagcaaagctttgggttgc<br>ctggagtttcccaaagcgggacttgtaagcccgtcctatatcaggaggcggacgagagacgc<br>agcatgttatcttctaatctgcgaacggtacccggtctttcgtcggctaactagatctgtgt<br>cctaggtattcaaatcgggcgaagtcggtatcgaagaaaagccctttaacgtaaactgtatt<br>cgtcggccgccgttagctcaatgatataactctcatcgcgtgagtcatggcgccacatttta<br>taaaacacgctactatccaatcgaggagatcgctgcaccaataacagtctcaatccaacgaa<br>ctagtatttcatcaccgatccgcgaatcgtgagaccgacccatcgtcttatcgtcctaagca<br>gcacgggccacgacgtgcgaggcggcgtaacttgcgagttctacctatatacttgacaatct |

| | |
|---|---|
| | cagctactccaaacgctgagagcagcactgtgcaagactggaaaggtggtgaagcaatacac<br>catgaatcccggatcgctgacgcgccctgtagcggcgcattaagcgcggcgggtgtggtggt<br>tacgcgcagcgtgaccgctacacttgccagcgccctagcgcccgctcctttcgctttcttcc<br>cttcctttctcgccacgttcgccggctttccccgtcaagctctaaatcgggggctcccttta<br>gggttccgatttagtgctttacggcacctcgaccccaaaaaacttgatttgggtgatggttc<br>acgtagtgggccatcgccctgatagacggtttttcgccctttgacgttggagtccacgttct<br>tt |
| 5040-nt ssDNA sequence for length test | aatagtggactcttgttccaaactggaacaacactcaaccctatctcggctattcttttga<br>tttataagggattttgccgatttcggaacgggtacctacgaagagttccagcagggattcca<br>agaaatggccaatgaagattgtttcccgcacctgggctctttcgcgccaatcgacgttaaac<br>acttgaatttaagatgagcatcgcagaagtggatgatatgcgcgcctacggtccaatcggct<br>gggtgagcgtccctataatctcacggataccgacgaaagtagggcaataatcgcgctgcatg<br>agaacggatagcccggatgcaaaggtggaacgtaattgttaagagaagcaaatacgttaagc<br>tatcatttcccgtaaatcttttacatacgtaaggtggaagtctaaactagcctagctgcaca<br>agcatcggactgcttgccctctatctttgtggaagttcaaggtaatggaagcggtagaacgt<br>atgatcgtcacgcagacctaaagaatcacgccgcgcttacgtccaaccttgggacctacatc<br>tgtacagacccaggtccaggccgaggttatggatgtagatttgacttggctgatctacagtc<br>tcgagttacaatggcgtcactgctacctccatatgaaggataagaggcagccacaactcagc<br>caccatcctaccggaaacgctttaagggcgaacgtgaagggaagtgcctacacgaccctcta<br>tgtgggaagtagtctacgatacacgatgattggaagtgtggcggcataggatgcatctacgg<br>accgacactagttatagaagggtgcagtccttgacgcatacgaatagcgttagctcgcgatt<br>tcccgctgggttactcatggagagcggctcagacgcctagcccatactgaccgtgtaccaac<br>gatccaccaattattttcagaacagcccttttatggcgaagggaaaacagagccttagtatg<br>ttacagtccgctctattgggagatctcgggcaggcggaagatattaattagaaccgcagacc<br>gcatgatgtgccgtcctcaaatggcctggatatcccgtcacacgaattatctcagttgaagt<br>gatcctttagggcccgagagcaccacaagctcccaaccgctgcgaagagtgctaacgagttg<br>tcctatgttaagttattcgtatttaacccacgcccgcaccacctccgggaataatgtccaag<br>atcatatcaaggaaacccagggcacgataagcggcacacagacaaccactttggctagttaa<br>gataacggatactctgacgtacagccggccattcgttctacttttgtgtagtcagatggtcc<br>gtaacaatggcccggctctgacaacaagtataagactgcatgacgggtgggtccacgttaga<br>tgttcgtgaggtagcccatacatcattctttacgaacaagcctctcccgagccagcagagga<br>ggaatcagtgatcgttcacatggcgtacaacaaggcgtgcggcgccgtcggctagcgctgtg<br>aacgtcgcagcgttgattccaaccctctgcccacgacttcgccgctaactggaagcggtatt<br>aaacccgaaggttccacgggtggccacttggtgaaggcggcatcttaaagttctgacctcaa<br>ttactatggtgccgtcatctaaccaaccattaagaagtagaattgtaaccggttatcggaat<br>cagtgtcgacatactcaatgcccttaccctccgtaactctatgttttccgggttgttcatat<br>cggacatgctatcgcttagacgactcgtctatatagggcgttagtatatcatgggcgtacta<br>tcccggtcaatcgtctacgtcgcaaatcacctccctttgggcaggagagcacttctacactg<br>cctggcgctggcaccgcacaatcaataatgagattgtccggtgagacttacaatcccatcag<br>cgagtctgcaatcccgctgtcacgataacatagtactcgccttctggcggttcctcagctat<br>cctaacgtctatctaaactcaatgtggcgttatttggttcaagcaggcgggtcgtcaacgcg<br>aaacgtaagtcccacacggccgtaacgcaatgtggcaattccagagttaccgatcgaggcca<br>atggcccggccaagccgcaaccgcatcctccgccggtaaaatgactaaatgagtggaaccgt<br>cgcgcttacgctctcggtgcccggaaaactctacagcatattgtctcattggtcctttggcg<br>tatacagacacttagactaactctgactgtcgagtaaattccaatgaacgccagcactcagt<br>acgcacggtgcgtccagggatggttacaaagctagagatgtatgtgctctcatcgaataacc<br>cgtagtatttggacctgaataccctggaatccgagggtgagaccattttacgttatcactgt<br>gacatgctttttccggtcaatacacgggaaactacggacgctatggtcgggataaacgccgc<br>atcatgtcaagctgaggccgctaaacaaagcatcaaggtagatttatttacaagcaaggcct<br>tgcgggcccagtccatgcgctgctgagattacagggattccgtgccagcccttcaacgtctc<br>aagacaactaacaggccttgaattcggccacactcaccggtcccacaatgtgccgggttcgc<br>atcaccgctgcttgggatagtatgcacacaaaagtagcttccacgagcggttgcccaattag<br>atggcgacccgcgtacaggctctaggaggctggaaagtccctctcaccgtcaaatatctgag<br>acgttatacccgacccatcatacgcgaataaagtactagtctcgcctgtcgccctcaggttt<br>accaccaataggacacggaaggcgctctgacaccagccaactagacagacatcggcgaggtg<br>tacttgacgcactttagaagctgctcccttgtgggaaccattgggcaaccgaacatagccgc<br>aatccagtcgcatcatcggtgggtcactgaccgaggattttggcggctcacgccttctgggc<br>tagcttcttctggcttgttagttattgcgtttacgttactctatatcccactaactatctat |

| | |
|---|---|
| | atacctgtgctttcactacaatggctgcacagttatcttattttagcaaagctttgggttgc<br>ctggagtttcccaaagcgggacttgtaagcccgtcctatatcaggaggcggacgagagacgc<br>agcatgttatcttctaatctgcgaacggtacccggtctttcgtcggctaactagatctgtgt<br>cctaggtattcaaatcgggcgaagtcggtatcgaagaaaagccctttaacgtaaactgtatt<br>cgtcggccgccgttagctcaatgatataactctcatcgcgtgagtcatggcgccacatttta<br>taaaacacgctactatccaatcgaggagatcgctgcaccaataacagtctcaatccaacgaa<br>ctagtatttcatcaccgatccgcgaatcgtgagaccgacccatcgtcttatcgtcctaagca<br>gcacgggccacgacgtgcgaggcggcgtaacttgcgagttctacctatatacttgacaatct<br>cagctactccaaactgcccatggttggttggccatatgtacaccgagtcctagtacatcctc<br>actggacacgcgttcgcttgttggagagatgaaatccaagatattccttgtagggagctaat<br>cttgccaacactcaaattcctgatgcctcccaaaatacccgggctcaggtcaaaaaagccat<br>gaagcttcaagcccatgctttcttgagtgattatcgctggccgggcgtataagttaatcca<br>gctaactggcgtgtcaacgaaagggtgggacacaatggttttccggctgtctcccagcaagt<br>gtcagaggcatttgcctttctctcaaccaaagcgctacactacacaggtcatcccgtgaaca<br>ttaggtagagttctccagtcagtcctcgacacgagtccagccactcgaacttagtttaaggt<br>cggcgagcaagaccaggtaacgagcaaccaatacatctgtcctttgacccggcatgtcctgc<br>tgtacaggtccgcattagatcagaagtgcggttccatgacgagccacgttccctacaacgaa<br>gcgtaaactagtacccttctacacaggcaccgcccggagtaggaaggattatgcttttgcct<br>ttaggaatttctagattctggtccgtgctgcggcctgcaacgtggacttacttataactgcg<br>gttaggacgattcatctgaaggaatacgctctttttcgactgcagctcgcgtgacgcttggc<br>tgaaaaattgaaactggagcttcctctacggatcaacgtttaactacccactgcctattcct<br>atgtactgatcgtcgagtcttgccaggattccgtcgcgggactcgtatcaacgttcagttga<br>gttgtgtcatgctaaccgcacattgtgagtcaccaagtgtcccatttggtcaactgatctcg<br>caaaaggtaagggccgtagcaaagtcgccagcttcgtcaattgatggcctatttttaactcg<br>ccgcttacggtcggaatctgaacggagacggctgagagcagcactgtgcaagactggaaagg<br>tggtgaagcaatacaccatgaatcccggatcgctgacgcgccctgtagcggcgcattaagcg<br>cggcgggtgtggtggttacgcgcagcgtgaccgctacacttgccagcgccctagcgcccgct<br>cctttcgctttcttcccttcctttctcgccacgttcgccggctttccccgtcaagctctaaa<br>tcgggggctccctttagggttccgatttagtgctttacggcacctcgaccccaaaaaacttg<br>atttgggtgatggttcacgtagtgggccatcgccctgatagacggtttttcgccctttgacg<br>ttggagtccacgttcttt |
| 6048-nt ssDNA sequence for length test | aatagtggactcttgttccaaactggaacaacactcaaccctatctcggctattcttttga<br>tttataagggattttgccgatttcggaacgggtacctacgaagagttccagcagggattcca<br>agaaatggccaatgaagattgtttcccgcacctgggctctttcgcgccaatcgacgttaaac<br>acttgaatttaagatgagcatcgcagaagtggatgatatgcgcgcctacggtccaatcggct<br>gggtgagcgtccctataatctcacggataccgacgaaagtagggcaataatcgcgctgcatg<br>agaacggatagcccggatgcaaaggtggaacgtaattgttaagagaagcaaatacgttaagc<br>tatcatttcccgtaaatcttttacatacgtaaggtggaagtctaaactagcctagctgcaca<br>agcatcggactgcttgccctctatctttgtggaagttcaaggtaatggaagcggtagaacgt<br>atgatcgtcacgcagacctaaagaatcacgccgcgcttacgtccaaccttgggacctacatc<br>tgtacagacccaggtccaggccgaggttatggatgtagatttgacttggctgatctacagtc<br>tcgagttacaatggcgtcactgctacctccatatgaaggataagaggcagccacaactcagc<br>caccatcctaccggaaacgctttaagggcgaacgtgaagggaagtgcctacacgaccctcta<br>tgtgggaagtagtctacgatacacgatgattggaagtgtggcggcataggatgcatctacgg<br>accgacactagttatagaagggtgcagtccttgacgcatacgaatagcgttagctcgcgatt<br>tcccgctgggttactcatggagagcggctcagacgcctagcccatactgaccgtgtaccaac<br>gatccaccaattattttcagaacagcccttttatggcgaagggaaaacagagccttagtatg<br>ttacagtccgctctattgggagatctcgggcaggcggaagatattaattagaaccgcagacc<br>gcatgatgtgccgtcctcaaatggcctggatatcccgtcacacgaattatctcagttgaagt<br>gatcctttagggcccgagagcaccacaagctcccaaccgctgcgaagagtgctaacgagttg<br>tcctatgttaagttattcgtatttaacccacgcccgcaccacctccgggaataatgtccaag<br>atcatatcaaggaaacccagggcacgataagcggcacacagacaaccactttggctagttaa<br>gataacggatactctgacgtacagccggccattcgttctacttttgtgtagtcagatggtcc<br>gtaacaatggcccggctctgacaacaagtataagactgcatgacgggtgggtccacgttaga<br>tgttcgtgaggtagcccatacatcattctttacgaacaagcctctcccgagccagcagagga<br>ggaatcagtgatcgttcacatggcgtacaacaaggcgtgcggcgccgtcggctagcgctgtg<br>aacgtcgcagcgttgattccaaccctctgcccacgacttcgccgctaactggaagcggtatt<br>aaacccgaaggttccacgggtggccacttggtgaaggcggcatcttaaagttctgacctcaa |

| | |
|---|---|
| | ttactatggtgccgtcatctaaccaaccattaagaagtagaattgtaaccggttatcggaat<br>cagtgtcgacatactcaatgcccttaccctccgtaactctatgttttccgggttgttcatat<br>cggacatgctatcgcttagacgactcgtctatatagggcgttagtatatcatgggcgtacta<br>tcccggtcaatcgtctacgtcgcaaatcacctccctttgggcaggagagcacttctacactg<br>cctggcgctggcaccgcacaatcaataatgagattgtccggtgagacttacaatcccatcag<br>cgagtctgcaatcccgctgtcacgataacatagtactcgccttctggcggttcctcagctat<br>cctaacgtctatctaaactcaatgtggcgttatttggttcaagcaggcgggtcgtcaacgcg<br>aaacgtaagtcccacacggccgtaacgcaatgtggcaattccagagttaccgatcgaggcca<br>atggcccggccaagccgcaaccgcatcctccgccggtaaaatgactaaatgagtggaaccgt<br>cgcgcttacgctctcggtgcccggaaaactctacagcatattgtctcattggtcctttggcg<br>tatacagacacttagactaactctgactgtcgagtaaattccaatgaacgccagcactcagt<br>acgcacggtgcgtccagggatggttacaaagctagagatgtatgtgctctcatcgaataacc<br>cgtagtatttggacctgaataccctggaatccgagggtgagaccattttacgttatcactgt<br>gacatgctttttccggtcaatacacgggaaactacggacgctatggtcgggataaacgccgc<br>atcatgtcaagctgaggccgctaaacaaagcatcaaggtagatttatttacaagcaaggcct<br>tgcgggcccagtccatgcgctgctgagattacagggattccgtgccagcccttcaacgtctc<br>aagacaactaacaggccttgaattcggccacactcaccggtcccacaatgtgccgggttcgc<br>atcaccgctgcttgggatagtatgcacacaaaagtagcttccacgagcggttgcccaattag<br>atggcgacccgcgtacaggctctaggaggctggaaagtccctctcaccgtcaaatatctgag<br>acgttatacccgacccatcatacgcgaataaagtactagtctcgcctgtcgccctcaggttt<br>accaccaataggacacggaaggcgctctgacaccagccaactagacagacatcggcgaggtg<br>tacttgacgcactttagaagctgctcccttgtgggaaccattgggcaaccgaacatagccgc<br>aatccagtcgcatcatcggtgggtcactgaccgaggattttggcggctcacgccttctgggc<br>tagcttcttctggcttgttagttattgcgtttacgttactctatatcccactaactatctat<br>atacctgtgctttcactacaatggctgcacagttatcttattttagcaaagctttgggttgc<br>ctggagtttcccaaagcgggacttgtaagcccgtcctatatcaggaggcggacgagagacgc<br>agcatgttatcttctaatctgcgaacggtacccggtctttcgtcggctaactagatctgtgt<br>cctaggtattcaaatcgggcgaagtcggtatcgaagaaaagccctttaacgtaaactgtatt<br>cgtcggccgccgttagctcaatgatataactctcatcgcgtgagtcatggcgccacattta<br>taaaacacgctactatccaatcgaggagatcgctgcaccaataacagtctcaatccaacgaa<br>ctagtatttcatcaccgatccgcgaatcgtgagaccgacccatcgtcttatcgtcctaagca<br>gcacgggccacgacgtgcgaggcggcgtaacttgcgagttctacctatatacttgacaatct<br>cagctactccaaactgcccatggttggttggccatatgtacaccgagtcctagtacatcctc<br>actggacacgcgttcgcttgttggagagatgaaatccaagatattccttgtagggagctaat<br>cttgccaacactcaaattcctgatgcctcccaaaatacccgggctcaggtcaaaaaagccat<br>gaagcttcaagcccatgcttttcttgagtgattatcgctggccgggcgtataagttaatcca<br>gctaactggcgtgtcaacgaaagggtgggacacaatggttttccggctgtctcccagcaagt<br>gtcagaggcatttgcctttctctcaaccaaagcgctacactacacaggtcatcccgtgaaca<br>ttaggtagagttctccagtcagtcctcgacacgagtccagccactcgaacttagtttaaggt<br>cggcgagcaagaccaggtaacgagcaaccaatacatctgtcctttgacccggcatgtcctgc<br>tgtacaggtccgcattagatcagaagtgcggttccatgacgagccacgttccctacaacgaa<br>gcgtaaactagtacccttctacacaggcaccgcccggagtaggaaggattatgcttttgcct<br>ttaggaatttctagattctggtccgtgctgcggcctgcaacgtggacttacttataactgcg<br>gttaggacgattcatctgaaggaatacgctctttttcgactgcagctcgcgtgacgcttggc<br>tgaaaaattgaaactggagcttcctctacggatcaacgtttaactacccactgcctattcct<br>atgtactgatcgtcgagtcttgccaggattccgtcgcgggactcgtatcaacgttcagttga<br>gttgtgtcatgctaaccgcacattgtgagtcaccaagtgtcccatttggtcaactgatctcg<br>caaaaggtaagggccgtagcaaagtcgccagcttcgtcaattgatggcctattttttaactcg<br>ccgcttacggtcggaatctgaacggagacgtctttggcagaatggcgttacgcaccaatcta<br>taaaaagtttttgttggaaaggaggataatttctactggaccggtgttgcgacggaggagat<br>cgaattgctaatcaaccggtatgcacacttccattgctgtgcagttgccctaatcagctatc<br>catccaacataaaactgtctgagtgcttaaacggtcacaccaaatgattgtggtgccgtatc<br>tatagaatatccttagagcgtctgcttcctcgtcacacgagaccggttagtcccaagcacaa<br>taacgaatccagcttctgtttgccttagctccggtgatgcatgtttctgcttccggcggtgc<br>ggatgccacagctgccactgcaggtggaggagaagctgccaagtccaaaccaactacattta<br>ctccaccagattccacccatatcgacttacttccaatagctggttttcgcgttgacccacat<br>tacaatgtatctacgactcaagttattttaatgtatagcgttctgtattcgacccttccata<br>atgccctctatgtgaaactaacaacaatttgaccctcaaactttaagtataccagcttatgg<br>caacagtctgcgagccatggaagggatataacctgacgcagattattgcactgtccaagatc |

| | |
|---|---|
| | cttcatgccacatcttcaggagggcgggtgattagcaccgtaaacagcgggttgatatcatc<br>aagcgaactgcagagaaatccgcgggaacactgggcttagcgcccatctcacccttaaaat<br>taaacgcatctcccggtttcaggcattgctaccctgcgccggctagcgccttcccaatcctg<br>tggcttaagtctactgcgaaacaggttttataacagttccaccgcaatcaggtggccatttg<br>tcctcactctaatcccatccacccgttgatagtcaaagattcctctaataggcccatgaacg<br>tgcaaagttcccaatcgaacccacttggcacatacagtatccggcagctgagagcagcactg<br>tgcaagactggaaaggtggtgaagcaatacaccatgaatcccggatcgctgacgcgccctgt<br>agcggcgcattaagcgcggcgggtgtggtggttacgcgcagcgtgaccgctacacttgccag<br>cgccctagcgcccgctcctttcgctttcttcccttcctttctcgccacgttcgccggctttc<br>cccgtcaagctctaaatcgggggctccctttagggttccgatttagtgctttacggcacctc<br>gaccccaaaaaacttgatttgggtgatggttcacgtagtgggccatcgccctgatagacggt<br>ttttcgccctttgacgttggagtccacgttcttt |
| 7056-nt ssDNA sequence for length test | aatagtggactcttgttccaaactggaacaacactcaaccctatctcgggctattcttttga<br>tttataagggattttgccgatttcggaacgggtacctacgaagagttccagcagggattcca<br>agaaatggccaatgaagattgtttcccgcacctgggctctttcgcgccaatcgacgttaaac<br>acttgaatttaagatgagcatcgcagaagtggatgatatgcgcgcctacggtccaatcggct<br>gggtgagcgtccctataatctcacggataccgacgaaagtagggcaataatcgcgctgcatg<br>agaacggatagcccggatgcaaaggtggaacgtaattgttaagagaagcaaatacgttaagc<br>tatcatttcccgtaaatcttttacatacgtaaggtggaagtctaaactagcctagctgcaca<br>agcatcggactgcttgccctctatctttgtggaagttcaaggtaatggaagcggtagaacgt<br>atgatcgtcacgcagacctaaagaatcacgccgcgcttacgtccaaccttgggacctacatc<br>tgtacagacccaggtccaggccgaggttatggatgtagatttgacttggctgatctacagtc<br>tcgagttacaatggcgtcactgctacctccatatgaaggataagaggcagccacaactcagc<br>caccatcctaccggaaacgctttaagggcgaacgtgaagggaagtgcctacacgaccctcta<br>tgtgggaagtagtctacgatacacgatgattggaagtgtggcggcataggatgcatctacgg<br>accgacactagttatagaagggtgcagtccttgacgcatacgaatagcgttagctcgcgatt<br>tcccgctgggttactcatggagagcggctcagacgcctagcccatactgaccgtgtaccaac<br>gatccaccaattattttcagaacagcccttttatggcgaagggaaaacagagccttagtatg<br>ttacagtccgctctattgggagatctcgggcaggcggaagatattaattagaaccgcagacc<br>gcatgatgtgccgtcctcaaatggcctggatatcccgtcacacgaattatctcagttgaagt<br>gatcctttagggcccgagagcaccacaagctcccaaccgctgcgaagagtgctaacgagttg<br>tcctatgttaagttattcgtatttaacccacgcccgcaccacctccgggaataatgtccaag<br>atcatatcaaggaaacccagggcacgataagcggcacacagacaaccactttggctagttaa<br>gataacggatactctgacgtacagccggccattcgttctacttttgtgtagtcagatggtcc<br>gtaacaatggcccggctctgacaacaagtataagactgcatgacgggtgggtccacgttaga<br>tgttcgtgaggtagcccatacatcattctttacgaacaagcctctcccgagccagcagagga<br>ggaatcagtgatcgttcacatggcgtacaacaaggcgtgcggcgccgtcggctagcgctgtg<br>aacgtcgcagcgttgattccaaccctctgcccacgacttcgccgctaactggaagcggtatt<br>aaacccgaaggttccacgggtggccacttggtgaaggcggcatcttaaagttctgacctcaa<br>ttactatggtgccgtcatctaaccaaccattaagaagtagaattgtaaccggttatcggaat<br>cagtgtcgacatactcaatgcccttaccctccgtaactctatgttttccgggttgttcatat<br>cggacatgctatcgcttagacgactcgtctatatagggcgttagtatatcatgggcgtacta<br>tcccggtcaatcgtctacgtcgcaaatcacctccctttgggcaggagagcacttctacactg<br>cctggcgctggcaccgcacaatcaataatgagattgtccggtgagacttacaatcccatcag<br>cgagtctgcaatcccgctgtcacgataacatagtactcgccttctggcggttcctcagctat<br>cctaacgtctatctaaactcaatgtggcgttatttggttcaagcaggcgggtcgtcaacgcg<br>aaacgtaagtcccacacggccgtaacgcaatgtggcaattccagagttaccgatcgaggcca<br>atggcccggccaagccgcaaccgcatcctccgccggtaaaatgactaaatgagtggaaccgt<br>cgcgcttacgctctcggtgcccggaaaactctacagcatattgtctcattggtcctttggcg<br>tatacagacacttagactaactctgactgtcgagtaaattccaatgaacgccagcactcagt<br>acgcacggtgcgtccagggatggttacaaagctagagatgtatgtgctctcatcgaataacc<br>cgtagtatttggacctgaataccctggaatccgagggtgagaccattttacgttatcactgt<br>gacatgctttttccggtcaatacacgggaaactacggacgctatggtcgggataaacgccgc<br>atcatgtcaagctgaggccgctaaacaaagcatcaaggtagatttatttacaagcaaggcct<br>tgcgggcccagtccatgcgctgctgagattacagggattccgtgccagcccttcaacgtctc<br>aagacaactaacaggccttgaattcggccacactcaccggtcccacaatgtgccgggttcgc<br>atcaccgctgcttgggatagtatgcacacaaaagtagcttccacgagcggttgcccaattag<br>atggcgacccgcgtacaggctctaggaggctggaaagtccctctcaccgtcaaatatctgag |

| | |
|---|---|
| | acgttatacccgacccatcatacgcgaataaagtactagtctcgcctgtcgccctcaggttt<br>accaccaataggacacggaaggcgctctgacaccagccaactagacagacatcggcgaggtg<br>tacttgacgcactttagaagctgctcccttgtgggaaccattgggcaaccgaacatagccgc<br>aatccagtcgcatcatcggtgggtcactgaccgaggattttggcggctcacgccttctgggc<br>tagcttcttctggcttgttagttattgcgtttacgttactctatatcccactaactatctat<br>atacctgtgctttcactacaatggctgcacagttatcttattttagcaaagctttgggttgc<br>ctggagtttcccaaagcgggacttgtaagcccgtcctatatcaggaggcggacgagagacgc<br>agcatgttatcttctaatctgcgaacggtacccggtctttcgtcggctaactagatctgtgt<br>cctaggtattcaaatcgggcgaagtcggtatcgaagaaaagccctttaacgtaaactgtatt<br>cgtcggccgccgttagctcaatgatataactctcatcgcgtgagtcatggcgccacatttta<br>taaaacacgctactatccaatcgaggagatcgctgcaccaataacagtctcaatccaacgaa<br>ctagtatttcatcaccgatccgcgaatcgtgagaccgacccatcgtcttatcgtcctaagca<br>gcacgggccacgacgtgcgaggcggcgtaacttgcgagttctacctatatacttgacaatct<br>cagctactccaaactgcccatggttggttggccatatgtacaccgagtcctagtacatcctc<br>actggacacgcgttcgcttgttggagagatgaaatccaagatattccttgtagggagctaat<br>cttgccaacactcaaattcctgatgcctcccaaaatacccgggctcaggtcaaaaaagccat<br>gaagcttcaagcccatgcttttcttgagtgattatcgctggccgggcgtataagttaatcca<br>gctaactggcgtgtcaacgaaagggtgggacacaatggttttccggctgtctcccagcaagt<br>gtcagaggcatttgcctttctctcaaccaaagcgctacactacacaggtcatcccgtgaaca<br>ttaggtagagttctccagtcagtcctcgacacgagtccagccactcgaacttagtttaaggt<br>cggcgagcaagaccaggtaacgagcaaccaatacatctgtcctttgacccggcatgtcctgc<br>tgtacaggtccgcattagatcagaagtgcggttccatgacgagccacgttccctacaacgaa<br>gcgtaaactagtacccttctacacaggcaccgcccggagtaggaaggattatgcttttgcct<br>ttaggaatttctagattctggtccgtgctgcggcctgcaacgtggacttacttataactgcg<br>gttaggacgattcatctgaaggaatacgctctttttcgactgcagctcgcgtgacgcttggc<br>tgaaaaattgaaactggagcttcctctacggatcaacgtttaactacccactgcctattcct<br>atgtactgatcgtcgagtcttgccaggattccgtcgcgggactcgtatcaacgttcagttga<br>gttgtgtcatgctaaccgcacattgtgagtcaccaagtgtcccatttggtcaactgatctcg<br>caaaaggtaagggccgtagcaaagtcgccagcttcgtcaattgatggcctatttttaactcg<br>ccgcttacggtcggaatctgaacggagacgtctttggcagaatggcgttacgcaccaatcta<br>taaaaagtttttgttggaaaggaggataatttctactggaccggtgttgcgacggaggagat<br>cgaattgctaatcaaccggtatgcacacttccattgctgtgcagttgccctaatcagctatc<br>catccaacataaaactgtctgagtgcttaaacggtcacaccaaatgattgtggtgccgtatc<br>tatagaatatccttagagcgtctgcttcctcgtcacacgagaccggttagtcccaagcacaa<br>taacgaatccagcttctgtttgccttagctccggtgatgcatgtttctgcttccggcggtgc<br>ggatgccacagctgccactgcaggtggaggagaagctgccaagtccaaaccaactacattta<br>ctccaccagattccacccatatcgacttacttccaatagctggttttcgcgttgacccacat<br>tacaatgtatctacgactcaagttattttaatgtatagcgttctgtattcgacccttccata<br>atgccctctatgtgaaactaacaacaatttgaccctcaaactttaagtataccagcttatgg<br>caacagtctgcgagccatggaagggatataacctgacgcagattattgcactgtccaagatc<br>cttcatgccacatcttcaggagggcgggtgattagcaccgtaaacagcgggttgatatcatc<br>aagcgaactgcagagaaatccgcgggaacactgggcttagcgcccatctcacccttaaaaat<br>taaacgcatctcccggtttcaggcattgctaccctgcgccggctagcgccttcccaatcctg<br>tggcttaagtctactgcgaaacaggttttataacagttccaccgcaatcaggtggccatttg<br>tcctcactctaatcccatccacccgttgatagtcaaagattcctctaataggcccatgaacg<br>tgcaaagttcccaatcgaacccacttggcacatacagtatccggcaaatgttatatcaacaa<br>gtcgcctgaacgtgccgcaacaacggatcaactgtagcttcgtgctgcctcagatgcatggc<br>tcgtgcccttctgttcgtgctgcatcggttgtatctcaatgactggactcagcatcgtagct<br>aagtaggcagggattttagtggtgtctaaagaataccggcggtaggcccaaaaaattacctc<br>gttcacccactaaagagatatcccgtacatctactatctactcaggaagatcaccactctag<br>cgtgggagccgctataatggatgcaggcagcccgggttagcgtgatgaaggacgttttaagt<br>tactactactggagttgcgggcgcaagacgatggctaagtaagagcccagagtttaggcctt<br>gtctaaaccgtaatgaaactgacatcggtagtcaatgtgtcgacgagttttgatttcagtat<br>atacgtactgttaaccgacgtctggatgtcagaaatttcgtgcatgtggcaggctcggttcg<br>aggaatctcgttccggaagttagggtatcggcgagggaactagtataaggactcgactgatg<br>catggctcagctaacagcgggcactctatgtcctaaggatagtaagaggagcaggacaacca<br>tccgggtgtaacgggttgatgcaagcgacatactaacaatgcctaggatagctgtgccatca<br>ggcggaaaagaatccaatatgatggtgctcaggactcattattacaatatagtacatttacc<br>agagaggtcccgcgggtcgccgaacacctacgcgaccctataagtttcttacactatcgata |

| | |
|---|---|
| | atggagaaagcttatttgagggactacgatcttttacacccatggactttcagccgagatat<br>caaaatcgtagttatgttgtagcctgtagatttgtattcacgggctgtactttagccgagga<br>cagacccatatctgcttataataggtcatcatccctacattgtgtgagccagtctccaccgc<br>gctgagagcagcactgtgcaagactggaaaggtggtgaagcaatacaccatgaatcccggat<br>cgctgacgcgccctgtagcggcgcattaagcgcggcgggtgtggtggttacgcgcagcgtga<br>ccgctacacttgccagcgccctagcgcccgctcctttcgctttcttcccttcctttctcgcc<br>acgttcgccggctttccccgtcaagctctaaatcgggggctccctttagggttccgatttag<br>tgctttacggcacctcgaccccaaaaaacttgatttgggtgatggttcacgtagtgggccat<br>cgccctgatagacggtttttcgccctttgacgttggagtccacgttcttt |
| 8064-nt ssDNA sequence for length test | aatagtggactcttgttccaaactggaacaacactcaaccctatctcggctattcttttga<br>tttataagggattttgccgatttcggaacgggtacctacgaagagttccagcagggattcca<br>agaaatggccaatgaagattgtttcccgcacctgggctctttcgcgccaatcgacgttaaac<br>acttgaatttaagatgagcatcgcagaagtggatgatatgcgcgcctacggtccaatcggct<br>gggtgagcgtccctataatctcacggataccgacgaaagtagggcaataatcgcgctgcatg<br>agaacggatagcccggatgcaaaggtggaacgtaattgttaagagaagcaaatacgttaagc<br>tatcatttcccgtaaatcttttacatacgtaaggtggaagtctaaactagcctagctgcaca<br>agcatcggactgcttgccctctatctttgtggaagttcaaggtaatggaagcggtagaacgt<br>atgatcgtcacgcagacctaaagaatcacgccgcgcttacgtccaaccttgggacctacatc<br>tgtacagacccaggtccaggccgaggttatggatgtagatttgacttggctgatctacagtc<br>tcgagttacaatggcgtcactgctacctccatatgaaggataagaggcagccacaactcagc<br>caccatcctaccggaaacgctttaagggcgaacgtgaagggaagtgcctacacgaccctcta<br>tgtgggaagtagtctacgatacacgatgattggaagtgtggcggcataggatgcatctacgg<br>accgacactagttatagaagggtgcagtccttgacgcatacgaatagcgttagctcgcgatt<br>tcccgctgggttactcatggagagcggctcagacgcctagcccatactgaccgtgtaccaac<br>gatccaccaattattttcagaacagcccttttatggcgaagggaaaacagagccttagtatg<br>ttacagtccgctctattgggagatctcgggcaggcggaagatattaattagaaccgcagacc<br>gcatgatgtgccgtcctcaaatggcctggatatcccgtcacacgaattatctcagttgaagt<br>gatcctttagggcccgagagcaccacaagctcccaaccgctgcgaagagtgctaacgagttg<br>tcctatgttaagttattcgtatttaacccacgcccgcaccacctccgggaataatgtccaag<br>atcatatcaaggaaacccagggcacgataagcggcacacagacaaccactttggctagttaa<br>gataacggatactctgacgtacagccggccattcgttctacttttgtgtagtcagatggtcc<br>gtaacaatggcccggctctgacaacaagtataagactgcatgacgggtgggtccacgttaga<br>tgttcgtgaggtagcccatacatcattctttacgaacaagcctctcccgagccagcagagga<br>ggaatcagtgatcgttcacatggcgtacaacaaggcgtgcggcgccgtcggctagcgctgtg<br>aacgtcgcagcgttgattccaaccctctgcccacgacttcgccgctaactggaagcggtatt<br>aaacccgaaggttccacgggtggccacttggtgaaggcggcatcttaaagttctgacctcaa<br>ttactatggtgccgtcatctaaccaaccattaagaagtagaattgtaaccggttatcggaat<br>cagtgtcgacatactcaatgcccttaccctccgtaactctatgttttccgggttgttcatat<br>cggacatgctatcgcttagacgactcgtctatatagggcgttagtatatcatgggcgtacta<br>tcccggtcaatcgtctacgtcgcaaatcacctccctttgggcaggagagcacttctacactg<br>cctggcgctggcaccgcacaatcaataatgagattgtccggtgagacttacaatcccatcag<br>cgagtctgcaatcccgctgtcacgataacatagtactcgccttctggcggttcctcagctat<br>cctaacgtctatctaaactcaatgtggcgttatttggttcaagcaggcgggtcgtcaacgcg<br>aaacgtaagtcccacacggccgtaacgcaatgtggcaattccagagttaccgatcgaggcca<br>atggcccggccaagccgcaaccgcatcctccgccggtaaaatgactaaatgagtggaaccgt<br>cgcgcttacgctctcggtgcccggaaaactctacagcatattgtctcattggtcctttggcg<br>tatacagacacttagactaactctgactgtcgagtaaattccaatgaacgccagcactcagt<br>acgcacggtgcgtccagggatggttacaaagctagagatgtatgtgctctcatcgaataacc<br>cgtagtatttggacctgaataccctggaatccgagggtgagaccattttacgttatcactgt<br>gacatgctttttccggtcaatacacgggaaactacggacgctatggtcgggataaacgccgc<br>atcatgtcaagctgaggccgctaaacaaagcatcaaggtagatttatttacaagcaaggcct<br>tgcgggcccagtccatgcgctgctgagattacagggattccgtgccagcccttcaacgtctc<br>aagacaactaacaggccttgaattcggccacactcaccggtcccacaatgtgccgggttcgc<br>atcaccgctgcttgggatagtatgcacacaaaagtagcttccacgagcggttgcccaattag<br>atggcgacccgcgtacaggctctaggaggctggaaagtccctctcaccgtcaaatatctgag<br>acgttatacccgacccatcatacgcgaataaagtactagtctcgcctgtcgccctcaggttt<br>accaccaataggacacggaaggcgctctgacaccagccaactagacagacatcggcgaggtg<br>tacttgacgcactttagaagctgctcccttgtgggaaccattgggcaaccgaacatagccgc |

| | |
|---|---|
| | aatccagtcgcatcatcggtgggtcactgaccgaggattttggcggctcacgccttctgggc<br>tagcttcttctggcttgttagttattgcgtttacgttactctatatcccactaactatctat<br>atacctgtgctttcactacaatggctgcacagttatcttattttagcaaagctttgggttgc<br>ctggagtttcccaaagcgggacttgtaagcccgtcctatatcaggaggcggacgagagacgc<br>agcatgttatcttctaatctgcgaacggtacccggtctttcgtcggctaactagatctgtgt<br>cctaggtattcaaatcgggcgaagtcggtatcgaagaaaagccctttaacgtaaactgtatt<br>cgtcggccgccgttagctcaatgatataactctcatcgcgtgagtcatggcgccacattta<br>taaaacacgctactatccaatcgaggagatcgctgcaccaataacagtctcaatccaacgaa<br>ctagtatttcatcaccgatccgcgaatcgtgagaccgacccatcgtcttatcgtcctaagca<br>gcacgggccacgacgtgcgaggcggcgtaacttgcgagttctacctatatacttgacaatct<br>cagctactccaaactgcccatggttggttggccatatgtacaccgagtcctagtacatcctc<br>actggacacgcgttcgcttgttggagagatgaaatccaagatattccttgtagggagctaat<br>cttgccaacactcaaattcctgatgcctcccaaaatacccgggctcaggtcaaaaaagccat<br>gaagcttcaagcccatgcttttcttgagtgattatcgctggccgggcgtataagttaatcca<br>gctaactggcgtgtcaacgaaagggtgggacacaatggttttccggctgtctcccagcaagt<br>gtcagaggcatttgcctttctctcaaccaaagcgctacactacacaggtcatcccgtgaaca<br>ttaggtagagttctccagtcagtcctcgacacgagtccagccactcgaacttagtttaaggt<br>cggcgagcaagaccaggtaacgagcaaccaatacatctgtcctttgacccggcatgtcctgc<br>tgtacaggtccgcattagatcagaagtgcggttccatgacgagccacgttccctacaacgaa<br>gcgtaaactagtacccttctacacaggcaccgcccggagtaggaaggattatgcttttgcct<br>ttaggaatttctagattctggtccgtgctgcggcctgcaacgtggacttacttataactgcg<br>gttaggacgattcatctgaaggaatacgctctttttcgactgcagctcgcgtgacgcttggc<br>tgaaaaattgaaactggagcttcctctacggatcaacgtttaactacccactgcctattcct<br>atgtactgatcgtcgagtcttgccaggattccgtcgcgggactcgtatcaacgttcagttga<br>gttgtgtcatgctaaccgcacattgtgagtcaccaagtgtcccatttggtcaactgatctcg<br>caaaaggtaagggccgtagcaaagtcgccagcttcgtcaattgatggcctatttttaactcg<br>ccgcttacggtcggaatctgaacggagacgtctttggcagaatggcgttacgcaccaatcta<br>taaaaagtttttgttggaaaggaggataatttctactggaccggtgttgcgacggaggagat<br>cgaattgctaatcaaccggtatgcacacttccattgctgtgcagttgccctaatcagctatc<br>catccaacataaaactgtctgagtgcttaaacggtcacaccaaatgattgtggtgccgtatc<br>tatagaatatccttagagcgtctgcttcctcgtcacacgagaccggttagtcccaagcacaa<br>taacgaatccagcttctgtttgccttagctccggtgatgcatgtttctgcttccggcggtgc<br>ggatgccacagctgccactgcaggtggaggagaagctgccaagtccaaaccaactacattta<br>ctccaccagattccacccatatcgacttacttccaatagctggttttcgcgttgacccacat<br>tacaatgtatctacgactcaagttattttaatgtatagcgttctgtattcgacccttccata<br>atgccctctatgtgaaactaacaacaatttgaccctcaaactttaagtataccagcttatgg<br>caacagtctgcgagccatggaagggatataacctgacgcagattattgcactgtccaagatc<br>cttcatgccacatcttcaggagggcgggtgattagcaccgtaaacagcgggttgatatcatc<br>aagcgaactgcagagaaatccgcgggaacactgggcttagcgcccatctcacccttaaaaat<br>taaacgcatctcccggtttcaggcattgctaccctgcgccggctagcgccttcccaatcctg<br>tggcttaagtctactgcgaaacaggttttataacagttccaccgcaatcaggtggccatttg<br>tcctcactctaatcccatccacccgttgatagtcaaagattcctctaataggcccatgaacg<br>tgcaaagttcccaatcgaacccacttggcacatacagtatccggcaaatgttatatcaacaa<br>gtcgcctgaacgtgccgcaacaacggatcaactgtagcttcgtgctgcctcagatgcatggc<br>tcgtgcccttctgttcgtgctgcatcggttgtatctcaatgactggactcagcatcgtagct<br>aagtaggcagggattttagtggtgtctaaagaataccggcggtaggcccaaaaaattacctc<br>gttcacccactaaagagatatcccgtacatctactatctactcaggaagatcaccactctag<br>cgtgggagccgctataatggatgcaggcagcccgggttagcgtgatgaaggacgttttaagt<br>tactactactggagttgcgggcgcaagacgatggctaagtaagagcccagagtttaggcctt<br>gtctaaaccgtaatgaaactgacatcggtagtcaatgtgtcgacgagttttgatttcagtat<br>atacgtactgttaaccgacgtctggatgtcagaaatttcgtgcatgtggcaggctcggttcg<br>aggaatctcgttccggaagttagggtatcggcgagggaactagtataaggactcgactgatg<br>catggctcagctaacagcgggcactctatgtcctaaggatagtaagaggagcaggacaacca<br>tccgggtgtaacgggttgatgcaagcgacatactaacaatgcctaggatagctgtgccatca<br>ggcggaaaagaatccaatatgatggtgctcaggactcattattacaatatagtacatttacc<br>agagaggtcccgcgggtcgccgaacacctacgcgaccctataagtttcttacactatcgata<br>atggagaaagcttatttgagggactacgatcttttacacccatggactttcagccgagatat<br>caaaatcgtagttatgttgtagcctgtagatttgtattcacgggctgtactttagccgagga<br>cagacccatatctgcttataataggtcatcatccctacattgtgtgagccagtctccaccgc |

| | |
|---|---|
| | tcgatgccaggcagtccaatgttatgtaaaacgaaggcgaaagggtgtctaacaccatctga<br>tcgatacaaactcgcactggccgcccacaaacgtcatggaaaggtagaaaattcgatgggct<br>gacgctattaccgtatcataggtcgactcacgtgggtacgtgccgtagctcccatgtttacc<br>gcttatgcgggtccagtggaaagagctttactggacgaataacgctgctgcttttaatcca<br>tatatgatactgcctagaacaagtacgggaaaatttcgacggcgtgcaatgtgcaatatgtt<br>tccgctatttgatcactcttggccgagtgcaatctctcactcgcgcttttgggctaacgaca<br>tagactcaatatcttagagtgagacgtgcggtctttcagtggagaaagccctgtttagacca<br>caggttcctattattcggatgaacgacctttaagataggtcaaccattatgacagttgcctg<br>agtaagaacacgagcggagtatctgattctgatgcttagacgctgtcgcatcccgtgaaagc<br>tcatccagaccgggtgagcgtagacctataactacgccaacctacccggccgggaaatcatg<br>taggcaactcaacccgctcgcatgtaagttgtccataatatgaatttacccatccgaattgt<br>atcgtggagttgttcggctagtggcaggagttcttagactaatgacaccctcactgttgcgg<br>cggtaacaaccattaactattacaagttgcggtttggtaaggttagggtaactgtagttaaa<br>agtatttctgcgattgctcgtttcgtcagatcacttcacagcgcagtctacggcactaggga<br>caagatttgtgtactgtggacccgtagccgagaaatccacggcattcatgagacgttactcg<br>ggaactattcagtcagtgtatgtagtcggcaaccggtagtggttccggaacaagcttttgaa<br>aatcagttaatgtggggctgagagcagcactgtgcaagactggaaaggtggtgaagcaatac<br>accatgaatcccggatcgctgacgcgccctgtagcggcgcattaagcgcggcgggtgtggtg<br>gttacgcgcagcgtgaccgctacacttgccagcgccctagcgcccgctcctttcgctttctt<br>cccttcctttctcgccacgttcgccggctttccccgtcaagctctaaatcgggggctccctt<br>tagggttccgatttagtgctttacggcacctcgaccccaaaaaacttgatttgggtgatggt<br>tcacgtagtgggccatcgccctgatagacggtttttcgccctttgacgttggagtccacgtt<br>cttt |
| 9072-nt ssDNA sequence for length test | aatagtggactcttgttccaaactggaacaacactcaaccctatctcgggctattcttttga<br>tttataagggattttgccgatttcggaacgggtacctacgaagagttccagcagggattcca<br>agaaatggccaatgaagattgtttcccgcacctgggctctttcgcgccaatcgacgttaaac<br>acttgaatttaagatgagcatcgcagaagtggatgatatgcgcgcctacggtccaatcggct<br>gggtgagcgtccctataatctcacggataccgacgaaagtagggcaataatcgcgctgcatg<br>agaacggatagcccggatgcaaaggtggaacgtaattgttaagagaagcaaatacgttaagc<br>tatcatttcccgtaaatcttttacatacgtaaggtggaagtctaaactagcctagctgcaca<br>agcatcggactgcttgccctctatctttgtggaagttcaaggtaatggaagcggtagaacgt<br>atgatcgtcacgcagacctaaagaatcacgccgcgcttacgtccaaccttgggacctacatc<br>tgtacagacccaggtccaggccgaggttatggatgtagatttgacttggctgatctacagtc<br>tcgagttacaatggcgtcactgctacctccatatgaaggataagaggcagccacaactcagc<br>caccatcctaccggaaacgctttaagggcgaacgtgaagggaagtgcctacacgaccctcta<br>tgtgggaagtagtctacgatacacgatgattggaagtgtggcggcataggatgcatctacgg<br>accgacactagttatagaagggtgcagtccttgacgcatacgaatagcgttagctcgcgatt<br>tcccgctgggttactcatggagagcggctcagacgcctagcccatactgaccgtgtaccaac<br>gatccaccaattattttcagaacagcccttttatggcgaagggaaaacagagccttagtatg<br>ttacagtccgctctattgggagatctcgggcaggcggaagatattaattagaaccgcagacc<br>gcatgatgtgccgtcctcaaatggcctggatatcccgtcacacgaattatctcagttgaagt<br>gatcctttagggcccgagagcaccacaagctcccaaccgctgcgaagagtgctaacgagttg<br>tcctatgttaagttattcgtatttaacccacgcccgcaccacctccgggaataatgtccaag<br>atcatatcaaggaaacccagggcacgataagcggcacacagacaaccactttggctagttaa<br>gataacggatactctgacgtacagccggccattcgttctacttttgtgtagtcagatggtcc<br>gtaacaatggcccggctctgacaacaagtataagactgcatgacgggtgggtccacgttaga<br>tgttcgtgaggtagcccatacatcattctttacgaacaagcctctcccgagccagcagagga<br>ggaatcagtgatcgttcacatggcgtacaacaaggcgtgcggcgccgtcggctagcgctgtg<br>aacgtcgcagcgttgattccaaccctctgcccacgacttcgccgctaactggaagcggtatt<br>aaacccgaaggttccacgggtggccacttggtgaaggcggcatcttaaagttctgacctcaa<br>ttactatggtgccgtcatctaaccaaccattaagaagtagaattgtaaccggttatcggaat<br>cagtgtcgacatactcaatgcccttaccctccgtaactctatgttttccgggttgttcatat<br>cggacatgctatcgcttagacgactcgtctatatagggcgttagtatatcatgggcgtacta<br>tcccggtcaatcgtctacgtcgcaaatcacctccctttgggcaggagagcacttctacactg<br>cctggcgctggcaccgcacaatcaataatgagattgtccggtgagacttacaatcccatcag<br>cgagtctgcaatcccgctgtcacgataacatagtactcgccttctggcggttcctcagctat<br>cctaacgtctatctaaactcaatgtggcgttatttggttcaagcaggcgggtcgtcaacgcg<br>aaacgtaagtcccacacggccgtaacgcaatgtggcaattccagagttaccgatcgaggcca |

| | |
|---|---|
| | atggcccggccaagccgcaaccgcatcctccgccggtaaaatgactaaatgagtggaaccgt<br>cgcgcttacgctctcggtgcccggaaaactctacagcatattgtctcattggtcctttggcg<br>tatacagacacttagactaactctgactgtcgagtaaattccaatgaacgccagcactcagt<br>acgcacggtgcgtccagggatggttacaaagctagagatgtatgtgctctcatcgaataacc<br>cgtagtatttggacctgaataccctggaatccgagggtgagaccattttacgttatcactgt<br>gacatgctttttccggtcaatacacgggaaactacggacgctatggtcgggataaacgccgc<br>atcatgtcaagctgaggccgctaaacaaagcatcaaggtagatttatttacaagcaaggcct<br>tgcgggcccagtccatgcgctgctgagattacagggattccgtgccagcccttcaacgtctc<br>aagacaactaacaggccttgaattcggccacactcaccggtcccacaatgtgccgggttcgc<br>atcaccgctgcttgggatagtatgcacacaaaagtagcttccacgagcggttgcccaattag<br>atggcgacccgcgtacaggctctaggaggctggaaagtccctctcaccgtcaaatatctgag<br>acgttatacccgacccatcatacgcgaataaagtactagtctcgcctgtcgccctcaggttt<br>accaccaataggacacggaaggcgctctgacaccagccaactagacagacatcggcgaggtg<br>tacttgacgcactttagaagctgctcccttgtgggaaccattgggcaaccgaacatagccgc<br>aatccagtcgcatcatcggtgggtcactgaccgaggattttggcggctcacgccttctgggc<br>tagcttcttctggcttgttagttattgcgtttacgttactctatatcccactaactatctat<br>atacctgtgctttcactacaatggctgcacagttatcttattttagcaaagctttgggttgc<br>ctggagtttcccaaagcgggacttgtaagcccgtcctatatcaggaggcggacgagagacgc<br>agcatgttatcttctaatctgcgaacggtacccggtctttcgtcggctaactagatctgtgt<br>cctaggtattcaaatcgggcgaagtcggtatcgaagaaaagccctttaacgtaaactgtatt<br>cgtcggccgccgttagctcaatgatataactctcatcgcgtgagtcatggcgccacatttta<br>taaaacacgctactatccaatcgaggagatcgctgcaccaataacagtctcaatccaacgaa<br>ctagtatttcatcaccgatccgcgaatcgtgagaccgacccatcgtcttatcgtcctaagca<br>gcacgggccacgacgtgcgaggcggcgtaacttgcgagttctacctatatacttgacaatct<br>cagctactccaaactgcccatggttggttggccatatgtacaccgagtcctagtacatcctc<br>actggacacgcgttcgcttgttggagagatgaaatccaagatattccttgtagggagctaat<br>cttgccaacactcaaattcctgatgcctcccaaaatacccgggctcaggtcaaaaaagccat<br>gaagcttcaagcccatgcttttcttgagtgattatcgctggccgggcgtataagttaatcca<br>gctaactggcgtgtcaacgaaagggtgggacacaatggttttccggctgtctcccagcaagt<br>gtcagaggcatttgcctttctctcaaccaaagcgctacactacacaggtcatcccgtgaaca<br>ttaggtagagttctccagtcagtcctcgacacgagtccagccactcgaacttagtttaaggt<br>cggcgagcaagaccaggtaacgagcaaccaatacatctgtcctttgacccggcatgtcctgc<br>tgtacaggtccgcattagatcagaagtgcggttccatgacgagccacgttccctacaacgaa<br>gcgtaaactagtacccttctacacaggcaccgcccggagtaggaaggattatgcttttgcct<br>ttaggaatttctagattctggtccgtgctgcggcctgcaacgtggacttacttataactgcg<br>gttaggacgattcatctgaaggaatacgctctttttcgactgcagctcgcgtgacgcttggc<br>tgaaaaattgaaactggagcttcctctacggatcaacgtttaactacccactgcctattcct<br>atgtactgatcgtcgagtcttgccaggattccgtcgcgggactcgtatcaacgttcagttga<br>gttgtgtcatgctaaccgcacattgtgagtcaccaagtgtcccatttggtcaactgatctcg<br>caaaaggtaagggccgtagcaaagtcgccagcttcgtcaattgatggcctattttttaactcg<br>ccgcttacggtcggaatctgaacggagacgtctttggcagaatggcgttacgcaccaatcta<br>taaaaagtttttgttggaaaggaggataatttctactggaccggtgttgcgacggaggagat<br>cgaattgctaatcaaccggtatgcacacttccattgctgtgcagttgccctaatcagctatc<br>catccaacataaaactgtctgagtgcttaaacggtcacaccaaatgattgtggtgccgtatc<br>tatagaatatccttagagcgtctgcttcctcgtcacacgagaccggttagtcccaagcacaa<br>taacgaatccagcttctgtttgccttagctccggtgatgcatgtttctgcttccggcggtgc<br>ggatgccacagctgccactgcaggtggaggagaagctgccaagtccaaaccaactacattta<br>ctccaccagattccacccatatcgacttacttccaatagctggttttcgcgttgacccacat<br>tacaatgtatctacgactcaagttattttaatgtatagcgttctgtattcgacccttccata<br>atgccctctatgtgaaactaacaacaatttgaccctcaaactttaagtataccagcttatgg<br>caacagtctgcgagccatggaagggatataacctgacgcagattattgcactgtccaagatc<br>cttcatgccacatcttcaggagggcgggtgattagcaccgtaaacagcgggttgatatcatc<br>aagcgaactgcagagaaatccgcgggaacactgggcttagcgcccatctcacccttaaaaat<br>taaacgcatctcccggtttcaggcattgctaccctgcgccggctagcgccttcccaatcctg<br>tggcttaagtctactgcgaaacaggttttataacagttccaccgcaatcaggtggccatttg<br>tcctcactctaatcccatccacccgttgatagtcaaagattcctctaataggcccatgaacg<br>tgcaaagttcccaatcgaacccacttggcacatacagtatccggcaaatgttatatcaacaa<br>gtcgcctgaacgtgccgcaacaacggatcaactgtagcttcgtgctgcctcagatgcatggc<br>tcgtgcccttctgttcgtgctgcatcggttgtatctcaatgactggactcagcatcgtagct |

| | |
|---|---|
| | aagtaggcagggattttagtggtgtctaaagaataccggcggtaggcccaaaaaattacctc<br>gttcacccactaaagagatatcccgtacatctactatctactcaggaagatcaccactctag<br>cgtgggagccgctataatggatgcaggcagcccgggttagcgtgatgaaggacgttttaagt<br>tactactactggagttgcgggcgcaagacgatggctaagtaagagcccagagtttaggcctt<br>gtctaaaccgtaatgaaactgacatcggtagtcaatgtgtcgacgagttttgatttcagtat<br>atacgtactgttaaccgacgtctggatgtcagaaatttcgtgcatgtggcaggctcggttcg<br>aggaatctcgttccggaagttagggtatcggcgagggaactagtataaggactcgactgatg<br>catggctcagctaacagcgggcactctatgtcctaaggatagtaagaggagcaggacaacca<br>tccgggtgtaacgggttgatgcaagcgacatactaacaatgcctaggatagctgtgccatca<br>ggcggaaaagaatccaatatgatggtgctcaggactcattattacaatatagtacatttacc<br>agagaggtcccgcgggtcgccgaacacctacgcgaccctataagtttcttacactatcgata<br>atggagaaagcttatttgagggactacgatcttttacacccatggactttcagccgagatat<br>caaaatcgtagttatgttgtagcctgtagatttgtattcacgggctgtactttagccgagga<br>cagacccatatctgcttataataggtcatcatccctacattgtgtgagccagtctccaccgc<br>tcgatgccaggcagtccaatgttatgtaaaacgaaggcgaaagggtgtctaacaccatctga<br>tcgatacaaactcgcactggccgcccacaaacgtcatggaaaggtagaaaattcgatgggct<br>gacgctattaccgtatcataggtcgactcacgtgggtacgtgccgtagctcccatgtttacc<br>gcttatgcgggtccagtggaaagagctttactggacgaataacgctgctgcttttttaatcca<br>tatatgatactgcctagaacaagtacgggaaaatttcgacggcgtgcaatgtgcaatatgtt<br>tccgctatttgatcactcttggccgagtgcaatctctcactcgcgcttttgggctaacgaca<br>tagactcaatatcttagagtgagacgtgcggtctttcagtggagaaagccctgtttagacca<br>caggttcctattattcggatgaacgacctttaagataggtcaaccattatgacagttgcctg<br>agtaagaacacgagcggagtatctgattctgatgcttagacgctgtcgcatcccgtgaaagc<br>tcatccagaccgggtgagcgtagacctataactacgccaacctacccggccgggaaatcatg<br>taggcaactcaacccgctcgcatgtaagttgtccataatatgaatttacccatccgaattgt<br>atcgtggagttgttcggctagtggcaggagttcttagactaatgacaccctcactgttgcgg<br>cggtaacaaccattaactattacaagttgcggtttggtaaggttagggtaactgtagttaaa<br>agtatttctgcgattgctcgtttcgtcagatcacttcacagcgcagtctacggcactaggga<br>caagatttgtgtactgtggacccgtagccgagaaatccacggcattcatgagacgttactcg<br>ggaactattcagtcagtgtatgtagtcggcaaccggtagtggttccggaacaagcttttgaa<br>aatcagttaatgtgggttgagctgctcagaggcgcccagttgtaccggaaaagtatgcatat<br>ctttaatggctatgcaagtatccaactatgctggcggcagggtagccatatccatcgggtcc<br>agaaggttcccattaaatacccgacactactagctatgttttgggtcctgttcccgagttac<br>ctgcataaggtgaataagccttagtaagttactgctctatgacattggctcagtcagactgt<br>tgcatacttatcttatagattcctatgcggaaaattccgaccgcagacttaataacttgatc<br>taatgcccggtttctcagtacgatactagttaaaccggccagcacaggttcgtatatatccc<br>tgtatggatatgatctcgggtcgactttaagcgactaagtgctctagcgcctgatggcgtgc<br>tttcttctcccacgttgctatatcaccgagtgagtacccgcggatagagttgcttcgcaggt<br>actcaaccacagttaggcaagtgcgaaggtattactgattaggcgtggccgcggccgtacac<br>ctcgttagtttgagggaagctgttccgatgactggctagcaggcctgggtgagtactagtga<br>tcagcaaagcctctctgtggtcgcataggtccgacaatatgagcgcagtatgcgagtcccac<br>cgaaattttgcaacgatcttgattctccctccagagtctaaatttcctatcctccaagtgta<br>actccagcatcagatgtttcttagaagtactcggagaatacaactttgagtacaatttgcgc<br>gtccgtccggtttcccggtgggatttaccaacttaaaacttctagaccattattcacagcct<br>agcgctgcgattggcggctcatgtgcggctcggctggtagggttggtgatgtggctaaccgt<br>cgaagcgttgtaggacaccttatttggagaagagacgttgtgacgggaaaagtgtctcatgc<br>gtaatgttccggagtatgggttcctgtgacgtgctgagagcagcactgtgcaagactggaaa<br>ggtggtgaagcaatacaccatgaatcccggatcgctgacgcgccctgtagcggcgcattaag<br>cgcggcgggtgtggtggttacgcgcagcgtgaccgctacacttgccagcgccctagcgcccg<br>ctcctttcgctttcttcccttcctttctcgccacgttcgccggctttccccgtcaagctcta<br>aatcgggggctccctttagggttccgatttagtgctttacggcacctcgaccccaaaaaact<br>tgatttgggtgatggttcacgtagtgggccatcgccctgatagacggtttttcgccctttga<br>cgttggagtccacgttcttt |
| 10,080-nt ssDNA sequence for length test | aatagtggactcttgttccaaactggaacaacactcaaccctatctcgggctattcttttga<br>tttataagggattttgccgatttcggaacgggtacctacgaagagttccagcagggattcca<br>agaaatggccaatgaagattgtttcccgcacctgggctctttcgcgccaatcgacgttaaac<br>acttgaatttaagatgagcatcgcagaagtggatgatatgcgcgcctacggtccaatcggct<br>gggtgagcgtccctataatctcacggataccgacgaaagtagggcaataatcgcgctgcatg |

| | |
|---|---|
| | agaacggatagcccggatgcaaaggtggaacgtaattgttaagagaagcaaatacgttaagc<br>tatcatttcccgtaaatcttttacatacgtaaggtggaagtctaaactagcctagctgcaca<br>agcatcggactgcttgccctctatctttgtggaagttcaaggtaatggaagcggtagaacgt<br>atgatcgtcacgcagacctaaagaatcacgccgcgcttacgtccaaccttgggacctacatc<br>tgtacagacccaggtccaggccgaggttatggatgtagatttgacttggctgatctacagtc<br>tcgagttacaatggcgtcactgctacctccatatgaaggataagaggcagccacaactcagc<br>caccatcctaccggaaacgctttaagggcgaacgtgaagggaagtgcctacacgaccctcta<br>tgtgggaagtagtctacgatacacgatgattggaagtgtggcggcataggatgcatctacgg<br>accgacactagttatagaagggtgcagtccttgacgcatacgaatagcgttagctcgcgatt<br>tcccgctgggttactcatggagagcggctcagacgcctagcccatactgaccgtgtaccaac<br>gatccaccaattattttcagaacagcccttttatggcgaagggaaaacagagccttagtatg<br>ttacagtccgctctattgggagatctcgggcaggcggaagatattaattagaaccgcagacc<br>gcatgatgtgccgtcctcaaatggcctggatatcccgtcacacgaattatctcagttgaagt<br>gatcctttagggcccgagagcaccacaagctcccaaccgctgcgaagagtgctaacgagttg<br>tcctatgttaagttattcgtatttaacccacgcccgcaccacctccgggaataatgtccaag<br>atcatatcaaggaaacccagggcacgataagcggcacacagacaaccactttggctagttaa<br>gataacggatactctgacgtacagccggccattcgttctacttttgtgtagtcagatggtcc<br>gtaacaatggcccggctctgacaacaagtataagactgcatgacgggtgggtccacgttaga<br>tgttcgtgaggtagcccatacatcattctttacgaacaagcctctcccgagccagcagagga<br>ggaatcagtgatcgttcacatggcgtacaacaaggcgtgcggcgccgtcggctagcgctgtg<br>aacgtcgcagcgttgattccaaccctctgcccacgacttcgccgctaactggaagcggtatt<br>aaacccgaaggttccacgggtggccacttggtgaaggcggcatcttaaagttctgacctcaa<br>ttactatggtgccgtcatctaaccaaccattaagaagtagaattgtaaccggttatcggaat<br>cagtgtcgacatactcaatgcccttaccctccgtaactctatgttttccgggttgttcatat<br>cggacatgctatcgcttagacgactcgtctatatagggcgttagtatatcatgggcgtacta<br>tcccggtcaatcgtctacgtcgcaaatcacctccctttgggcaggagagcacttctacactg<br>cctggcgctggcaccgcacaatcaataatgagattgtccggtgagacttacaatcccatcag<br>cgagtctgcaatcccgctgtcacgataacatagtactcgccttctggcggttcctcagctat<br>cctaacgtctatctaaactcaatgtggcgttatttggttcaagcaggcgggtcgtcaacgcg<br>aaacgtaagtcccacacggccgtaacgcaatgtggcaattccagagttaccgatcgaggcca<br>atggcccggccaagccgcaaccgcatcctccgccggtaaaatgactaaatgagtggaaccgt<br>cgcgcttacgctctcggtgcccggaaaactctacagcatattgtctcattggtcctttggcg<br>tatacagacacttagactaactctgactgtcgagtaaattccaatgaacgccagcactcagt<br>acgcacggtgcgtccagggatggttacaaagctagagatgtatgtgctctcatcgaataacc<br>cgtagtatttggacctgaataccctggaatccgagggtgagaccattttacgttatcactgt<br>gacatgctttttccggtcaatacacgggaaactacggacgctatggtcgggataaacgccgc<br>atcatgtcaagctgaggccgctaaacaaagcatcaaggtagatttatttacaagcaaggcct<br>tgcgggcccagtccatgcgctgctgagattacagggattccgtgccagcccttcaacgtctc<br>aagacaactaacaggccttgaattcggccacactcaccggtcccacaatgtgccgggttcgc<br>atcaccgctgcttgggatagtatgcacacaaaagtagcttccacgagcggttgcccaattag<br>atggcgacccgcgtacaggctctaggaggctggaaagtccctctcaccgtcaaatatctgag<br>acgttatacccgacccatcatacgcgaataaagtactagtctcgcctgtcgccctcaggttt<br>accaccaataggacacggaaggcgctctgacaccagccaactagacagacatcggcgaggtg<br>tacttgacgcactttagaagctgctcccttgtgggaaccattgggcaaccgaacatagccgc<br>aatccagtcgcatcatcggtgggtcactgaccgaggattttggcggctcacgccttctgggc<br>tagcttcttctggcttgttagttattgcgtttacgttactctatatcccactaactatctat<br>atacctgtgctttcactacaatggctgcacagttatcttattttagcaaagctttgggttgc<br>ctggagtttcccaaagcgggacttgtaagcccgtcctatatcaggaggcggacgagagacgc<br>agcatgttatcttctaatctgcgaacggtacccggtctttcgtcggctaactagatctgtgt<br>cctaggtattcaaatcgggcgaagtcggtatcgaagaaaagccctttaacgtaaactgtatt<br>cgtcggccgccgttagctcaatgatataactctcatcgcgtgagtcatggcgccacatttta<br>taaaacacgctactatccaatcgaggagatcgctgcaccaataacagtctcaatccaacgaa<br>ctagtatttcatcaccgatccgcgaatcgtgagaccgacccatcgtcttatcgtcctaagca<br>gcacgggccacgacgtgcgaggcggcgtaacttgcgagttctacctatatacttgacaatct<br>cagctactccaaactgcccatggttggttggccatatgtacaccgagtcctagtacatcctc<br>actggacacgcgttcgcttgttggagagatgaaatccaagatattccttgtagggagctaat<br>cttgccaacactcaaattcctgatgcctcccaaaatacccgggctcaggtcaaaaaagccat<br>gaagcttcaagcccatgcttttcttgagtgattatcgctggccgggcgtataagttaatcca<br>gctaactggcgtgtcaacgaaagggtgggacacaatggttttccggctgtctcccagcaagt |

gtcagaggcatttgcctttctctcaaccaaagcgctacactacacaggtcatcccgtgaaca
ttaggtagagttctccagtcagtcctcgacacgagtccagccactcgaacttagtttaaggt
cggcgagcaagaccaggtaacgagcaaccaatacatctgtcctttgacccggcatgtcctgc
tgtacaggtccgcattagatcagaagtgcggttccatgacgagccacgttccctacaacgaa
gcgtaaactagtacccttctacacaggcaccgcccggagtaggaaggattatgcttttgcct
ttaggaatttctagattctggtccgtgctgcggcctgcaacgtggacttacttataactgcg
gttaggacgattcatctgaaggaatacgctctttttcgactgcagctcgcgtgacgcttggc
tgaaaaattgaaactggagcttcctctacggatcaacgtttaactacccactgcctattcct
atgtactgatcgtcgagtcttgccaggattccgtcgcgggactcgtatcaacgttcagttga
gttgtgtcatgctaaccgcacattgtgagtcaccaagtgtcccatttggtcaactgatctcg
caaaaggtaagggccgtagcaaagtcgccagcttcgtcaattgatggcctatttttaactcg
ccgcttacggtcggaatctgaacggagacgtctttggcagaatggcgttacgcaccaatcta
taaaaagtttttgttggaaaggaggataatttctactggaccggtgttgcgacggaggagat
cgaattgctaatcaaccggtatgcacacttccattgctgtgcagttgccctaatcagctatc
catccaacataaaactgtctgagtgcttaaacggtcacaccaaatgattgtggtgccgtatc
tatagaatatccttagagcgtctgcttcctcgtcacacgagaccggttagtcccaagcacaa
taacgaatccagcttctgtttgccttagctccggtgatgcatgtttctgcttccggcggtgc
ggatgccacagctgccactgcaggtggaggagaagctgccaagtccaaaccaactacattta
ctccaccagattccacccatatcgacttacttccaatagctggttttcgcgttgacccacat
tacaatgtatctacgactcaagttattttaatgtatagcgttctgtattcgacccttccata
atgccctctatgtgaaactaacaacaatttgaccctcaaactttaagtataccagcttatgg
caacagtctgcgagccatggaagggatataacctgacgcagattattgcactgtccaagatc
cttcatgccacatcttcaggagggcgggtgattagcaccgtaaacagcgggttgatatcatc
aagcgaactgcagagaaatccgcgggaacactgggcttagcgcccatctcacccttaaaaat
taaacgcatctcccggtttcaggcattgctaccctgcgccggctagcgccttcccaatcctg
tggcttaagtctactgcgaaacaggttttataacagttccaccgcaatcaggtggccatttg
tcctcactctaatcccatccacccgttgatagtcaaagattcctctaataggcccatgaacg
tgcaaagttcccaatcgaacccacttggcacatacagtatccggcaaatgttatatcaacaa
gtcgcctgaacgtgccgcaacaacggatcaactgtagcttcgtgctgcctcagatgcatggc
tcgtgcccttctgttcgtgctgcatcggttgtatctcaatgactggactcagcatcgtagct
aagtaggcagggattttagtggtgtctaaagaataccggcggtaggcccaaaaaattacctc
gttcacccactaaagagatatcccgtacatctactatctactcaggaagatcaccactctag
cgtgggagccgctataatggatgcaggcagcccgggttagcgtgatgaaggacgttttaagt
tactactactggagttgcgggcgcaagacgatggctaagtaagagcccagagtttaggcctt
gtctaaaccgtaatgaaactgacatcggtagtcaatgtgtcgacgagttttgatttcagtat
atacgtactgttaaccgacgtctggatgtcagaaatttcgtgcatgtggcaggctcggttcg
aggaatctcgttccggaagttagggtatcggcgagggaactagtataaggactcgactgatg
catggctcagctaacagcgggcactctatgtcctaaggatagtaagaggagcaggacaacca
tccgggtgtaacgggttgatgcaagcgacatactaacaatgcctaggatagctgtgccatca
ggcggaaaagaatccaatatgatggtgctcaggactcattattacaatatagtacatttacc
agagaggtcccgcgggtcgccgaacacctacgcgaccctataagtttcttacactatcgata
atggagaaagcttatttgagggactacgatcttttacacccatggactttcagccgagatat
caaaatcgtagttatgttgtagcctgtagatttgtattcacgggctgtactttagccgagga
cagacccatatctgcttataataggtcatcatccctacattgtgtgagccagtctccaccgc
tcgatgccaggcagtccaatgttatgtaaaacgaaggcgaaagggtgtctaacaccatctga
tcgatacaaactcgcactggccgcccacaaacgtcatggaaaggtagaaaattcgatgggct
gacgctattaccgtatcataggtcgactcacgtgggtacgtgccgtagctcccatgtttacc
gcttatgcgggtccagtggaaagagctttactggacgaataacgctgctgcttttaatcca
tatatgatactgcctagaacaagtacgggaaaatttcgacggcgtgcaatgtgcaatatgtt
tccgctatttgatcactcttggccgagtgcaatctctcactcgcgcttttgggctaacgaca
tagactcaatatcttagagtgagacgtgcggtctttcagtggagaaagccctgtttagacca
caggttcctattattcggatgaacgacctttaagataggtcaaccattatgacagttgcctg
agtaagaacacgagcggagtatctgattctgatgcttagacgctgtcgcatcccgtgaaagc
tcatccagaccgggtgagcgtagacctataactacgccaacctacccggccgggaaatcatg
taggcaactcaacccgctcgcatgtaagttgtccataatatgaatttacccatccgaattgt
atcgtggagttgttcggctagtggcaggagttcttagactaatgacaccctcactgttgcgg
cggtaacaaccattaactattacaagttgcggtttggtaaggttagggtaactgtagttaaa
agtatttctgcgattgctcgtttcgtcagatcacttcacagcgcagtctacggcactaggga
caagatttgtgtactgtggacccgtagccgagaaatccacggcattcatgagacgttactcg

| | |
|---|---|
| | ggaactattcagtcagtgtatgtagtcggcaaccggtagtggttccggaacaagcttttgaa<br>aatcagttaatgtgggttgagctgctcagaggcgcccagttgtaccggaaaagtatgcatat<br>ctttaatggctatgcaagtatccaactatgctggcggcagggtagccatatccatcgggtcc<br>agaaggttcccattaaatacccgacactactagctatgttttgggtcctgttcccgagttac<br>ctgcataaggtgaataagccttagtaagttactgctctatgacattggctcagtcagactgt<br>tgcatacttatcttatagattcctatgcggaaaattccgaccgcagacttaataacttgatc<br>taatgcccggtttctcagtacgatactagttaaaccggccagcacaggttcgtatatatccc<br>tgtatggatatgatctcgggtcgactttaagcgactaagtgctctagcgcctgatggcgtgc<br>tttcttctcccacgttgctatatcaccgagtgagtacccgcggatagagttgcttcgcaggt<br>actcaaccacagttaggcaagtgcgaaggtattactgattaggcgtggccgcggccgtacac<br>ctcgttagtttgagggaagctgttccgatgactggctagcaggcctgggtgagtactagtga<br>tcagcaaagcctctctgtggtcgcataggtccgacaatatgagcgcagtatgcgagtcccac<br>cgaaattttgcaacgatcttgattctccctccagagtctaaatttcctatcctccaagtgta<br>actccagcatcagatgtttcttagaagtactcggagaatacaactttgagtacaatttgcgc<br>gtccgtccggtttcccggtgggatttaccaacttaaaacttctagaccattattcacagcct<br>agcgctgcgattggcggctcatgtgcggctcggctggtagggttggtgatgtggctaaccgt<br>cgaagcgttgtaggacaccttatttggagaagagacgttgtgacgggaaaagtgtctcatgc<br>gtaatgttccggagtatgggttcctgtgacgtgtagcagccttcaaatggacggatttacgc<br>caatatttcattaagtgagcctaaaattagcccgcaacgaggtgatctgggagactacaatc<br>taaggttgcgagtgtgatcagctcagaaacaatagctcgtgaacctcgattcagcagggatt<br>gcactctgtggtccgtttctccactctctggttggtttacgagtgagccctttaaccctggt<br>aggatccataaaacatgtaaaaccaccttgcttgtgctagcattagggaccggtgttcacac<br>ccatctatatggaggctcctggtgcacctcggacaaatgaggctttaagctatcgattccga<br>atggctacctggcgcatgcaaggtgccctgttcatgtccagtcagtgctgatgggaccgctg<br>cgagagacctagtagattcttcgcataaaaatatatgtgttaacacggtactgtccgtggta<br>tgccgttagaaataggtgcccatccgactcacgtcctgccagcgaccaatggaatcccgcta<br>aacagaaatgattatagtgagcaaggcatagtaaatatcagtccggtactactctacttcga<br>gtccctaaagaacgacaaggatctgtgaataaaactatagatcaattcagtgcttaccctcc<br>tatctgcttaaggtgtaattgtatgagggtgccgcgaccttcgtcatttcgaaaatgcccag<br>ttgacctaaaaccaggcaatgtctagaaccgcatagggacttaatgacaggccgccgatcca<br>gcaatgatttattcgccgaatagtcccaagtgttcacgggagatcgttattgggaggcgggt<br>gtatgtgtgagcgccctacacccttctatcttctcgtacggccttgttgagtatttatctgc<br>cgacttgatcaaaatagggtccgccatcgtcacattacgacttcgaaatcccgatgtgctgc<br>ttagaaattttgtaacatcggtaccaagagagtgcaaaacgaataattgctgagagcagcac<br>tgtgcaagactggaaaggtggtgaagcaatacaccatgaatcccggatcgctgacgcgccct<br>gtagcggcgcattaagcgcggcgggtgtggtggttacgcgcagcgtgaccgctacacttgcc<br>agcgccctagcgcccgctcctttcgctttcttcccttcctttctcgccacgttcgccggctt<br>tccccgtcaagctctaaatcgggggctccctttagggttccgatttagtgctttacggcacc<br>tcgaccccaaaaaacttgatttgggtgatggttcacgtagtgggccatcgccctgatagacg<br>gtttttcgccctttgacgttggagtccacgttcttt |
| 20,724-nt ssDNA sequence for length test | aatagtggactcttgttccaaactggaacaacactcaaccctatctcgggctattcttttga<br>tttataagggattttgccgatttcggaacgggtacctacgaagagttccagcagggattcca<br>agaaatggccaatgaagattgtttcccgcacctgggctctttcgcgccaatcgacgttaaac<br>acttgaatttaagatgagcatcgcagaagtggatgatatgcgcgcctacggtccaatcggct<br>gggtgagcgtccctataatctcacggataccgacgaaagtagggcaataatcgcgctgcatg<br>agaacggatagcccggatgcaaaggtggaacgtaattgttaagagaagcaaatacgttaagc<br>tatcatttcccgtaaatcttttacatacgtaaggtggaagtctaaactagcctagctgcaca<br>agcatcggactgcttgccctctatctttgtggaagttcaaggtaatggaagcggtagaacgt<br>atgatcgtcacgcagacctaaagaatcacgccgcgcttacgtccaaccttgggacctacatc<br>tgtacagacccaggtccaggccgaggttatggatgtagatttgacttggctgatctacagtc<br>tcgagttacaatggcgtcactgctacctccatatgaaggataagaggcagccacaactcagc<br>caccatcctaccggaaacgctttaagggcgaacgtgaagggaagtgcctacacgaccctcta<br>tgtgggaagtagtctacgatacacgatgattggaagtgtggcggcataggatgcatctacgg<br>accgacactagttatagaagggtgcagtccttgacgcatacgaatagcgttagctcgcgatt<br>tcccgctgggttactcatggagagcggctcagacgcctagcccatactgaccgtgtaccaac<br>gatccaccaattattttcagaacagcccttttatggcgaagggaaaacagagccttagtatg<br>ttacagtccgctctattgggagatctcgggcaggcggaagatattaattagaaccgcagacc<br>gcatgatgtgccgtcctcaaatggcctggatatcccgtcacacgaattatctcagttgaagt |

| | |
|---|---|
| | gatcctttagggcccgagagcaccacaagctcccaaccgctgcgaagagtgctaacgagttg<br>tcctatgttaagttattcgtatttaacccacgcccgcaccacctccgggaataatgtccaag<br>atcatatcaaggaaacccagggcacgataagcggcacacagacaaccactttggctagttaa<br>gataacggatactctgacgtacagccggccattcgttctacttttgtgtagtcagatggtcc<br>gtaacaatggcccggctctgacaacaagtataagactgcatgacgggtgggtccacgttaga<br>tgttcgtgaggtagcccatacatcattctttacgaacaagcctctcccgagccagcagagga<br>ggaatcagtgatcgttcacatggcgtacaacaaggcgtgcggcgccgtcggctagcgctgtg<br>aacgtcgcagcgttgattccaaccctctgcccacgacttcgccgctaactggaagcggtatt<br>aaacccgaaggttccacgggtggccacttggtgaaggcggcatcttaaagttctgacctcaa<br>ttactatggtgccgtcatctaaccaaccattaagaagtagaattgtaaccggttatcggaat<br>cagtgtcgacatactcaatgcccttaccctccgtaactctatgttttccgggttgttcatat<br>cggacatgctatcgcttagacgactcgtctatatagggcgttagtatatcatgggcgtacta<br>tcccggtcaatcgtctacgtcgcaaatcacctccctttgggcaggagagcacttctacactg<br>cctggcgctggcaccgcacaatcaataatgagattgtccggtgagacttacaatcccatcag<br>cgagtctgcaatcccgctgtcacgataacatagtactcgccttctggcggttcctcagctat<br>cctaacgtctatctaaactcaatgtggcgttatttggttcaagcaggcgggtcgtcaacgcg<br>aaacgtaagtcccacacggccgtaacgcaatgtggcaattccagagttaccgatcgaggcca<br>atggcccggccaagccgcaaccgcatcctccgccggtaaaatgactaaatgagtggaaccgt<br>cgcgcttacgctctcggtgcccggaaaactctacagcatattgtctcattggtcctttggcg<br>tatacagacacttagactaactctgactgtcgagtaaattccaatgaacgccagcactcagt<br>acgcacggtgcgtccagggatggttacaaagctagagatgtatgtgctctcatcgaataacc<br>cgtagtatttggacctgaataccctggaatccgagggtgagaccattttacgttatcactgt<br>gacatgctttttccggtcaatacacgggaaactacggacgctatggtcgggataaacgccgc<br>atcatgtcaagctgaggccgctaaacaaagcatcaaggtagatttatttacaagcaaggcct<br>tgcgggcccagtccatgcgctgctgagattacagggattccgtgccagcccttcaacgtctc<br>aagacaactaacaggccttgaattcggccacactcaccggtcccacaatgtgccgggttcgc<br>atcaccgctgcttgggatagtatgcacacaaaagtagcttccacgagcggttgcccaattag<br>atggcgacccgcgtacaggctctaggaggctggaaagtccctctcaccgtcaaatatctgag<br>acgttatacccgacccatcatacgcgaataaagtactagtctcgcctgtcgccctcaggttt<br>accaccaataggacacggaaggcgctctgacaccagccaactagacagacatcggcgaggtg<br>tacttgacgcactttagaagctgctcccttgtgggaaccattgggcaaccgaacatagccgc<br>aatccagtcgcatcatcggtgggtcactgaccgaggattttggcggctcacgccttctgggc<br>tagcttcttctggcttgttagttattgcgtttacgttactctatatcccactaactatctat<br>atacctgtgctttcactacaatggctgcacagttatcttattttagcaaagctttgggttgc<br>ctggagtttcccaaagcgggacttgtaagcccgtcctatatcaggaggcggacgagagacgc<br>agcatgttatcttctaatctgcgaacggtacccggtctttcgtcggctaactagatctgtgt<br>cctaggtattcaaatcgggcgaagtcggtatcgaagaaaagccctttaacgtaaactgtatt<br>cgtcggccgccgttagctcaatgatataactctcatcgcgtgagtcatggcgccacattta<br>taaaacacgctactatccaatcgaggagatcgctgcaccaataacagtctcaatccaacgaa<br>ctagtatttcatcaccgatccgcgaatcgtgagaccgacccatcgtcttatcgtcctaagca<br>gcacgggccacgacgtgcgaggcggcgtaacttgcgagttctacctatatacttgacaatct<br>cagctactccaaactgcccatggttggttggccatatgtacaccgagtcctagtacatcctc<br>actggacacgcgttcgcttgttggagagatgaaatccaagatattccttgtagggagctaat<br>cttgccaacactcaaattcctgatgcctcccaaaatacccgggctcaggtcaaaaaagccat<br>gaagcttcaagcccatgcttttcttgagtgattatcgctggccgggcgtataagttaatcca<br>gctaactggcgtgtcaacgaaagggtgggacacaatggttttccggctgtctcccagcaagt<br>gtcagaggcatttgcctttctctcaaccaaagcgctacactacacaggtcatcccgtgaaca<br>ttaggtagagttctccagtcagtcctcgacacgagtccagccactcgaacttagtttaaggt<br>cggcgagcaagaccaggtaacgagcaaccaatacatctgtcctttgacccggcatgtcctgc<br>tgtacaggtccgcattagatcagaagtgcggttccatgacgagccacgttccctacaacgaa<br>gcgtaaactagtacccttctacacaggcaccgcccggagtaggaaggattatgcttttgcct<br>ttaggaatttctagattctggtccgtgctgcggcctgcaacgtggacttacttataactgcg<br>gttaggacgattcatctgaaggaatacgctctttttcgactgcagctcgcgtgacgcttggc<br>tgaaaaattgaaactggagcttcctctacggatcaacgtttaactacccactgcctattcct<br>atgtactgatcgtcgagtcttgccaggattccgtcgcgggactcgtatcaacgttcagttga<br>gttgtgtcatgctaaccgcacattgtgagtcaccaagtgtcccatttggtcaactgatctcg<br>caaaaggtaagggccgtagcaaagtcgccagcttcgtcaattgatggcctatttttaactcg<br>ccgcttacggtcggaatctgaacggagacgtctttggcagaatggcgttacgcaccaatcta<br>taaaaagtttttgttggaaaggaggataatttctactggaccggtgttgcgacggaggagat |

| | |
|---|---|
| | cgaattgctaatcaaccggtatgcacacttccattgctgtgcagttgccctaatcagctatc<br>catccaacataaaactgtctgagtgcttaaacggtcacaccaaatgattgtggtgccgtatc<br>tatagaatatccttagagcgtctgcttcctcgtcacacgagaccggttagtcccaagcacaa<br>taacgaatccagcttctgtttgccttagctccggtgatgcatgtttctgcttccggcggtgc<br>ggatgccacagctgccactgcaggtggaggagaagctgccaagtccaaaccaactacattta<br>ctccaccagattccacccatatcgacttacttccaatagctggttttcgcgttgacccacat<br>tacaatgtatctacgactcaagttattttaatgtatagcgttctgtattcgacccttccata<br>atgccctctatgtgaaactaacaacaatttgaccctcaaactttaagtataccagcttatgg<br>caacagtctgcgagccatggaagggatataacctgacgcagattattgcactgtccaagatc<br>cttcatgccacatcttcaggagggcgggtgattagcaccgtaaacagcgggttgatatcatc<br>aagcgaactgcagagaaatccgcgggaacactgggcttagcgcccatctcacccttaaaaat<br>taaacgcatctcccggtttcaggcattgctaccctgcgccggctagcgccttcccaatcctg<br>tggcttaagtctactgcgaaacaggttttataacagttccaccgcaatcaggtggccatttg<br>tcctcactctaatcccatccacccgttgatagtcaaagattcctctaataggcccatgaacg<br>tgcaaagttcccaatcgaacccacttggcacatacagtatccggcaaatgttatatcaacaa<br>gtcgcctgaacgtgccgcaacaacggatcaactgtagcttcgtgctgcctcagatgcatggc<br>tcgtgcccttctgttcgtgctgcatcggttgtatctcaatgactggactcagcatcgtagct<br>aagtaggcagggattttagtggtgtctaaagaataccggcggtaggcccaaaaaattacctc<br>gttcacccactaaagagatatcccgtacatctactatctactcaggaagatcaccactctag<br>cgtgggagccgctataatggatgcaggcagcccgggttagcgtgatgaaggacgttttaagt<br>tactactactggagttgcgggcgcaagacgatggctaagtaagagcccagagtttaggcctt<br>gtctaaaccgtaatgaaactgacatcggtagtcaatgtgtcgacgagttttgatttcagtat<br>atacgtactgttaaccgacgtctggatgtcagaaatttcgtgcatgtggcaggctcggttcg<br>aggaatctcgttccggaagttagggtatcggcgagggaactagtataaggactcgactgatg<br>catggctcagctaacagcgggcactctatgtcctaaggatagtaagaggagcaggacaacca<br>tccgggtgtaacgggttgatgcaagcgacatactaacaatgcctaggatagctgtgccatca<br>ggcggaaaagaatccaatatgatggtgctcaggactcattattacaatatagtacatttacc<br>agagaggtcccgcgggtcgccgaacacctacgcgaccctataagtttcttacactatcgata<br>atggagaaagcttatttgagggactacgatcttttacacccatggactttcagccgagatat<br>caaaatcgtagttatgttgtagcctgtagatttgtattcacgggctgtactttagccgagga<br>cagacccatatctgcttataataggtcatcatccctacattgtgtgagccagtctccaccgc<br>gaaagtgaaacgtgatttcatgcgtcattttgaacattttgtaaatcttatttaataatgtg<br>tgcggcaattcacatttaatttatgaatgttttcttaacatcgcggcaactcaagaaacggc<br>aggttcggatcttagctactagagaaagaggagaaatactagaatgatggcttcctccgagg<br>atgttatcaaagagttcatgcgtttcaaagttcgtatggaaggttccgttaacggtcacgag<br>ttcgaaatcgaaggtgaaggtgaaggtcgtccgtacgaaggtacccagaccgctaaactgaa<br>agttaccaaaggtggtccgctgccgttcgcttgggacatcctgtccccgcagttccagtacg<br>gttccaaagcttacgttaaacacccggctgacatcccggactacctgaaactgtccttcccg<br>gaaggtttcaaatgggaacgtgttatgaacttcgaagatggtggtgttgttaccgttaccca<br>ggactcctccctgcaagacggtgagttcatctacaaagttaaactgcgtggtaccaacttcc<br>cgtccgacggtccggttatgcagaaaaaaaccatgggttgggaagcttccaccgaacgtatg<br>tacccggaggatggtgctctgaaaggtgaaatcaaaatgcgtctgaaactgaaagacggtgg<br>tcactacgacgctgaagttaaaaccacctacatggctaaaaaaccggttcagctgccgggtg<br>cttacaaaaccgacatcaaactggacatcacctcccacaacgaggactacaccatcgttgaa<br>cagtacgaacgtgctgaaggtcgtcactccaccggtgcttaataaaggtccaggcatcaaat<br>aaaacgaaaggctcagtcgaaagactgggcctttcgttttatctgttgtttgtcggtgaacg<br>ctctctactagagtcacactggctcaccttcgggtgggcctttctgcgtttataggttctGA<br>CGATTTGGAAGTGACACGCAAGAAGCTGGTCGATGACTGTCACCACTTCCGCCTCGAAGAAC<br>CTAACTTCTCTCTCGCCTCCAGCATCAGCAAGGATATTGAGTCCTGTGCTCAGATTTGGGCC<br>TTCTACGAAGAGTTCCAGCAGGGATTCCAAGAAATGGCCAATGAAGATTGGATCACCTTTCG<br>CACTAAGACCTACTTGTTTGAGGAGTTTCTGATGAATTGGCACGACCGCCTCAGGAAAGTGG<br>AGGAGCATTCTGTGATGACTGTCAAGCTCCAATCTGAGGTGGACAAATATAAGATTGTTATC<br>CCTATCCTGAAGTACGTCCGCGGAGAACACCTGTCACCCGATCACTGGCTGGATCTGTTCCG<br>CTTGCTGGGTCTGCCTCGCGGCACATCTCTGGAGAAACTGCTGTTCGGTGACCTGCTGAGAG<br>TTGCCGATACCATCGTGGCCAAGGCTGCTGACCTGAAAGATCTGAACTCACGCGCCCAGGGT<br>GAAGTGACCATCCGCGAAGCACTCAGGGAACTGGATTTGTGGGGCGTGGGTGCTGTGTTCAC<br>ACTGATCGACTATGAGGACTCCCAGAGCCGCACCATGAAGCTGATCAAGGATTGGAAGGACA<br>TCGTCAACCAGGTGGGCGACAATAGATGCCTCCTGCAGTCCTTGAAGGACTCACCATACTAT<br>AAAGGCTTTGAAGACAAGGTCAGCATCTGGGAAAGGAAACTCGCCGAACTGGACGAATATTT |

| | |
|---|---|
| | GCAGAACCTCAACCATATTCAGAGAAAGTGGGTTTACCTCGAACCAATCTTTGGTCGCGGAG<br>CCCTGCCCAAAGAGCAGACCAGATTCAACAGGGTGGATGAAGATTTCCGCAGCATCATGACA<br>GATATCAAGAAGGACAATCGCGTCACAACCTTGACTACCCACGCAGGCATTCGCAACTCACT<br>GCTGACCATCCTGGACCAATTGCAGAGATGCCAGCGCAGCCTCAACGAGTTCCTGGAGGAGA<br>AGCGCAGCGCCTTCCCTCGCTTCTACTTCATCGGAGACGATGACCTGCTGGAGATCTTGGGC<br>CAGTCAACCAATCCATCCGTGATTCAGTCTCACCTCAAGAAGCTGTTTGCTGGTATCAACTC<br>TGTCTGTTTCGATGAGAAGTCTAAGCACATTACTGCAATGAAGTCCTTGGAGGGAGAAGTTG<br>TGCCATTCAAGAATAAGGTGCCCTTGTCCAATAACGTCGAAACCTGGCTGAACGATCTGGCC<br>CTGGAGATGAAGAAGACCCTGGAGCAGCTGCTGAAGGAGTGCGTGACAACCGGACGCAGCTC<br>TCAGGGAGCTGTGGACCCTTCTCTGTTCCCATCACAGATCCTGTGCTTGGCCGAACAGATCA<br>AGTTTACCGAAGATGTGGAGAACGCAATTAAAGATCACTCCCTGCACCAGATTGAGACACAG<br>CTGGTGAACAAATTGGAGCAGTATACTAACATCGACACATCTTCCGAGGACCCAGGTAACAC<br>AGAGTCCGGTATTCTGGAGCTGAAACTGAAAGCACTGATTCTCGACATTATCCATAACATCG<br>ACGTGGTCAAGCAGCTGAACCAAATCCAAGTGCACACCACCGAAGATTGGGCCTGGAAGAAG<br>CAGTTGAGGTTCTACATGAAGTCCGACCACACCTGTTGCGTTCAGATGGTTGACAGCGAGTT<br>CCAGTACACCTATGAGTACCAAGGAAATGCCAGCAAGCTCGTTTACACTCCACTCACTGACA<br>AGTGTTACCTCACCTTGACACAGGCTATGAAGATGGGCCTGGGAGGCAACCCATACGGTCCA<br>GCTGGCACTGGTAAGACAGAGAGCGTTAAGGCACTCGGAGGTCTGCTGGGCAGGCAGGTCCT<br>CGTGTTCAACTGTGATGAAGGAATCGACGTTAAGTCCATGGGAAGAATCTTTGTTGGCCTCG<br>TTAAGTGTGGAGCTTGGGGTTGCTTCGACGAGTTCAACAGGCTGGAGGAATCTGTGCTGAGC<br>GCCGTCTCTATGCAGATCCAGACCATCCAGGACGCATTGAAGAACCACAGGACCGTCTGCGA<br>GCTGTTGGGTAAGGAAGTGGAGGTGAACTCCAACTCCGGAATCTTCATCACAATGAATCCCG<br>CAGGTAAAGGATATGGAGGAAGACAGAAACTCCCAGACAACCTGAAGCAGCTGTTCCGCCCA<br>GTGGCTATGTCCCATCCAGACAATGAGCTGATCGCCGAAGTCATCCTCTATTCCGAGGGATT<br>CAAAGATGCTAAAGTTCTCTCCAGAAAGCTCGTGGCCATCTTCAATCTGTCAAGAGAACTCC<br>TGACACCTCAGCAGCATTACGACTGGGGTCTGAGAGCCCTCAAGACCGTCCTGAGAGGTTCA<br>GGAAATCTCCTCAGGCAGCTGAACAAGAGCGGTACAACACAGAATGCAAATGAGAGCCACAT<br>TGTCGTCCAGGCTCTGAGGCTGAATACCATGTCAAAGTTCACATTCACAGACTGCACAAGAT<br>TTGACGCTCTGATTAAAGATGTGTTCCCTGGTATTGAACTCAAAGAAGTGGAGTATGACGAG<br>CTGAGCGCCGCTTTGAAGCAGGTGTTTGAGGAGGCTAACTATGAGATTATCCCTAATCAGAT<br>CAAGAAAGCATTGGAACTGTATGAACAGCTGTGTCAGAGGATGGGAGTGGTGATTGTGGGCC<br>CATCAGGCGCAGGTAAGAGCACTCTCTGGAGAATGCTGAGAGCAGCACTGTGCAAGACTGGA<br>AAGGTGGTGAAGCAATACACCATGAATCCCAAGGCCATGCCCAGGTACCAACTGCTGGGCCA<br>TATCGACATGGACACCAGAGAATGGAGCGACGGCGTGCTCACAAACTCCGCCAGACAAGTCG<br>TGCGCGAACCTCAAGACGTCAGCTCTTGGATCATCTGCGATGGTGATATTGACCCTGAGTGG<br>ATCGAGTCCCTGAATTCCGTGTTGGATGACAACAGGCTCCTCACAATGCCTTCTGGTGAGAG<br>AATCCAGTTCGGTCCTAACGTGAACTTCGTGTTCGAGACACACGATCTCAGCTGTGCTAGCC<br>CAGCTACTATCTCCCGCATGGGAATGATCTTCTTGTCCGACGAGGAGACAGATTTGAACTCA<br>TTGATCAAGTCTTGGCTCAGAAACCAGCCTGCAGAATATAGGAATAACCTGGAGAACTGGAT<br>CGGTGATTACTTCGAGAAGGCTTTGCAGTGGGTGCTGAAACAGAACGACTATGTCGTCGAAA<br>CCAGCCTGGTCGGTACAGTTATGAACGGACTCTCCCATCTGCACGGATGCAGAGATCACGAT<br>GAGTTTATCATCAATTTGATCCGCGGACTGGGAGGTAACTTGAATATGAAATCTCGCCTGGA<br>GTTCACTAAAGAAGTGTTCCACTGGGCTAGGGAGTCACCACCTGACTTCCACAAACCTATGG<br>ACACCTACTATGATTCCACAAGAGGCAGGTTGGCCACCTACGTGCTGAAGAAGCCTGAGGAC<br>CTCACCGCTGACGACTTCTCCAACGGACTGACTCTGCCCGTGATCCAGACTCCAGACATGCA<br>GCGCGGACTCGATTACTTTAAGCCCTGGCTCAGCTCCGATACCAAGCAACCTTTCATTCTCG<br>TGGGACCAGAGGGATGTGGTAAAGGAATGCTCCTGAGGTATGCATTCTCCCAGCTCCGCTCA<br>ACCCAAATTGCCACTGTTCACTGTTCAGCCCAAACAACTTCAAGGCATCTCCTCCAGAAGCT<br>CAGCCAGACCTGTATGGTTATCAGCACCAACACCGGCAGAGTTTACCGCCCAAAGGATTGTG<br>AGCGCCTCGTGTTGTACCTCAAAGATATCAATCTGCCCAAACTCGATAAGTGGGGCACTTCC<br>ACCCTCGTGGCATTTCTCCAACAGGTGCTGACCTACCAGGGCTTCTACGACGAGAACCTGGA<br>GTGGGTCGGATTGGAGAACATCCAGATTGTGGCTTCCATGTCTGCCGGCGGTAGGTTGGGAA<br>GGCATAAGCTCACCACCAGGTTTACATCAATTGTGAGACTGTGCTCAATCGATTATCCCGAG<br>CGCGAACAGTTGCAGACCATCTATGGCGCCTACCTCGAACCTGTCCTCCACAAGAATTTGAA<br>GAACCATAGCATCTGGGGCTCATCAAGCAAGATCTACCTCTTGGCTGGCTCTATGGTTCAGG<br>TGTACGAACAAGTCCGCGCCAAGTTCACCGTCGATGATTACTCACATTACTTCTTCACACCC<br>TGCATTCTGACACAATGGGTTCTGGGACTGTTTCGCTATGACCTGGAGGGAGGCTCATCAAA<br>CCACCCATTGGATTATGTCCTCGAAATCGTGGCTTACGAAGCCCGCAGGCTCTTTAGAGATA<br>AGATTGTTGGCGCTAAGGAACTGCATCTGTTCGATATTATCCTCACCTCTGTGTTTCAAGGT |

| | |
|---|---|
| | GATTGGGGCTCTGATATCCTGGATAATATGTCCGATTCCTTCTATGTGACATGGGGAGCCAG<br>GCACAACTCCGGTGCAAGGGCTGCTCCAGGCCAACCATTGCCACCTCATGGCAAGCCTCTCG<br>GCAAACTGAACTCAACTGACTTGAAGGACGTGATCAAGAAGGGCCTGATTCACTACGGCCGC<br>GACAACCAGAACCTGGACATTCTGTTGTTCCACGAGGTCCTGGAGTATATGTCCAGAATTGA<br>TCGCGTCCTGTCTTTCCCTGGTGGATCACTGCTGCTGGCCGGACGCTCTGGAGTTGGAAGAC<br>GCACTATTACTTCTCTGGTCAGCCATATGCATGGAGCCGTCCTGTTCTCTCCAAAGATCAGC<br>AGAGGCTACGAACTCAAGCAATTCAAGAACGATCTGAAACACGTCTTGCAACTCGCCGGTAT<br>CGAGGCCCAGCAGGTCGTCCTCCTCTTGGAGGACTATCAATTCGTCCACCCAACCTTCCTGG<br>AGATGATCAACTCCCTGCTGTCCTCTGGCGAGGTGCCCGGCTTGTACACTTTGGAGGAACTG<br>GAGCCACTGCTCCTCCCATTGAAGGATCAGGCATCACAGGACGGCTTCTTCGGCCCAGTGTT<br>CAATTACTTCACCTATCGCATTCAACAGAATCTCCACATTGTGCTGATTATGGACAGCGCTA<br>ATTCCAATTTCATGATCAATTGCGAGAGCAATCCCGCCCTCCATAAGAAGTGCCAGGTCCTC<br>TGGATGGAGGGTTGGTCTAATTCTTCTATGAAGAAGATTCCCGAGATGTTGTTTAGCGAGAC<br>TGGAGGCGGTGAGAAGTACAACGACAAGAAGCGCAAAGAGGAGAAGAAGAAGAACTCCGTCG<br>ATCCTGATTTCCTCAAGAGCTTCCTGCTGATCCACGAGTCTTGCAAAGCTTACGGAGCTACT<br>CCTAGCCAGTACATGACCTTCCTCCACGTCTATTCCGCCATCTCCAGCTCAAAGAAGAAGGA<br>GCTGCTGAAGCGCCAATCTCATCTGCAGGCCGGAGTCAGCAAGCTGAACGAAGCCAAAGCTC<br>TGGTGGATGAACTGAATCGCAAGGCTGGCGAACAATCAGTCCTCCTGAAGACTAAGCAGGAT<br>GAAGCTGACGCTGCCCTGCAAATGATTACCGTGTCTATGCAGGATGCTTCCGAGCAGAAGAC<br>AGAGCTGGAGAGGCTGAAGCACCGCATCGCAGAGGAGGTGGTCAAGATCGAGGAGAGAAAGA<br>ACAAGATTGACGACGAACTCAAAGAGGTGCAGCCTCTGGTGAACGAGGCCAAGCTCGCCGTG<br>GGTAATATCAAACCAGAGTCTCTCTCAGAGATCAGGTCACTGAGAATGCCACCAGACGTTAT<br>CCGCGACATCCTGGAGGGCGTCCTGCGCTTGATGGGTATCTTTGACACCTCTTGGGTGTCTA<br>TGAAGTCTTTCTTGGCCAAGCGCGGTGTCAGGGAGGACATTGCTACTTTCGACGCTAGGAAC<br>ATCTCCAAGGAAATTAGGGAATCTGTGGAGGAACTGCTGTTCAAGAATAAAGGTTCATTTGA<br>TCCCAAGAACGCTAAGAGAGCATCAACTGCAGCTGCACCCTTGGCAGCCTGGGTTAAGGCCA<br>ATATCCAGTACTCTCACGTGCTCGAACGCATCCACCCTCTGGAGACTGAGCAAGCCGGCCTG<br>GAAAGCAACCTGAAGAAGACCGAGGATAGAAAGAGGAAACTGGAAGAACTCCTCAATTCTGT<br>CGGTCAGAAGGTGTCAGAACTGAAGGAGAAATTTCAGAGCAGGACCTCAGAAGCAGCTAAGT<br>TGGAAGCTGAGGTGTCCAAGGCTCAGGAGACTATCAAAGCAGCTGAAGTGTTGATTAATCAG<br>CTGGACCGCGAACACAAGAGATGGAATGCTCAGGTCGTCGAGATTACTGAGGAGCTGGCAAC<br>CCTCCCAAAGAGGGCTCAGTTGGCCGCAGCCTTTATCACCTACTTGTCCGCTGCACCTGAGT<br>CTCTCAGAAAGACATGTCTGGAGGAGTGGACCAAGTCTGCCGGCCTGGAGAAGTTTGACCTG<br>AGAAGATTTCTCTGCACCGAGTCAGAGCAGCTGATCTGGAAGTCTGAAGGTCTGCCCAGCGA<br>TGACCTCTCAATCGAGAACGCACTGGTTATCTTGCAATCCCGCGTTTGCCCTTTCCTCATCG<br>ATCCCAGCTCACAGGCTACTGAGTGGCTGAAGACTCACTTGAAGGATTCCAGGCTGGAAGTG<br>ATCAACCAGCAAGACTCCAACTTCATCACTGCCCTGGAACTCGCCGTCCGCTTCGGCAAGAC<br>CTTGATCATCCAGGAGATGGATGGCGTGGAGCCAGTTCTGTACCCTCTGTTGCGCAGAGATT<br>TGGTGGCCCAAGGCCCACGCTATGTGGTCCAGATTGGAGATAAGATCATCGACTATAACGAG<br>GAGTTTCGCCTGTTCCTGAGCACTAGGAATCCCAACCCATTCATTCCACCAGACGCTGCCAG<br>CATCGTCACAGAGGTTAATTTCACAACCACCCGCTCAGGACTGAGAGGCCAGCTCCTGGCCC<br>TCACCATCCAACATGAGAAACCCGATTTGGAAGAACAGAAGACTAAGCTGCTCCAACAGGAG<br>GAGGATAAGAAGATCCAACTGGCCAAACTCGAAGAATCTCTGCTGGAGACATTGGCTACTTC<br>TCAGGGCAACATCCTGGAGAACAAGGACCTGATCGAGTCTCTGAATCAGACTAAGCATCTT<br>CCGCCCTCATCCAAGAGTCTCTGAAGGAATCTTATAAGCTGCAAATCTCTCTCGACCAGGAG<br>CGCGACGCATATCTGCCACTCGCTGAGTCTGCTTCAAAGATGTACTTCATTATCTCTGACCT<br>CTCCAAGATCAATAATATGTACAGGTTCTCCCTGGCCGCCTTTCTGAGGTTGTTTCAAAGGG<br>CACTCCAGAATAAGCAGGATTCTGAGAACACAGAACAGAGGATTCAATCCCTGATCAGCTCC<br>CTGCAGCACATGGTGTACGAATACATCTGCCGCTGCCTGTTCAAGGCCGACCAGCTGATGTT<br>CGCCCTGCATTTCGTCAGAGGTATGCATCCAGAATTGTTCCAGGAGAACGAATGGGATACCT<br>TCACTGGCGTGGTGGTCGGAGACATGTTGCGCAAAGCCGACTCCCAGCAGAAGATCAGGGAC<br>CAGTTGCCTTCATGGATCGATCAAGAGAGATCCTGGGCTGTGGCAACCTTGAAGATCGCTCT<br>GCCTTCCCTGTACCAGACTCTGTGTTTCGAGGACGCCGCTTTGTGGCGCACCTACTACAACA<br>ACAGCATGTGTGAGCAGGAATTTCCTAGCATCTTGGCTAAGAAGGTCAGCTTGTTCCAGCAG<br>ATCCTCGTCGTGCAGGTGCTCAGGCCAGATAGATTGCAGTCAGCTATGGCCCTCTTTGCCTG<br>CAAGACCTTGGGTTTGAAGGAAGTCTCTCCACTCCCACTCAATCTCAAGAGGCTGTACAAGG<br>AGACACTCGAAATCGAGCCCATTCTGATCATTATTTCTCCAGGAGCCGATCCCTCCCAGGAA<br>CTCCAGGAGCTCGCCAACGCCGAAAGATCAGGAGAATGTTACCACCAGGTTGCAATGGGCCA<br>GGGCCAAGCTGACTTGGCTATCCAAATGCTCAAGGAATGCGCAAGGAATGGTGACTGGCTGT |

GTTTGAAGAACTTGCACCTGGTTGTCTCCTGGCTGCCTGTCTTGGAGAAGGAACTGAACACC
TTGCAACCCAAAGACACTTTCAGGCTCTGGTTGACAGCCGAAGTGCACCCAAACTTTACACC
TATCCTCCTCCAGTCAAGCCTCAAGATCACTTATGAATCACCACCTGGACTCAAGAAGAATC
TCATGCGCACATATGAATCTTGGACACCTGAACAAATCTCTAAGAAAGATAACACTCACCGC
GCACATGCTCTGTTCTCTCTGGCCTGGTTTCACGCCGCATGCCAAGAGAGGCGCAACTACAT
TCCTCAAGGTTGGACCAAATTCTACGAGTTCTCTCTCTCCGACCTGAGGGCCGGTTACAATA
TCATTGACAGGCTCTTTGATGGTGCCAAGGACGTGCAATGGGAGTTTGTCCATGGTCTGCTG
GAGAACGCCATCTACGGAGGCCGCATCGATAACTATTTCGACTTGCGCGTCCTGCAGAGCTA
TTTGAAGCAATTCTTCAACTCTAGCGTCATTGACGTGTTCAATCAGCGCAACAAGAAGTCCA
TCTTTCCCTACAGCGTGAGCCTGCCACAAAGCTGCAGCATTCTGGATTACAGGGCTGTCATT
GAGAAGATTCCAGAAGACGATAAGCCATCCTTCTTCGGTCTGCCTGCCAACATTGCACGCTC
ATCACAGCGCATGATCTCATCTCAGGTGATTTCCCAACTCCGCATCCTGGGCCGCTCTATTA
CTGCAGGCTCAAAGTTCGATCGCGAGATCTGGAGCAATGAACTCAGCCCAGTGCTCAACCTG
TGGAAGAAACTGAACCAGAACTCCAACCTGATCCACCAGAAGGTCCCTCCTCCCAATGACCG
CCAAGGATCACCAATTCTGTCCTTTATTATCTTGGAGCAGTTCAACGCCATCAGATTGGTCC
AGTCAGTTCATCAATCACTGGCAGCCTTGAGCAAAGTGATCCGCGGCACTACACTGCTCTCA
TCAGAAGTCCAGAAGTTGGCCTCTGCCCTGCTCAACCAGAAGTGCCCTCTGGCCTGGCAATC
CAAGTGGGAGGGACCCGAAGACCCTCTGCAATATCTCAGAGGCCTCGTGGCTAGAGCACTGG
CCATCCAGAATTGGGTCGATAAAGCAGAGAAGCAGGCCCTCTTGTCCGAAACACTGGACCTC
TCTGAATTGTTCCATCCCGACACATTCCTGAACGCCCTCCGCCAGGAAACAGCAAGGGCTGT
TGGAAGATCAGTGGATTCTCTGAAATTTGTCGCCTCCTGGAAGGGTAGACTGCAAGAGGCCA
AACTCCAGATCAAGATCTCAGGACTGCTCCTGGAAGGCTGCTCCTTCGACGGTAATCAACTG
TCCGAGAATCAGCTCGACAGCCCAAGCGTCTCTAGCGTTCTGCCATGCTTCATGGGTTGGAT
TCCTCAAGACGCTTGCGGCCCATACTCACCCGATGAGTGCATTTCTTTGCCAGTGTACACAT
CCGCTGAGCGCGACAGAGTCGTGACCAACATCGATGTGCCTTGCGGTGGTAATCAGGATCAA
TGGATTCAGTGTGGAGCCGCCTTGTTTCTCAAGAATCAGGTCtcgatgccaggcagtccaat
gttatgtaaaacgaaggcgaaagggtgtctaacaccatctgatcgatacaaactcgcactgg
ccgcccacaaacgtcatggaaaggtagaaaattcgatgggctgacgctattaccgtatcata
ggtcgactcacgtgggtacgtgccgtagctcccatgtttaccgcttatgcgggtccagtgga
aagagctttactggacgaataacgctgctgcttttttaatccatatatgatactgcctagaac
aagtacgggaaaaatttcgacggcgtgcaatgtgcaatatgtttccgctatttgatcactctt
ggccgagtgcaatctctcactcgcgcttttgggctaacgacatagactcaatatcttagagt
gagacgtgcggtctttcagtggagaaagccctgtttagaccacaggttcctattattcggat
gaacgacctttaagataggtcaaccattatgacagttgcctgagtaagaacacgagcggagt
atctgattctgatgcttagacgctgtcgcatcccgtgaaagctcatccagaccgggtgagcg
tagacctataactacgccaacctacccggccgggaaatcatgtaggcaactcaacccgctcg
catgtaagttgtccataatatgaatttacccatccgaattgtatcgtggagttgttcggcta
gtggcaggagttcttagactaatgacaccctcactgttgcggcggtaacaaccattaactat
tacaagttgcggtttggtaaggttagggtaactgtagttaaaagtatttctgcgattgctcg
tttcgtcagatcacttcacagcgcagtctacggcactagggacaagatttgtgtactgtgga
cccgtagccgagaaatccacggcattcatgagacgttactcgggaactattcagtcagtgta
tgtagtcggcaaccggtagtggttccggaacaagcttttgaaaatcagttaatgtgggttga
gctgctcagaggcgcccagttgtaccggaaaagtatgcatatctttaatggctatgcaagta
tccaactatgctggcggcagggtagccatatccatcgggtccagaaggttcccattaaatac
ccgacactactagctatgttttgggtcctgttcccgagttacctgcataaggtgaataagcc
ttagtaagttactgctctatgacattggctcagtcagactgttgcatacttatcttatagat
tcctatgcggaaaattccgaccgcagacttaataacttgatctaatgcccggtttctcagta
cgatactagttaaaccggccagcacaggttcgtatatatccctgtatggatatgatctcggg
tcgactttaagcgactaagtgctctagcgcctgatggcgtgctttcttctcccacgttgcta
tatcaccgagtgagtacccgcggatagagttgcttcgcaggtactcaaccacagttaggcaa
gtgcgaaggtattactgattaggcgtggccgcggccgtacacctcgttagtttgagggaagc
tgttccgatgactggctagcaggcctgggtgagtactagtgatcagcaaagcctctctgtgg
tcgcataggtccgacaatatgagcgcagtatgcgagtcccaccgaaattttgcaacgatctt
gattctccctccagagtctaaatttcctatcctccaagtgtaactccagcatcagatgtttc
ttagaagtactcggagaatacaactttgagtacaatttgcgcgtccgtccggtttcccggtg
ggatttaccaacttaaaacttctagaccattattcacagcctagcgctgcgattggcggctc
atgtgcggctcggctggtagggttggtgatgtggctaaccgtcgaagcgttgtaggacacct
tatttggagaagagacgttgtgacgggaaaagtgtctcatgcgtaatgttccggagtatggg
ttcctgtgacgtgtagcagccttcaaatggacggatttacgccaatatttcattaagtgagc

| | |
|---|---|
| | ctaaaattagcccgcaacgaggtgatctgggagactacaatctaaggttgcgagtgtgatca<br>gctcagaaacaatagctcgtgaacctcgattcagcagggattgcactctgtggtccgtttct<br>ccactctctggttggtttacgagtgagccctttaaccctggtaggatccataaaacatgtaa<br>aaccaccttgcttgtgctagcattagggaccggtgttcacacccatctatatggaggctcct<br>ggtgcacctcggacaaatgaggctttaagctatcgattccgaatggctacctggcgcatgca<br>aggtgccctgttcatgtccagtcagtgctgatgggaccgctgcgagagacctagtagattct<br>tcgcataaaaatatatgtgttaacacggtactgtccgtggtatgccgttagaaataggtgcc<br>catccgactcacgtcctgccagcgaccaatggaatcccgctaaacagaaatgattatagtga<br>gcaaggcatagtaaatatcagtccggtactactctacttcgagtccctaaagaacgacaagg<br>atctgtgaataaaactatagatcaattcagtgcttaccctcctatctgcttaaggtgtaatt<br>gtatgagggtgccgcgaccttcgtcatttcgaaaatgcccagttgacctaaaaccaggcaat<br>gtctagaaccgcatagggacttaatgacaggccgccgatccagcaatgatttattcgccgaa<br>tagtcccaagtgttcacgggagatcgttattgggaggcgggtgtatgtgtgagcgccctaca<br>cccttctatcttctcgtacggccttgttgagtatttatctgccgacttgatcaaaatagggt<br>ccgccatcgtcacattacgacttcgaaatcccgatgtgctgcttagaaattttgtaacatcg<br>gtaccaagagagtgcaaaacgaataattgctgagagcagcactgtgcaagactggaaaggtg<br>gtgaagcaatacaccatgaatcccggatcgctgacgcgccctgtagcggcgcattaagcgcg<br>gcgggtgtggtggttacgcgcagcgtgaccgctacacttgccagcgccctagcgcccgctcc<br>tttcgctttcttcccttcctttctcgccacgttcgccggctttccccgtcaagctctaaatc<br>gggggctccctttagggttccgatttagtgctttacggcacctcgaccccaaaaaacttgat<br>ttgggtgatggttcacgtagtgggccatcgccctgatagacggtttttcgccctttgacgtt<br>ggagtccacgttcttt |
| 3031-nt ssDNA sequence for composition test | aatagtggactcttgttccaaactggaacaacactcaaccctatctcgggctattcttttga<br>tttataagggattttgccgatttcggaacgggtacctacgaagagttccagcagggattcca<br>agaaatggccaatgaagattggatcacctttcgcactaagacctacttgtttgaggagtttc<br>tgatgaattggcacgaccgcctcaggaaagtggaggagcattctgtgatgactgtcaagctc<br>caatctgaggtggacaaatataagattgttatccctatcctgaagtacgtccgcggagaaca<br>cctgtcacccgatcactggctggatctgttccgcttgctgggtctgcctcgcggcacatctc<br>tggagaaactgctgttcggtgacctgctgagagttgccgataccatcgtggccaaggctgct<br>gacctgaaagatctgaactcacgcgcccagggtgaagtgaccatccgcgaagcactcaggga<br>actggatttgtggggcgtgggtgctgtgttcacactgatcgactatgaggactcccagagcc<br>gcaccatgaagctgatcaaggattggaaggacatcgtcaaccaggtgggcgacaatagatgc<br>ctcctgcagtccttgaaggactcaccatactataaaggctttgaagacaaggtcagcatctg<br>ggaaaggaaactcgccgaactggacgaatatttgcagaacctcaaccatattcagagaaagt<br>gggtttacctcgaaccaatctttggtcgcggagccctgcccaaagagcagaccagattcaac<br>agggtggatgaagatttccgcagcatcatgacagatatcaagaaggacaatcgcgtcacaac<br>cttgactacccacgcaggcattcgcaactcactgctgaccatcctggaccaattgcagagat<br>gccagcgcagcctcaacgagttcctggaggagaagcgcagcgccttccctcgcttctacttc<br>atcggagacgatgacctgctggagatcttgggccagtcaaccaatccatccgtgattcagtc<br>tcacctcaagaagctgtttgctggtatcaactctgtctgtttcgatgagaagtctaagcaca<br>ttactgcaatgaagtccttggagggagaagttgtgccattcaagaataaggtgcccttgtcc<br>aataacgtcgaaacctggctgaacgatctggccctggagatgaagaagaccctggagcagct<br>gctgaaggagtgcgtgacaaccggacgcagctctcagggagctgtggacccttctctgttcc<br>catcacagatcctgtgcttggccgaacagatcaagtttaccgaagatgtggagaacgcaatt<br>aaagatcactccctgcaccagattgagacacagctggtgaacaaattggagcagtatactaa<br>catcgacacatcttccgaggacccaggtaacacagagtccggtattctggagctgaaactga<br>aagcactgattctcgacattatccataacatcgacgtggtcaagcagctgaaccaaatccaa<br>gtgcacaccaccgaagattgggcctggaagaagcagttgaggttctacatgaagtccgacca<br>cacctgttgcgttcagatggttgacagcgagttccagtacacctatgagtaccaaggaaatg<br>ccagcaagctcgtttacactccactcactgacaagtgttacctcaccttgacacaggctatg<br>aagatgggcctgggaggcaacccatacggtccagctggcactggtaagacagagagcgttaa<br>ggcactcggaggtctgctgggcaggcaggtcctcgtgttcaactgtgatgaaggaatcgacg<br>ttaagtccatgggaagaatctttgttggcctcgttaagtgtggagcttggggttgcttcgac<br>gagttcaacaggctggaggaatctgtgctgagcgccgtctctatgcagatccagaccatcca<br>ggacgcattgaagaaccacaggaccgtctgcgagctgttgggtaaggaagtggaggtgaact<br>ccaactccggaatcttcatcacaatgaatcccgcaggtaaaggatatggaggaagacagaaa<br>ctcccagacaacctgaagcagctgttccgcccagtggctatgtcccatccagacaatgagct<br>gatcgccgaagtcatcctctattccgagggattcaaagatgctaaagttctctccagaaagc |

| | |
|---|---|
| | tcgtggccatcttcaatctgtcaagagaactcctgacacctcagcagcattacgactggggt<br>ctgagagccctcaagaccgtcctgagaggttcaggaaatctcctcaggcagctgaacaagag<br>cggtacaacacagaatgcaaatgagagccacattgtcgtccaggctctgaggctgaatacca<br>tgtcaaagttcacattcacagactgcacaagatttgacgctctgattaaagatgtgttccct<br>ggtattgaactcaaagaagtggagtatgacgagctgagcgccgctttgaagcaggtgtttga<br>ggaggctaactatgagattatccctaatcagatcaagaaagcattggaactgtatgaacagc<br>tgtgtcagaggatgggagtggtgattgtgggcccatcaggcgcaggtaagagcactctctgg<br>agaatgctgagagcagcactgtgcaagactggaaaggtggtgaagcaatacaccatgaatcc<br>cggatcgctgacgcgccctgtagcggcgcattaagcgcggcgggtgtggtggttacgcgcag<br>cgtgaccgctacacttgccagcgccctagcgcccgctcctttcgctttcttcccttcctttc<br>tcgccacgttcgccggctttccccgtcaagctctaaatcgggggctccctttagggttccga<br>tttagtgctttacggcacctcgaccccaaaaaacttgatttgggtgatggttcacgtagtgg<br>gccatcgccctgatagacggtttttcgccctttgacgttggagtccacgttcttt |
| 3384-nt BCMA CAR template | aatagtggactcttgttccaaactggaacaacactcaaccctatctcggctattcttttga<br>tttataagggattttgccgatttcggaacgggtacctacgaagagttccCTTACATCTAGTT<br>GAGCTGTCACAGAATGTGACGTTGAAGgatgtaaggagctgctgtgacttgctcaaggcctt<br>atatcgagtaaacggtagtgctggggcttagacgcaggtgttctgatttatagttcaAAACC<br>TCTATCAATGAGAGAGCAATCTCCTGGTAATGTGATAGATTTCCCAACTTAATGCCAACATA<br>CCATAAACCTCCCATTCTGCTAATGCCCAGCCTAAGTTGGGGAGACCACTCCAGATTCCAAG<br>ATGTACAGTTTGCTTTGCTGGGCCTTTTTCCCATGCCTGCCTTTACTCTGCCAGAGTTATAT<br>TGCTGGGGTTTTGAAGAAGATCCTATTAAATAAAAGAATAAGCAGTATTATTAAGTAGCCCT<br>GCATTTCAGGTTTCCTTGAGTGGCAGGCCAGGCCTGGCCGTGAACGTTCACTGAAATCATGG<br>CCTCTTGGCCAAGATTGATAGCTTGTGCCTGTCCCTGAGTCCCAGTCCATCACGAGCAGCTG<br>GTTTCTAAGATGCTATTTCCCGTATAAAGCATGAGACCGTGACTTGCCAGCCCCACAGAGCC<br>CCGCCCTTGTGCATCACTGGCATCTGGACTCCAGCCTGGGTTGGGGCAAAGAGCGAAATGAG<br>ATCATGTCCTAACCCTGgaattggATCCTCTTGTCttACAGATGGATCTGGAGCAACAAACT<br>TCTCACTACTCAAACAAGCAGGTGACGTGGAGGAGAATCCCGGCCCCATGAAATGGAAAGCA<br>CTCTTTACCGCCGCAATCCTTCAAGCACAGTTGCCAATTACCGAGGCTgaacagaagcttat<br>ctctgaagaggatcttGACATTGTACTGACGCAAAGTCCCCCTAGCTTGGCGATGAGTCTCG<br>GGAAGCGAGCGACGATTAGTTGCCGAGCTTCTGAAAGTGTCACAATCCTTGGCTCCCACCTG<br>ATCCATTGGTACCAACAAAAACCTGGGCAGCCCCCGACGCTTCTCATTCAGTTGGCGTCTAA<br>CGTGCAAACAGGAGTACCGGCCAGATTTTCAGGCTCAGGCTCTCGCACCGACTTTACTCTGA<br>CCATCGACCCTGTTGAGGAGGATGATGTAGCAGTTTACTACTGTCTTCAGAGCAGAACCATT<br>CCTCGCACATTCGGCGGTGGAACGAAGTTGGAAATCAAGGGCTCAACAAGTGGGAGTGGGAA<br>GCCCGGCAGCGGGGAGGGTTCTACTAAAGGCCAAATACAGTTGGTTCAATCCGGGCCTGAAC<br>TGAAAAAGCCGGGAGAGACCGTGAAAATTTCTTGCAAGGCTAGCGGGTACACTTTTACGGAT<br>TACTCTATTAACTGGGTTAAGAGGGCACCGGGCAAAGGGCTGAAATGGATGGGCTGGATAAA<br>CACCGAGACTCGGGAGCCTGCATATGCTTATGATTTCAGAGGTAGATTTGCGTTCTCTTTGG<br>AAACCTCAGCTTCAACGGCCTACTTGCAGATTAATAACTTGAAGTACGAAGACACCGCCACT<br>TACTTCTGCGCTCTCGATTACTCATACGCTATGGATTACTGGGGCCAGGGCACGTCCGTGAC<br>CGTGTCCAGCgcaattgaagttatgtatcctcctccttacctagacaatgagaagagcaatg<br>gaaccattatccatgtgaaagggaaacacctttgtccaagtcccctatttcccggaccttct<br>aagcccttttgggtgctggtggtggttggtggagtcctggcttgctatagcttgctagtaac<br>agtggcctttattattttctgggtgaggagtaagaggagcaggctcctgcacagtgactaca<br>tgaacatgactccccgccgccccgggcccacccgcaagcattaccagccctatgccccacca<br>cgcgacttcgcagcctatcgctccagagtgaagttcagcaggagcgcagacgcccccgcgta<br>ccagcagggccagaaccagctctataacgagctcaatctaggacgaagagaggagtacgatg<br>ttttggacaagagacgtggccgggaccctgagatggggggaaagccgagaaggaagaaccct<br>caggaaggcctgtacaatgaactgcagaaagataagatggcggaggcctacagtgagattgg<br>gatgaaaggcgagcgccggaggggcaaggggcacgatggcctttaccagggtctcagtacag<br>ccaccaaggacacctacgacgcccttcacatgcaggccctgccccctcgcGGAAGCGGAGCT<br>ACTAACTTCAGCCTGCTGAAGCAGGCTGGAGACGTGGAGGAGAACCCTGGACCCaaTATCCA<br>GAACCCTGACCCTGCCGTGTACCAGCTGAGAGACTCTAAATCCAGTGACAAGTCTGTCTGCC<br>TATTCACCGATTTTGATTCTCAAACAAATGTGTCACAAAGTAAGGATTCTGATGTGTATATC<br>ACAGACAAAACTGTGCTAGACATGAGGTCTATGGACTTCAAGAGCAACAGTGCTGTGGCCTG<br>GAGCAACAAATCTGACTTTGCATGTGCAAACGCCTTCAACAACAGCATTATTCCAGAAGACA<br>CCTTCTTCCCCAGCCCAGGTAAGGGCAGCTTTGGTGCCTTCGCAGGCTGTTTCCTTGCTTCA<br>GGAATGGCCAGGTTCTGCCCAGAGCTCTGGTCAATGATGTCTAAAACTCCTCTGATTGGTGG |

| | |
|---|---|
| | TCTCGGCCTTATCCATTGCCACCAAAACCCTCTTTTTACTAAGAAACAGTGAGCCTTGTTCT<br>GGCAGTCCAGAGAATGACACGGGAAAAAAGCAGATGAAGAGAAGGTGGCAGGAGAGGGCACG<br>TGGCCCAGCCTCAGTCTCTCCAACTGAGTTCCTGCCTGCCTGCCTTTGCTCAGACTGTTTGC<br>CCCTTACTGCTCTTCTAGGCCTCATTCTAAGCCCCTTCTCCAAGTTGACGTTGAAGCGTTAC<br>CTGTTAGGTAACGTAGTTGAGCTGTCAACTTGGccatgaatcccggatcgctgacgcgccct<br>gtagcggcgcattaagcgcggcgggtgtggtggttacgcgcagcgtgaccgctacacttgcc<br>agcgccctagcgcccgctcctttcgctttcttcccttcctttctcgccacgttcgccggctt<br>tccccgtcaagctctaaatcgggggctccctttagggttccgatttagtgctttacggcacc<br>tcgaccccaaaaaacttgatttgggtgatggttcacgtagtgggccatcgccctgatagacg<br>gtttttcgccctttgacgttggagtccacgttcttt |
| 3396-nt BCMA CAR template | aatagtggactcttgttccaaactggaacaacactcaaccctatctcggctattcttttga<br>tttataagggattttgccgatttcggaacgggtacctacgaagagttccCTTACATCTAGTT<br>GAGCTGTCACAGAATGTGACGTTGAAGGATGTAAGGAGCTGCTGTGACTTGCTCAAGGCCTT<br>ATATCGAGTAAACGGTAGTGCTGGGGCTTAGACGCAGGTGTTCTGATTTATAGTTCAAAACC<br>TCTATCAATGAGAGAGCAATCTCCTGGTAATGTGATAGATTTCCCAACTTAATGCCAACATA<br>CCATAAACCTCCCATTCTGCTAATGCCCAGCCTAAGTTGGGGAGACCACTCCAGATTCCAAG<br>ATGTACAGTTTGCTTTGCTGGGCCTTTTTCCCATGCCTGCCTTTACTCTGCCAGAGTTATAT<br>TGCTGGGGTTTTGAAGAAGATCCTATTAAATAAAAGAATAAGCAGTATTATTAAGTAGCCCT<br>GCATTTCAGGTTTCCTTGAGTGGCAGGCCAGGCCTGGCCGTGAACGTTCACTGAAATCATGG<br>CCTCTTGGCCAAGATTGATAGCTTGTGCCTGTCCCTGAGTCCCAGTCCATCACGAGCAGCTG<br>GTTTCTAAGATGCTATTTCCCGTATAAAGCATGAGACCGTGACTTGCCAGCCCCACAGAGCC<br>CCGCCCTTGTCCATCACTGGCATCTGGACTCCAGCCTGGGTTGGGGCAAAGAGGGAAATGAG<br>ATCATGTCCTAACCCTGgaattggATCCTCTTGTCttACAGATGGATCTGGAGCAACAAACT<br>TCTCACTACTCAAACAAGCAGGTGACGTGGAGGAGAATCCCGGCCCcatggcacttccagta<br>actgcgctgctgctcccgctcgcactcctgctgcatgcggcccgaccagaacagaagcttat<br>ctctgaagaggatcttGACATTGTACTGACGCAAAGTCCCCCTAGCTTGGCGATGAGTCTCG<br>GGAAGCGAGCGACGATTAGTTGCCGAGCTTCTGAAAGTGTCACAATCCTTGGCTCCCACCTG<br>ATCCATTGGTACCAACAAAAACCTGGGCAGCCCCCGACGCTTCTCATTCAGTTGGCGTCTAA<br>CGTGCAAACAGGAGTACCGGCCAGATTTTCAGGCTCAGGCTCTCGCACCGACTTTACTCTGA<br>CCATCGACCCTGTTGAGGAGGATGATGTAGCAGTTTACTACTGTCTTCAGAGCAGAACCATT<br>CCTCGCACATTCGGCGGTGGAACGAAGTTGGAAATCAAGGGCTCAACAAGTGGGAGTGGGAA<br>GCCCGGCAGCGGGGAGGGTTCTACTAAAGGCCAAATACAGTTGGTTCAATCCGGGCCTGAAC<br>TGAAAAAGCCGGGAGAGACCGTGAAAATTTCTTGCAAGGCTAGCGGGTACACTTTTACGGAT<br>TACTCTATTAACTGGGTTAAGAGGGCACCGGGCAAAGGGCTGAAATGGATGGGCTGGATAAA<br>CACCGAGACTCGGGAGCCTGCATATGCTTATGATTTCAGAGGTAGATTTGCGTTCTCTTTGG<br>AAACCTCAGCTTCAACGGCCTACTTGCAGATTAATAACTTGAAGTACGAAGACACCGCCACT<br>TACTTCTGCGCTCTCGATTACTCATACGCTATGGATTACTGGGGCCAGGGCACGTCCGTGAC<br>CGTGTCCAGCgcaaccacgacgccagcgccgcgaccaccaacaccggcgcccaccatcgcgt<br>cgcagccTctgtccctgcgcccagaggcgtgccgAccagcggcgggTggAgcagtgcacacg<br>agggggctggacttcgcctgtgatatctacatctgggcgcccttggccgggacttgtggggt<br>ccttctcctgtcactggttatcaccctttaCTGCAAGCGGGGCAGAAAGAAGCTGCTGTACA<br>TCTTCAAGCAGCCCTTCATGCGGCCCGTGCAGACCACCCAGGAAGAGGACGGCTGCTCCTGC<br>AGATTCCCCGAGGAAGAAGAAGGCGGCTGCGAGCTGagagtgaagttcagcaggagcgcaga<br>cgcccccgcgtaccagcagggccagaaccagctctataacgagctcaatctaggacgaagag<br>aggagtacgatgttttggacaagagGcgtggccgggaccctgagatggggggaaagccgaga<br>aggaagaaccctcaggaaggcctgtacaatgaactgcagaaagataagatggcggaggccta<br>cagtgagattgggatgaaaggcgagcgccggaggggcaaggggcacgatggcctttaccagg<br>gtctcagtacagccaccaaggacacctacgacgcccttcacatgcaggccctgccccctcgc<br>GGAAGCGGAGCTACTAACTTCAGCCTGCTGAAGCAGGCTGGAGACGTGGAGGAGAACCCTGG<br>ACCCaaTATCCAGAACCCTGACCCTGCCGTGTACCAGCTGAGAGACTCTAAATCCAGTGACA<br>AGTCTGTCTGCCTATTCACCGATTTTGATTCTCAAACAAATGTGTCACAAAGTAAGGATTCT<br>GATGTGTATATCACAGACAAAACTGTGCTAGACATGAGGTCTATGGACTTCAAGAGCAACAG<br>TGCTGTGGCCTGGAGCAACAAATCTGACTTTGCATGTGCAAACGCCTTCAACAACAGCATTA<br>TTCCAGAAGACACCTTCTTCCCCAGCCCAGGTAAGGGCAGCTTTGGTGCCTTCGCAGGCTGT<br>TTCCTTGCTTCAGGAATGGCCAGGTTCTGCCCAGAGCTCTGGTCAATGATGTCTAAAACTCC<br>TCTGATTGGTGGTCTCGGCCTTATCCATTGCCACCAAAACCCTCTTTTTACTAAGAAACAGT<br>GAGCCTTGTTCTGGCAGTCCAGAGAATGACACGGGAAAAAAGCAGATGAAGAGAAGGTGGCA<br>GGAGAGGGCACGTGGCCCAGCCTCAGTCTCTCCAACTGAGTTCCTGCCTGCCTGCCTTTGCT |

| | |
|---|---|
| | CAGACTGTTTGCCCCTTACTGCTCTTCTAGGCCTCATTCTAAGCCCCTTCTCCAAGTTGACG<br>TTGAAGCGTTACCTGTTAGGTAACGTAGTTGAGCTGTCAACTTGGccatgaatcccggatcg<br>ctgacgcgccctgtagcggcgcattaagcgcggcgggtgtggtggttacgcgcagcgtgacc<br>gctacacttgccagcgccctagcgcccgctcctttcgctttcttcccttcctttctcgccac<br>gttcgccggctttccccgtcaagctctaaatcgggggctccctttagggttccgatttagtg<br>ctttacggcacctcgaccccaaaaaacttgatttgggtgatggttcacgtagtgggccatcg<br>ccctgatagacggtttttcgccctttgacgttggagtccacgttcttt |
| 3024-nt scaffold for DNA origami tile | aatagtggactcttgttccaaactggaacaacactcaaccctatctcgggctattcttttga<br>tttataagggattttgccgatttcggaacggattactctacttcctcaacaaaccacattac<br>catcatcttccatcacttatcaaagtctttatttatccactcctctaccttactaacctctt<br>ctatccatataactctctggcttaaatacctcctatacaaattactcctacctctaataatt<br>aaaaaatttattgcacacaaataacctctcctaaactcaataatcaaccttacctcctcacc<br>aaataacgaactcctaactactatttcaaaaaacctcttacactcaatatctttaaataaat<br>agtactattcaaacatctccaactacttcacttattacacaattcctcaacactcctaattc<br>aaattacaaacacaccaattacgtaccttatcctttatcaaatttctcggaaactaccacct<br>ctcaaccttcatacattaccaccaatacttcattaatatatatcgttattccattattttta<br>aacaaaccgacaggattggaaggacatcgtcaaccaggtgggcgacaatagatgcctcctgc<br>agtccttgaaggactcaccatactataaaggctttgaagacaaggtcagcatctgggaaagg<br>aaactcgccgaactggacgaatatttgcagaacctcaaccatattcagagaaagtgggttta<br>cctcgaaccaatctttggtcgcggagccctgcccaaagagcagaccagattcaacagggtgg<br>atgaagatttccgcagcatcatgacagatatcaagaaggacaatcgcgtcacaaccttgact<br>acccacgcaggcattcgcaactcactgctgaccatcctggaccaattgcagagatgccagcg<br>cagcctcaacgagttcctggaggagaagcgcagcgccttccctcgcttctacttcatcggag<br>acgatgacctgctggagatcttgggccagtcaaccaatccatccgtgattcagtctcacctc<br>aagaagctgtttgctggtatcaactctgtctgtttcgatgagaagtctaagcacattactgc<br>aatgaagtccttggagggagaagttgtgccattcaagaataaggtgcccttgtccaataacg<br>tcgaaacctggctgaacgatctggccctggagatgaagaagaccctggagcagctgctgaag<br>gagtgcgtgacaaccggacgcagctctcagggagctgtggacccttctctgttcccatcaca<br>gatcctgtgcttggccgaacagatcaagtttaccgaagatgtggagaacgcaattaaagatc<br>actccctgcaccagattgagacacagctggtgaacaaattggagcagtatactaacatcgac<br>acatcttccgaggacccaggtaacacagagtccggtattctggagctgaaactgaaagcact<br>gattctcgacattatccataacatcgacgtggtcaagcagctgaaccaaatccaagtgcaca<br>ccaccgaagattgggcctggaagaagcagttgaggttctacatgaagtccgaccacacctgt<br>tgcgttcagatggttgacagcgagttccagtacacctatgagtaccaaggaaatgccagcaa<br>gctcgtttacactccactcactgacaagtgttacctcaccttgacacaggctatgaagatgg<br>gcctgggaggcaacccatacggtccagctggcactggtaagacagagagcgttaaggcactc<br>ggaggtctgctgggcaggcaggtcctcgtgttcaactgtgatgaaggaatcgacgttaagtc<br>catgggaagaatctttgttggcctcgttaagtgtggagcttggggttgcttcgacgagttca<br>acaggctggaggaatctgtgctgagcgccgtctctatgcagatccagaccatccaggacgca<br>ttgaagaaccacaggaccgtctgcgagctgttgggtaaggaagtggaggtgaactccaactc<br>cggaatcttcatcacaatgaatcccgcaggtaaaggatatggaggaagacagaaactcccag<br>acaacctgaagcagctgttccgcccagtggctatgtcccatccagacaatgagctgatcgcc<br>gaagtcatcctctattccgagggattcaaagatgctaaagttctctccagaaagctcgtggc<br>catcttcaatctgtcaagagaactcctgacacctcagcagcattacgactggggtctgagag<br>ccctcaagaccgtcctgagaggttcaggaaatctcctcaggcagctgaacaagagcggtaca<br>acacagaatgcaaatgagagccacattgtcgtccaggctctgaggctgaataccatgtcaaa<br>gttcacattcacagactgcacaagatttgacgctctgattaaagatgtgttccctggtattg<br>aactcaaagaagtggagtatgacgagctgagcgccgctttgaagcaggtgtttgaggaggct<br>aactatgagattatccctaatcagatcaagaaagcattggaactgtatgaacagctgtgtca<br>gaggatgggagtggtgattgtgggcccatcaggcgcaggtaagagcactctctggagaatgc<br>tgagagcagcactgtgcaagactggaaaggtggtgaagcaatacaccatgaatcccggatcg<br>ctgacgcgccctgtagcggcgcattaagcgcggcgggtgtggtggttacgcgcagcgtgacc<br>gctacacttgccagcgccctagcgcccgctcctttcgctttcttcccttcctttctcgccac<br>gttcgccggctttccccgtcaagctctaaatcgggggctccctttagggttccgatttagtg<br>ctttacggcacctcgaccccaaaaaacttgatttgggtgatggttcacgtagtgggccatcg<br>ccctgatagacggtttttcgccctttgacgttggagtccacgttcttt |

| | |
|---|---|
| Staple for DNA origami tile | GGCATCTATTGTCGCCCACCTTCTGGTCAAAATGTATGGTGGGAGCCCCTTGCTTC |
| Staple for DNA origami tile | TGCTCTTGTCATGATGCTGCGCCCAAGAACAGTAAAGGTTCACCTTTCCCTCAGCT |
| Staple for DNA origami tile | AGGTTTTGGGGTCGAGGGAGTGTTGTTCCAGTTTGGAATCAAGTTTAAACCC |
| Staple for DNA origami tile | AATAAACCATCGGAAGAAAGCGAAACCACACCGGAACTCTTGCAGTAATGTGCCGTTATT |
| Staple for DNA origami tile | TGTGACGCGATTGTCCTTCTTTCGTCTCGGATTCACACAGTGCTGCTCTAACACCT |
| Staple for DNA origami tile | TGGGTAGTCAAGGTCGAGGGAACAGGGCGCGTCAG |
| Staple for DNA origami tile | TCTCCAGTGAGGTGAGACTGACCTTCAGCGGGAGTAGGTATCATACTCCCTCTCAG |
| Staple for DNA origami tile | GGCACCTGACTTCAGTTGAGGGGTCAGCGGTTCTGCCCAGATGCTGACCTTGTCTTCTG |
| Staple for DNA origami tile | CAGCTGCCACAGCTCCCTGAGAATACCGGATATTGTTGATCGGTCTTGA |
| Staple for DNA origami tile | CATCTCCAGGGCCAACAGAGTTGATACCAGCAAACTCTTCTTGCTTCAATGAGTTCAATA |
| Staple for DNA origami tile | GTTTCGATTAGACTGCGCTTCGCCTGCGACTTTCTCAAAGCCTTTATAGTATGGTGTTCG |
| Staple for DNA origami tile | GACTCTGATAATGTCGAGAATGGAGTGT |
| Staple for DNA origami tile | ACCAGCTGCCCAATCTTCGGTACCATCTACTTCGGATTCCGGAGTTGGAAATGCGT |
| Staple for DNA origami tile | GTTATGGTGTTACCAAGGGTCTCCAGGGAGCTTCTCAGGTCAGATATCTTGGGCAG |
| Staple for DNA origami tile | CAGCTGCTTGACCACGTCGATGCATTTC |

| | |
|---|---|
| Staple for DNA origami tile | CTCCCAGGCCCACTTAACATTCCTCCAGCCTGTTTCTGTACAGCTGGGGCTCTCCTGAAC |
| Staple for DNA origami tile | ATTGTGTAATAAGTGTGAGGTAGTGAGTCAGTGCTGCTCCAGAGCTGCGACGCACT |
| Staple for DNA origami tile | TTCTTCCCATGGACTTAACGTTTACCAGGCTGTCAGGTGTGCGCTCCAATGATCTG |
| Staple for DNA origami tile | CGATTCCCCTGGATGGTCTGGGATGAAGCGATCAGGTTCTCTTTCTGTGGCGTCAA |
| Staple for DNA origami tile | TTCATCACAGTTGAAGTGCCTACAGGTGCTTCCAGGTGTCTCTCTCCAC |
| Staple for DNA origami tile | TCCTGTGGTTCTTCGTTCACCCGGAATATGGCCACACAATGTGTGAATGCAATGCTCACA |
| Staple for DNA origami tile | ATCTGCATAGAGACGGCGCTCTTTACCTGCGGGATTCATTGT |
| Staple for DNA origami tile | ATTGAGTTTTTGAAGGTAAGGAGTTTAGTTATTAGAATTTGTATAGAAGGGTAGAG |
| Staple for DNA origami tile | TTAGGTTGTAAGGTCTGGGAGTTGAACTACCCCAAGCTCCACATCTTCACTTGCTG |
| Staple for DNA origami tile | CTTCAGGAAACGAGTAGCCTGTGTCAAGGAAAGCACGTCGAGTAGTTGG |
| Staple for DNA origami tile | GGGCGGACTTCCTCCATATCCAGCACAGGAGGCCAACAAAGATATGGGTTGC |
| Staple for DNA origami tile | AATGCTGCTGAGGTGTCAGGACTCATTGGGAACTCTGCCAGCTGGACCG |
| Staple for DNA origami tile | TGACAGATTGAAGAGAGGATGGAACGCATAACGCTCTCTGTC |
| Staple for DNA origami tile | GAGATTTCAGACCCCAGTCGTAGCCACTCTTGGTACTCATAG |
| Staple for DNA origami tile | ACTTCTTAGCGGCGAGTCTTGTGGTGTACGATTTAAAATCGGAGAATAGCCCGAGATAGG |

| | |
|---|---|
| Staple for DNA origami tile | CCAGCCTGAGTCGGAAGATGTGTCGGATCTGTGATGGGAACAGAGTGGGTCC |
| Staple for DNA origami tile | TTTAATCAGATTGTACCGTATACTACTTGGATTTGGTTGTGTACTTCTGGATGGGACAT |
| Staple for DNA origami tile | TTCTTGAGGACAAGATCTTCGGTAAACTTTTGTTCATTTGCA |
| Staple for DNA origami tile | AGTTAGCCTCCTCACAGCATTCTCCAGAGAGTGCT |
| Staple for DNA origami tile | ATCTCATCAGCCAGTTCGGCCAAGCACAGATGTTAGCTCTTGTTCAGCTGGGAACACATC |
| Staple for DNA origami tile | AGGGATACTTACCTCTTAATGAAGGAAGACCCAAACAAGAGTCCACTATTAAAGAA |
| Staple for DNA origami tile | TCTGATTATCTTGTGCAGTCTGGCTCTC |
| Staple for DNA origami tile | GAAGATGGGTTTGTCAAAATCCCTTATAAATCAAAAACCCTA |
| Staple for DNA origami tile | AAGCCGGCGAACGTGGCGAGACGCCGCTAGGCGCTTCTCATCGAAACAGGATCGTT |
| Staple for DNA origami tile | CTACGTGTGGTTGAAGTGAGTTGCGAATTCCTCCACGCCGCGGCGCCTGATGGGCC |
| Staple for DNA origami tile | AAGCACTGAGCTTGACGGGGACGATCCGCGATGAAGTAGAAG |
| Staple for DNA origami tile | GTTTGCCGTACGACCAAAGATTGGAGTCCTTCAAGGACTGCAGGAGGCTCCG |
| Staple for DNA origami tile | TTTTCGGCGAGTTTCCTTTCAAATATAGGGCGAGCGCTAGGGCGCTGGCAAGTTT |
| Staple for DNA origami tile | TTTGCAATTGGTCCAGGATCTGCGCTGCTGCGCACCACTCCCATCCTCTGACTTT |
| Staple for DNA origami tile | TTTAACTTCTCCCTCCAAGTATTCTTACAGTTCTGAACTTTGACATGGTATTTTT |

| | |
|---|---|
| Staple for DNA origami tile | TTTTGATCTTTAATTGCGTAATCTGGCTGGACGGAGCTTTCTGGAGAGAACTTTT |
| Staple for DNA origami tile | TTTAGAACCTCAACTGCTTTGGTCGGAATCCCTTCCACTTCCTTACCCAACATTT |
| Staple for DNA origami tile | TTTGCCCAGCAGACCTCCGACACGAG |
| Staple for DNA origami tile | TTTGCTCGCAGACGGGACCTGCCTTTT |
| Staple for DNA origami tile | TTTTTAGCATCTTTGACTTCATGTTTT |
| Staple for DNA origami tile | TTTCAGCCTCAGAGCTGCAGGGAGTTT |
| Staple for DNA origami tile | TTTACAGCTGTTCATGAATGGCACTTT |
| Staple for DNA origami tile | ATCGTAACCAGGAGCGGTGGCCCACGTGGACTCCAACGTCAAAGGGCGAATTT |
| Staple for DNA origami tile | TTTTGTAGCGGTCACGGCATCTCTTTT |
| Staple for DNA origami tile | TTTAAACCGTCTATCTCGTCCAGTTTT |
| Staple for DNA origami tile | TTTCGGTTTGTTTAAGTAGAGTAATTT |
| Staple for DNA origami tile | TTTATATATATTAATTAAATAAAGTTT |
| Staple for DNA origami tile | ATCACGGTGACTGGGAAATCTTGTTGAAGGTTGACGATGTCCTTCCAATCCTGTTTT |
| Staple for DNA origami tile | TTTTGTATGAAGGTTGTATTTAAGTTT |
| Staple for DNA origami tile | TTTCGAGAAATTTGATATTTGTGTTTT |

| | |
|---|---|
| Staple for DNA origami tile | TTTTAATTGGTGTGTTTAGGAGTTTTT |
| Staple for DNA origami tile | TTTGAGTGTTGAGGATGAATAGTATTT |
| Staple for DNA origami tile | TTTCTATTTATTTAAAGATAGATGTT |
| Staple for DNA origami tile | TTTCGTTATTTGGTGAGGAATAGTAGTTTTGTCAACACGTAATTTGAATTAGTTT |
| Staple for DNA origami tile | TTTGCAATAAATTTTTTAAGAGAGGTTATTTCATTCAGAAGGATAAGGTACGTTT |
| Staple for DNA origami tile | TTTCCAGAGAGTTATATGGATAGGAGGATTGTCTCCGGGAGGTGGTAGTTTCTTT |
| Staple for DNA origami tile | TTTACTTTGATAAGTGATGGAGTGGAGATTGGTATGGAAGTATTGGTGGTAATTT |
| Staple for DNA origami tile | TTTTCCGTTCCGAAATCGGTGAGGAAAACACCCTCATCATAATGGAATAACGTTT |
